# A Functional Assay For Mining Non-Inhibitory Enzyme Ligands From One Bead One Compound Libraries: Application to E3 Ubiquitin Ligases

**DOI:** 10.1101/2025.04.29.651224

**Authors:** Weijun Gui, Allison Goss, Thomas Kodadek

## Abstract

Chemical dimerizers are synthetic molecules that bring into proximity two or more proteins that do not normally interact with one another. A major application of this technology is to recruit an enzyme to a target protein, resulting in its post-translational modification (PTM). In particular, chemical dimerizer-mediated poly-Ubiquitylation of proteins has garnered an enormous amount of interest as a new drug modality. A fundamental requirement for the construction of new PTM-driving dimerizers is an enzyme ligand that does not inhibit its activity. Traditional activity-based high-throughput screening platforms are not suited for this purpose. Here we describe a novel platform for screening libraries of bead-displayed compounds that links a requirement for small molecule binding to the enzyme with enzyme-mediated modification of a nearby substrate. This system ensures that the enzyme-recruiting small molecules do not interfere with the catalytic function of the enzyme. We demonstrate the utility of this system in the context of E3 Ubiquitin ligase-recruiting molecules and report the discovery of a novel, low molecular mass ligand for the Von Hippel Landau (VHL) protein.

## Introduction

Chemical dimerizers are of increasing utility in chemical biology and drug development.^1^ These are synthetic molecules capable of bringing two or more proteins into proximity that are not associated in the absence of the dimerizer. A particularly important application of this concept is to bring a target protein into proximity with an enzyme capable of mediating some type of post-translation modification (PTM) of the target that modifies its activity and/or level. Prominent examples include proteolysis-targeting chimeras (PROTACs)^2, 3^ and molecular glue degraders ^4-6^ that recruit an E3 Ubiquitin ligase complex to a “neo-substrate”, resulting in its poly-Ubiquitylation and subsequent destruction by the proteasome. Dimerizers that mediate other chemical modifications of target proteins, such as phosphorylation and de-Ubiquitylation, are also being explored.^7-10^ In all such applications, it is critical that the dimerizer bind the modifying enzyme without inhibiting its activity. Unfortunately, the vast majority of small molecules that engage enzymes, including proteases, phosphatases, kinases, ligases, etc. are antagonists. These are most commonly identified through activity-based high-throughput screening (HTS) campaigns, which can register a small molecule-dependent decrease (or increase) in enzyme activity but are unable to distinguish innocent ligands from non-binders.

In contrast, other screening formats are designed to simply register ligand-protein binding and are agnostic to the effect of the ligand on target protein activity. For example, there is a long history of creating libraries by split and pool solid phase synthesis and then screening the resultant one bead one compound (OBOC) library for compounds able to engage a target protein.^11^ This can be done on-resin or after release of the compounds into solution and arraying them on a suitable surface, such as a glass slide.^12, 13^

Over the last several years DNA-encoded library (DEL) technology^14^ has emerged as a powerful approach to ligand discovery. In the most common DEL format,^15^ millions of small molecules are created by solution phase split and pool synthesis, each tethered to a piece of DNA whose sequence reflects the structure of the attached molecule. These libraries are screened by incubation with an immobilized protein. The structures of the putative ligands are inferred by subsequent PCR amplification and deep sequencing of the protein-associated encoding tags. DEL technology has also been adapted to OBOC libraries. An elaboration of the basic screening methodology, in which screens against a given target are carried out in the presence and absence of a partner protein, has allowed the isolation of chemical dimerizers from DELs.^16^

While DELs are not without their own technical issues, it is fair to say that this is now the method of choice for protein ligand discovery. However, enzyme ligands mined from a DEL, or through any binding assay, may or may not be inhibitors. This must be determined empirically through hit re-synthesis and testing. Therefore, there remains a need for a high-throughput screening platform that couples small molecule-enzyme binding with a measure of enzyme activity, thus discouraging the identification of enzyme inhibitors as hits.

Here we report a screening platform that satisfies this criterion (Figure 1). The crux of the method is to create an OBOC library on TentaGel (TG) resin in which an enzyme substrate is co-immobilized with the small molecule, then expose the beads to soluble enzyme under conditions where ligand-independent, “in trans” modification of the bead-displayed substrate is negligible. Ligand-enzyme binding will create a high concentration of the enzyme in the proximity of the immobilized substrate, which we postulate will drive the PTM of interest.

**Figure 1:**
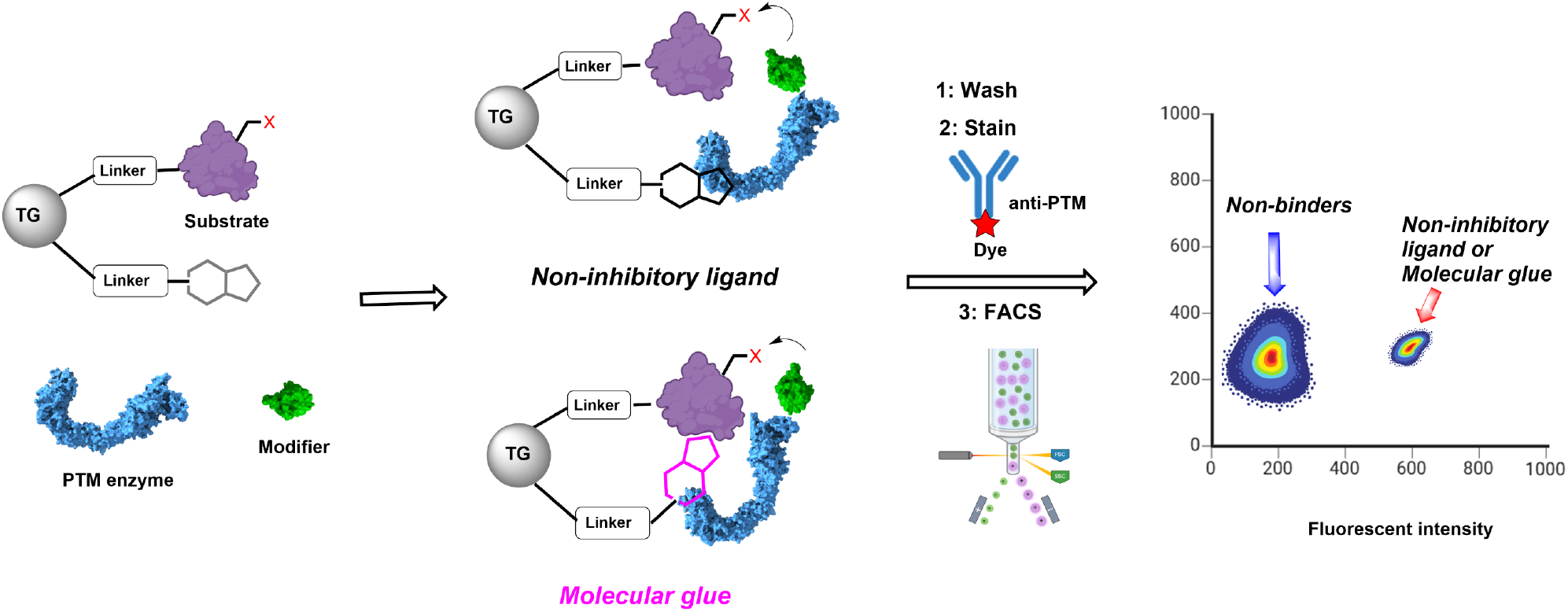
On-resin functional assay for mining non-inhibitory enzyme ligands and molecular glue from one bead one compound libraries. See text for details.

Proteins on beads that display molecules incapable of binding the enzyme, or which act as inhibitors, should not be modified. Beads displaying a modified protein substrate can then be identified by staining with a fluorescently labeled anti-PTM antibody or some other suitable detection agent. This type of screen should also recover molecular glues so long as formation of a ternary complex results in the PTM of interest. We demonstrate here the utility of this platform for E3 ubiquitin ligase ligand discovery.

## RESULTS

### Establishment of a proximity-driven, on-resin Ubiquitylation assay for the VHL ligase

Given the importance of targeted protein degradation agents, which mediate the association of a substrate protein and an activated E3 Ubiquitin ligase complex, we focused our initial efforts on the discovery of E3 ligase ligands, with the initial goal of establishing a “minimalist” system capable of detecting ligands for the von Hippel Landau (VHL) E3 Ubiquitin ligase complex. ^17, 18^ To this end, we generated six different types of 10 μm TG beads (Figure 2A), that displayed either the VHL ligand VH 032, or, as a negative control, an acetyl group, along with one, two, or three lysine residues. Synthetic details are provided in Supplementary Figure S1. These beads were incubated with purified Ube1 (50 nM; an E1 Ubiquitin ligase), UbcH5a (500 nM; an E2 Ubiquitin ligase), Neddylated VHL E3 ligase complex (Neddylated CUL2/RBX, Elongin B/Elongin C/VHL; 20 nM) as well as Ubiquitin (Ub; 20 μM) and ATP (1 mM) at 37°C for 18 hours.

**Figure 2:**
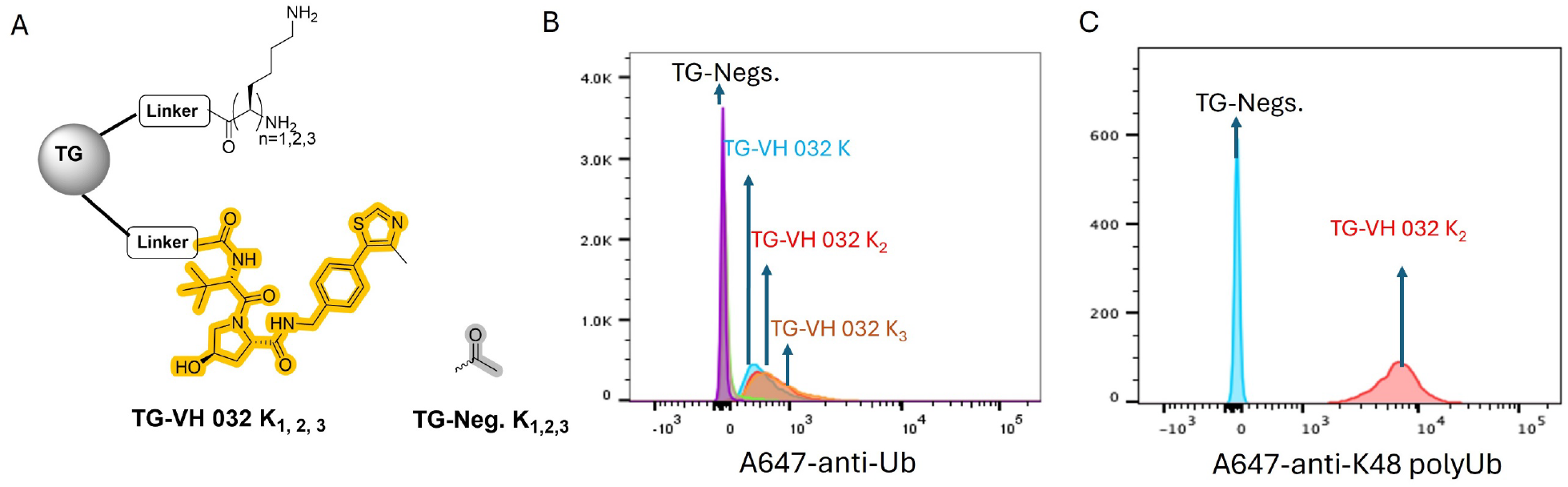
Establishment of an activity-coupled assay for VHL E3 ligase ligands using a minimalist peptide substrate. (A) Chemical structure of TG-VH 032 K_1,2,3_ and TG-Neg. K_1,2,3_. (B&C) Flow cytometry analysis of the indicated beads after incubation with Ube1 (50 nM), UbcH5a (500 nM), Neddylated CUL2/RBX, Elongin B/Elongin C/VHL (20 nM), Ubiquitin (Ub; 20 μM) and ATP (1 mM) at 37°C for 18 hours and staining with Alexa Fluor 647 labeled anti-ubiquitin antibody or anti-K48 polyubiquitin antibody.

After thorough washing, the beads were then stained with Alexafluor 647 (A647)-conjugated anti-Ubiquitin antibody. After another wash, the beads, which are comparable in size to a red blood cell, were analyzed by flow cytometry. As shown in Figure 2B, the beads displaying the VHL ligand VH 032^19^ and one, two or three lysine residues, displayed much higher Ub-dependent fluorescence than the corresponding beads displaying an acetyl unit and thus unable to engage the VHL complex. The number of lysine residues made little difference in the intensity of the signal. A similar result, but with even better separation between displaying a ligand and those that do not, was obtained when the experiment was repeated but the beads were stained with a labeled antibody that recognizes K48-linked poly-Ub chains (Figure 2C). These data demonstrate that this activity-based assay using a “minimalist substrate” (1-3 lysine residues), can readily distinguish between beads displaying a VHL ligand and those displaying a non-binder, and that the enzymatic complex recruited to the beads is capable of mediating multiple Ubiquitylation events, resulting in chain formation.

We next examined the Ubiquitylation of a protein substrate in this format. The bromodomains of the BRD proteins are often used as a paradigmatic substrate for targeted protein degradation assays, in part because of the availability of high affinity ligands such as JQ1.^20, 21^ To attach this substrate to the resin, we expressed and purified a fusion protein containing the HaloTag protein (HTP),^22^ the bromodomain 2 of BRD4, and a C-terminal His_6_ tag. Instead of placing a lysine substrate on the resin, we attached a chloroalkane, allowing covalent conjugation of the fusion protein to the bead surface, as shown in Figure 3A. Loading of HTP-BRD4bd2 onto the resin was optimized and confirmed by staining the beads with a fluorescently labeled anti-His_6_ antibody followed by flow cytometry analysis (Supplementary Figure S2). These beads were also equipped with VH032, an acetyl group, or, as additional control, a diastereomer of VH032 (VH032(−)), which has a drastically reduced affinity for VHL^23^ (Figure 3A and Supplementary Figure S3).

**Figure 3:**
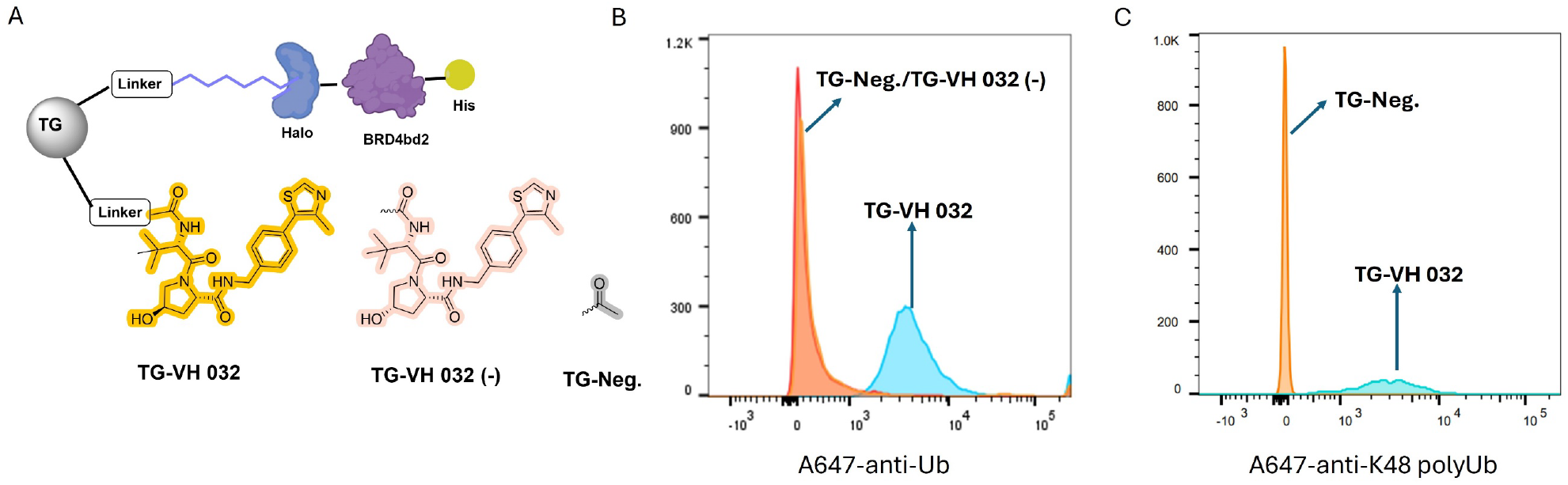
Establishment of an activity-coupled assay for VHL E3 ligase ligands using a protein substrate comprised of the HaloTag protein fused to one of the bromodomains of BRD4. (A) Chemical structure of TG-VH 032 K_1,2,3_, TG-VH032(−), an inactive diastereomer, and TG-Neg. K_1,2,3_. (B&C) Flow cytometry analysis of the indicated beads after incubation with Ube1 (50 nM), UbcH5a (500 nM), Neddylated CUL2/RBX, Elongin B/Elongin C/VHL (20 nM), Ubiquitin (Ub; 20 μM) and ATP (1 mM) at 37°C for 18 hours and staining with Alexa Fluor 647 labeled anti-ubiquitin antibody or anti-K48 polyubiquitin antibody.

The resin was incubated with the E1, E2 and VHL E3 ligase enzymes under the same conditions used for the Figure 2 experiment. As shown in Figure 3B & 3C, beads displaying VH032 displayed a much higher level of Ubiquitin-dependent fluorescence than beads displaying the acetyl unit or VH032(−), the inactive diastereomer. Little or no fluorescent signal was observed on beads displaying TG-VH 032 incubated in the presence of the E1 and E2 enzymes and ATP but in the absence of the VHL E3 ligase complex, or in the absence of Ubiquitin (Figure S4).

We also investigated the effect of varying the length of the linker connecting the ligand to the TG resin on the Ubiquitylation efficiency of beads displaying VH032 and HTP-BRD4bd2-His6. We found that linker length had almost no effect (Supplementary Figures S5 and S6).

### Comparison of VHL E3 ligands and PROTACs

These data show that when the E2/E3 Ubiquitylation complex is brought into close proximity with a substrate by virtue of binding to a resin-displayed ligand, Ubiquitin transfer occurs readily in the crowded environment of the bead. We hypothesized that substrate Ubiquitylation might be even more efficient when the resin displays a *bona fide* PROTAC or glue due to the additional free energy contributed by neo-protein-protein interactions (neo-PPIs) between the ligase and the substrate in the ternary complex. Moreover, it seems likely that such a difference will be more pronounced when the soluble enzyme is present at lower concentrations. If so, this would be an important finding with respect to optimizing conditions in high throughput screens using this assay to favor the inclusion or the exclusion of simple E3 ligands from the hit pool.

These data are shown in Figure 4 for beads displaying the VH032 ligand or the PROTAC MZ1^24^ (Figure 4A). Under the standard conditions, the ligand- and PROTAC-displaying beads supported similar levels of HTP-BRD4bd2 Ubiquitylation (Figure 4B & 4C). However, as the enzymes were diluted, the difference in signal intensity became more pronounced. For example, at a 20-fold dilution of the enzymes relative to the original conditions, the beads displaying VH032 were not distinguishable from those that were simply acetylated (Figure 4D & 4E). These data indicate that the assay stringency can readily be manipulated to favor the identification of glues/PROTACs and disfavor the isolation of simple E3 ligands, though very high affinity E3 ligands would likely still be identified.

**Figure 4:**
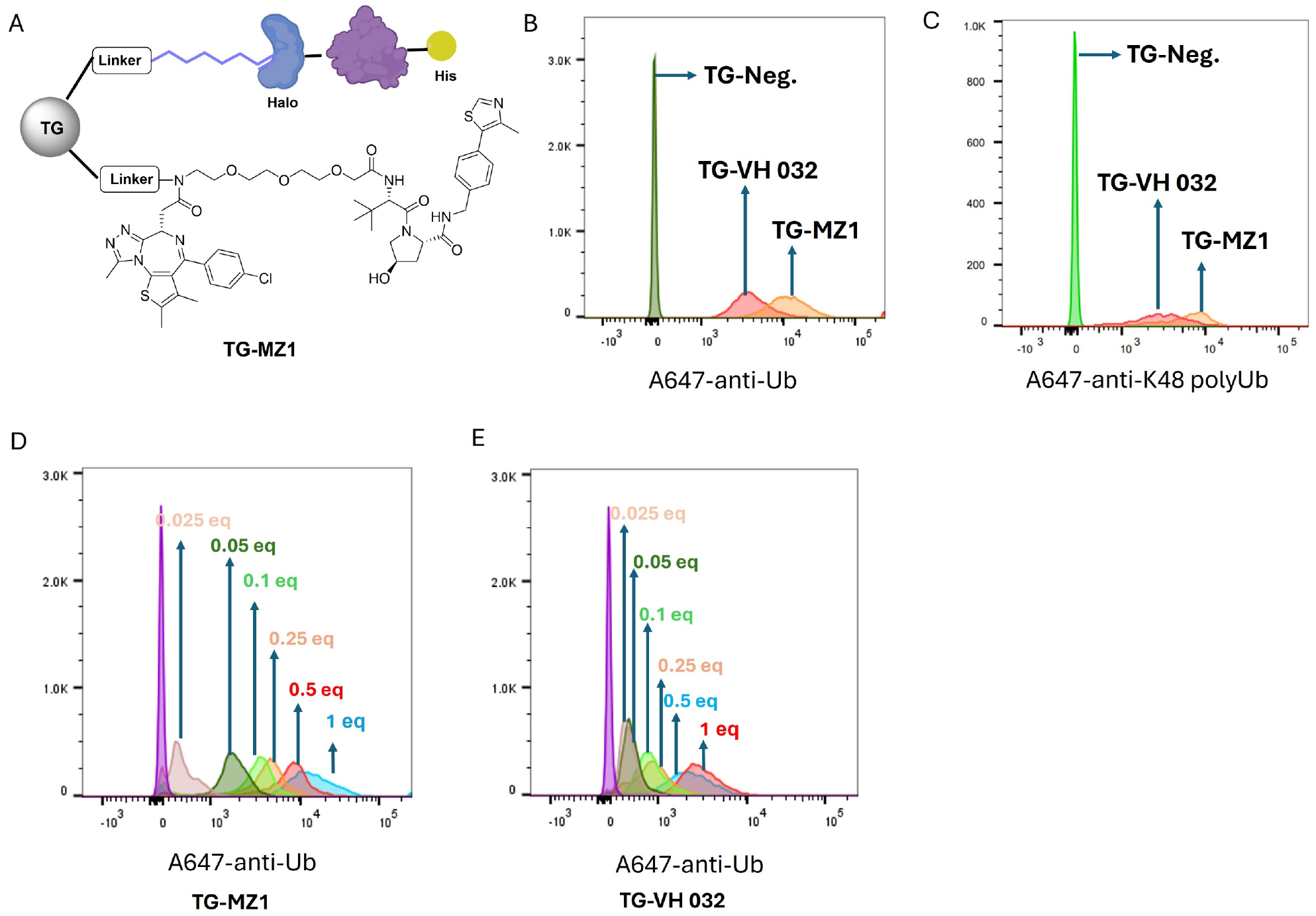
On-resin comparison of the degree of substrate protein Ubiquitylation mediated by an E3 ligase ligand and a PROTAC VHL E3 ligands and PROTACs. (A) The structure of TG-MZ1. (B&C) Flow cytometry analysis of indicated beads after loading with Halo-BRD4bd2-His_6_, and incubation with “one equivalent” the Ubiquitylation machinery at 37°C for 18 hours followed by staining with Alexa Fluor 647 labeled anti-Ubiquitin antibody or anti-K48 poly-Ubiquitin antibody. “1 eq.” represents Ube1 (50 nM), UbcH5a (500 nM), Neddylated CUL2/RBX, Elongin B/Elongin C/VHL (20 nM), Ubiquitin (Ub; 20 μM) and ATP (1 mM). (D & E) Effect of enzyme dilution on the degree of substrate poly-Ubiquitylation supported by immobilized VH032 and MZ1. The numbers shown indicate the concentrations of the E1, E2, and E3 enzymes relative to “one equivalent” For example, 0.05 equivalents represents a 20-fold dilution.

### Application to the Cereblon E3 ligase complex

Cereblon is another E3 Ubiquitin ligase commonly recruited by protein degraders.^21^ To determine if this assay format is also suitable for the analysis of ligand-Cereblon interactions, we created TG beads that display HTP-BRD4bd2-His_6_ and either the Cereblon ligand Pomalidomide, a crippled derivative of Pomalidomide in which the glutaramide nitrogen is methylated, or an Cereblon-based PROTAC dBET1 for BRD4 (Figure 5A). These beads were incubated for 18 hours at 37°C with purified 0.05 μM Ube1 (E1), 0.5 μM UbcH5a (E2), 0.02 μM neddylated CRBN E3 ligases complex (Neddylated CUL4A/RBX, DDB1/CRBN) and 20 μM ubiquitin and 1 mM ATP. Again, after thorough washing and staining with A647-labeled anti-Ub antibody, the beads were analyzed by flow cytometry. As shown in Figure 5B, the TG beads displaying dBET1 displayed a higher level of Ubiquitin-dependent fluorescence than the beads displaying Pomalidomide, and the methylated derivative (TG-mePom.) has baseline ubiquitin signal). Again, a similar result was observed when the resin was probed with antibody recognizing K48-linked poly-Ub chains (Figure 5C).

**Figure 5:**
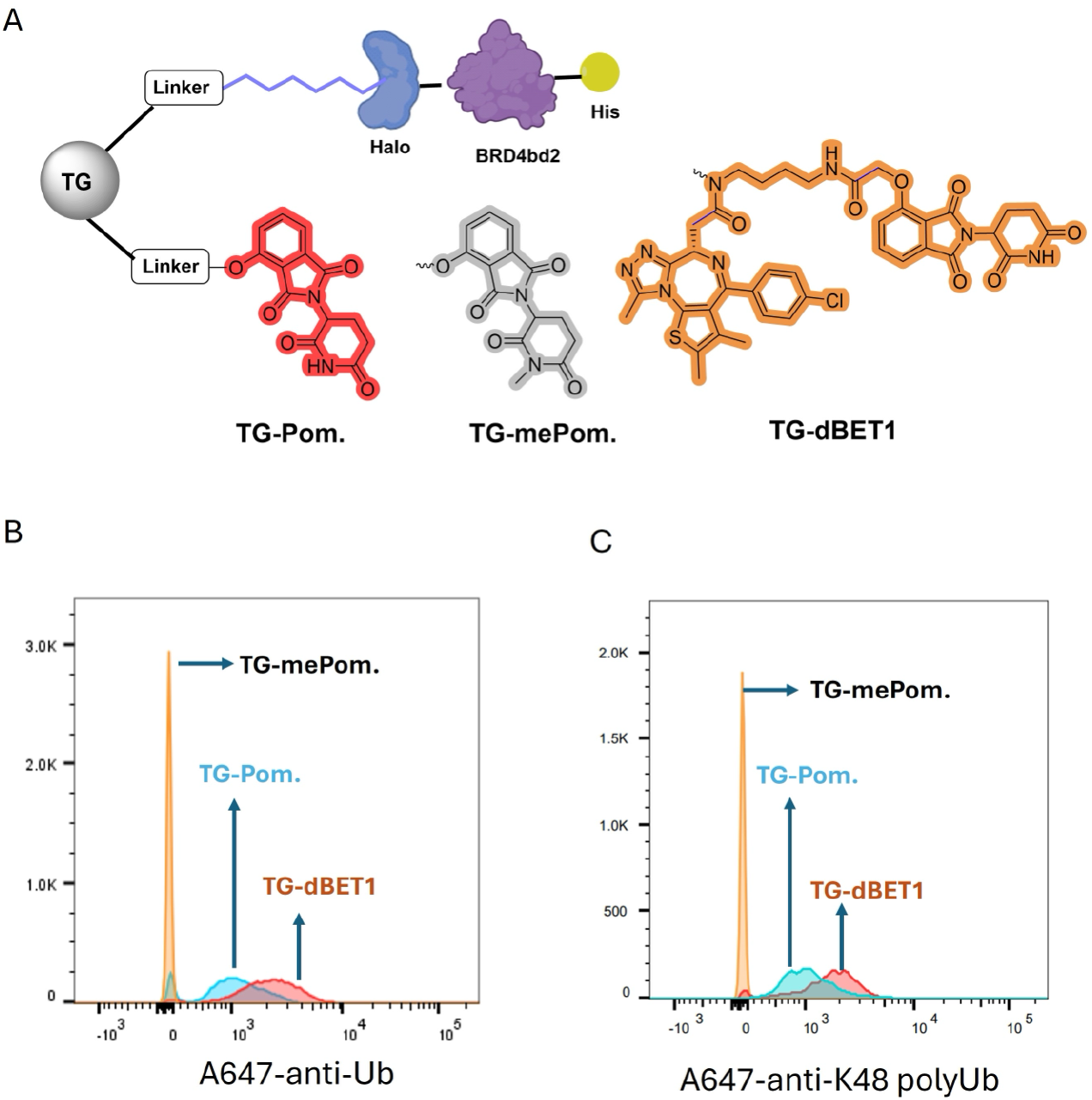
On-resin Ubiquitylation of HTP-BRD4bd2-His_6_ by Cereblon. (A) The structure of TG-Pom, TG-mePom, TG-dBET1. (B &C) Flow cytometry analysis of Alexa 647 signal of indicated beads after being loaded with Halo-BRD4bd2-His, and incubated with 0.05 μM Ube1 (E1), 0.5 μM UbcH5a (E2), 0.02 μM Neddylated CRBN E3 ligases complex (Neddylated CUL4A/RBX, DDB1/CRBN) and 20 μM ubiquitin and 1 mM ATP, then stained with Alexa647 labeled anti-ubiquitin antibody or anti-K48 polyubiquitin antibody.

### Discovery of a novel VHL ligand

Having established effective conditions for on-resin peptide or protein poly-Ubiquitylation, we turned to the application of this technique to the discovery of a novel VHL ligand using a typical microtiter plate-based high-throughput screening format. 92 hydroxyproline-containing compounds were synthesized by parallel solid-phase synthesis on 10 μm TG beads in a 96 well microtiter filter plate using the building blocks shown in Figure 6. The beads also displayed a chloroalkane for substrate attachment. The library contained four possible amino acids on the C-terminal side of the central hydroxyproline and 23 different units, derived from different amino acids, on the N-terminal side. All of the molecules were C-terminally acylated. Beads displaying an acetyl group or VH032 were included as negative and positive controls, respectively.

**Figure 6:**
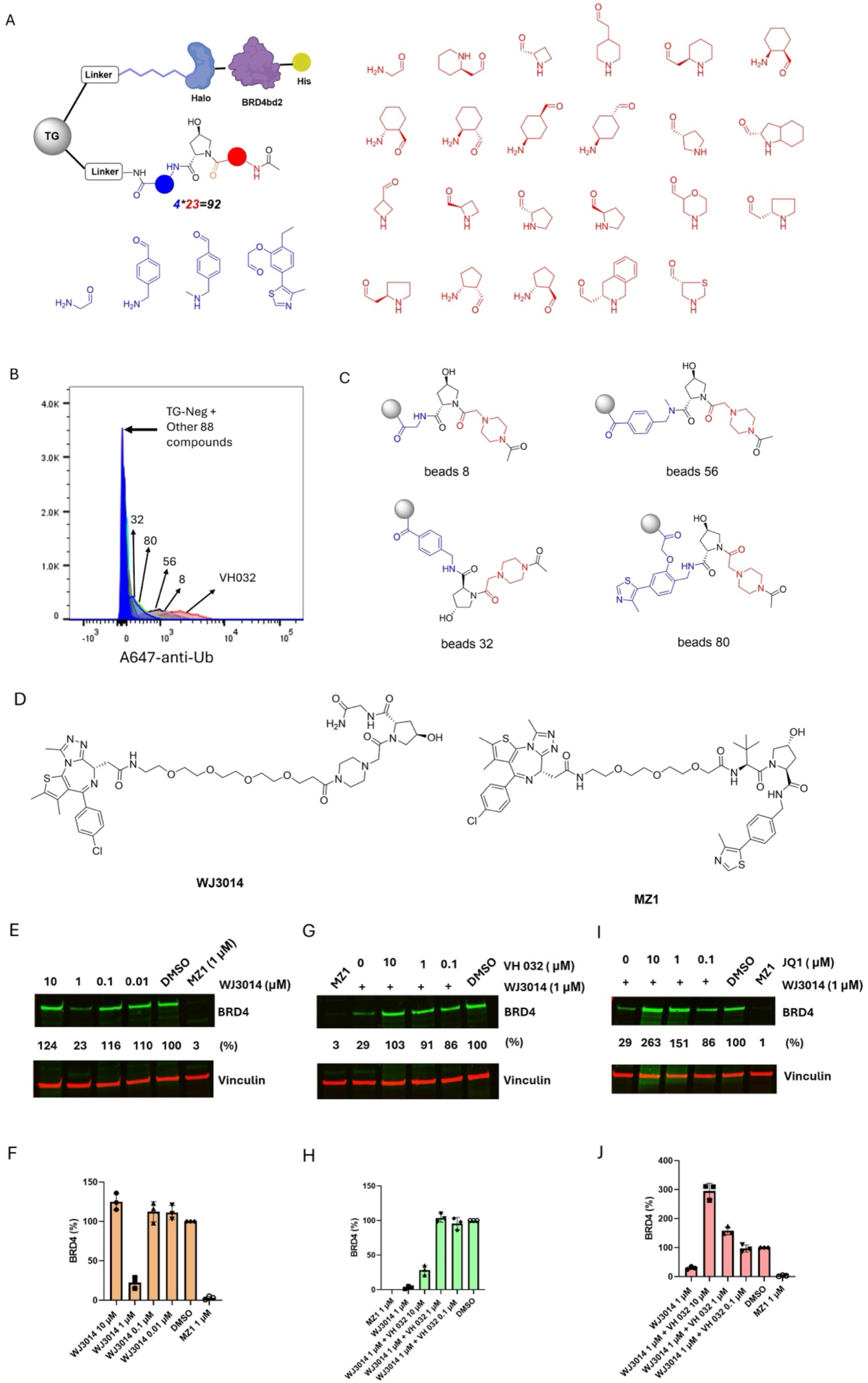
Discovery of a novel VHL ligand using the on-resin Ubiquitylation assay. See text for details. (A) Schematic of the beads employed in the screen, as well as the building blocks employed to construct the library by parallel solid-phase synthesis. (B) Flow cytometry analysis of the degree of Ubiquitylation-dependent fluorescence in the 647 nm channel after incubation with the E1, E2, VHL E3 enzymes, Ubiquitin and ATP under the same conditions used in Fig. 2. (C) Chemical structure of the screening hits **8, 56, 32** and **80**. (D) The chemical structure of WJ3014 and MZ1. (E &F) Western blot analysis of the level of BRD4 resulting from exposure of HeLa cells to the indicated concentration of WJ3014 or MZ1 for 12 h. (G & H) Western blot analysis of the level of BRD4 in HeLa cells resulting from coincubation of 1 μM WJ3014 or MZ1 with the indicated concentration of VH 032 to for 12 h. (I & J) Western blot analysis of the level of BRD4 by coincubation of 1 μM WJ3014 with indicated concentration of JQ1 to HeLa cells for 12 h. DMSO and MZ1 (1 μM) were used as vehicle and positive controls. The amount of BRD4 normalized to the vehicle control (DMSO) is indicated as “%”. Three biological replicates were performed. The Western blots shown are typical of all of the results obtained. The data shown in the quantification bar graph is the individual value of each replicate.

After loading HTP-BRD4bd2-His_6_ protein onto the beads, about 10^5^ beads in each well were transferred to an assay plate and incubated with the E1 and E2 enzymes, the VHL E3 complex, Ub and ATP for 18 hours at 37°C. After thorough washing, the beads in each well were stained with A647-labeled anti-Ub antibody, washed again, and analyzed using a flow cytometer. As shown in Figure 6B, the intensity of Ub-dependent fluorescence on 88 of the 92 beads was indistinguishable from that displayed by the negative control. Two sets of beads displayed weak fluorescence that was barely above background and two others displayed a stronger signal that was well separated from the background but was not as strong as that seen for VH032. The structures of the two weak hits (compounds **80** and **32**) and the two apparently stronger hits (compounds **8** and **56**), are shown in Figure 6C. All four compounds shared the same unit on the N-terminal side of hydroxyproline (highlighted in red).

We decided to focus on compound **8** for further testing given it was one of the apparently more potent hits as well as it’s extraordinarily low molecule weight (355 Da). To determine if this molecule is indeed a VHL ligand capable of supporting targeted protein degradation, a series of JQ1-**8** chimeras with different linkers separating them (see Supplementary Figure S7) were synthesized. Each (at both 1 μM and 10 μM) was tested for their ability to trigger BRD4 degradation in HeLa cells over a twelve-hour period. Three conjugates showed activity at 1 μM (Supplementary Figure S8) with WJ3014 displaying the best activity. The chemical structure of WJ3014 is shown in Figure 6D, along with that of the established PROTAC MZ1, for comparison. At a concentration of 1 μM, WJ3014 reduced the level of native BRD4 to 23% that observed in cells treated with vehicle (Figure 6E and 6F). MZ1 achieved 97% degradation under the same conditions (Figure 6E and 6F). Examination of the concentration dependence revealed, as expected, a pronounced hook effect^25^ (Figure 6E and 6F). At 10 μM WJ3014, greater-than-vehicle levels of BRD4 are seen due to competition of the all-important ternary complex by bivalent complexes.^25^ At concentrations of 10 nM and 100 nM WJ3014, the level of BRD4 was actually slightly higher than that observed for the vehicle control, presumably reflecting the well-known compensation mechanism employed by cells in which expression of BRD4 is increased in response to BRD4 inhibition by JQ1.^26^ As shown in Figure 6G and 6H, WJ3014-mediated degradation of BRD4 is competed by the addition of increasing amounts of VH032. Degradation was also abolished by the addition of free JQ1 (Figure 6I and 6J), as expected. These data show that compound **8** is indeed a VHL ligand and is capable of supporting targeted protein degradation in the context of a PROTAC.

## DISCUSSION

One of the major applications of molecular glues is to drive the enzyme-mediated post-translational modification of “neo-substrate” proteins. For this to occur, the ligand employed to engage the modifying enzyme must be either an agonist or an innocent ligand, not an inhibitor. We report here a powerful new screening platform that couples binding of a small molecule to a protein-modifying enzyme with a functional readout of that activity. Thus, inhibitors should not score as a hit in screens using this technology. The system is a variant of well-established one bead one compound (OBOC) library screening systems that have been used previously for ligand discovery, usually by incubating the beads with a labeled protein target and isolating beads that retain the label.^27^ In order to couple the binding event to a functional readout of enzyme activity, we modified the bead surface to display multiple copies of a neo-substrate. We hypothesized that engagement of the enzyme by a bead-displayed ligand would drive PTM of the immobilized protein in the concentrated microenvironment of the bead. After thorough washing, beads displaying a modified protein could be identified by staining with a fluorescently-labeled antibody that recognizes the PTM followed by analysis on a flow cytometer (Figure 1)

Given the importance of targeted protein degradation as a novel therapeutic modality, we chose to first develop this system in the context of E3 Ubiquitin ligase ligand discovery. Using the well-characterized VH032 ligand as a positive control, we demonstrated that incubation of neo-substrate-displaying beads with E1 and E2 enzymes, the VHL E3 ligase complex, Ubiquitin and ATP results in the poly-Ubiquitylation of the immobilized substrate. This was found to occur with both a minimalist peptide substrate containing one, two, or three lysines (Figure 2) and a fusion protein substrate (Figure 3). We also demonstrated that Cereblon-mediated poly-Ubiquitylation of a neo-substrate can be readily observed using this platform (Figure 5).

We then proceeded to apply this system to screening a model library of 92 compounds, each containing a central hydroxyproline (Figure 6), for engagement of VHL. This library was created by parallel solid-phase synthesis in a 96 well microtiter plate format and screened in the same one compound per well format. As shown in Figure 6B, when exposed to the Ubiquitylation machinery, four of these compounds mediated above-background poly-Ubiquitylation of immobilized HTP-BRD4bd2-His6 protein. All of these species shared the same N-terminal building block, suggesting that this unit is critical for activity. Of the two apparently more potent hits, compound **8** was of interest due to its simple structure and low molecular mass of 355 Da (for comparison the mass of VH032 is 430 Da). A series of compounds in which **8** was tethered to the BRD ligand JQ1 via linkers of different lengths was synthesized and tested for the ability to support degradation of BRD4 in HeLa cells. WJ3014 was found to have the best activity and reduced the level of native BRD4 in Hela cells to 23% that found in vehicle-treated cells (Figure 7). This activity was potently competed by VH032, demonstrating that WJ3014 functions by recruiting the VHL E3 ligase to BRD4 and that **8** is indeed a VHL ligand. WJ3014 is very likely a weaker ligand than VH032, but its low molecule weight and simple, modular structure make it a good starting point for further elaboration into a higher affinity compound.

As indicated in Figure 1, this assay format should also register molecular glues as screening hits. Indeed, given that BRD4 is a reader protein that recognizes acetylated lysine residues^20^ and that the acylated piperidine unit in compound **8** might be considered as a possible acetyl lysine analogue, we checked compound **8** (WJ3004) for molecular glue activity. These data are shown in Supplementary Figure S9. It is possible that **8** is a weak molecular glue degrader as some loss of BRD4 was seen in some experiments. But the data did not achieve statistical significance, and a more definite conclusion will have to await further development of the compound.

Highly relevant to detection of glues using this assay are the data shown in Figure 4. We compared the level of Ubiquitylation of immobilized HTP-BRD4-bd2-His6 by the VHL E3 ligase complex on beads displaying either VH032 or the MZ1. Under the standard conditions there was little difference but as the enzymes were diluted a much greater loss of signal on the VH032-displaying beads than the MZ1-displaying beads was observed. We attribute this result to the fact that the PROTAC MZ1 is capable of forming a ternary complex with the substrate and the ligase, whereas VH032 can only bind the ligase, not the bromodomains of BRD4. Because VH032 is only a modest affinity ligand for VHL (K_D_ = 185 nM), it seems reasonable to suggest that the additional free energy provided by the neo-protein-protein interactions between the ligase and the immobilized substrate in the ternary complex endows the MZ1/bromodomain-displaying beads with an effectively higher affinity for the soluble ligase complex. The practical impact of this finding is that screens done at relatively low enzyme concentrations are likely to favor the recovery of glues and not simple E3 ligands, in the hit pool, unless the E3 ligands are of high affinity.

This work represents a rare example of OBOC libraries being employed in a functional screening format but it is not the only one. Paegel and colleagues screened OBOC DNA-encoded libraries^28^ in a functional fashion by encapsulating beads into microfluific droplets along with an enzyme and a fluorogenic substrate.^29^ In these studies, the compound was linked to the resin by a photocleavable linker, allowing release of the molecules from the beads by moving the droplets past a laser in a microfluidic device.^30^ Droplets in which an enzyme inhibitor was released from the resin displayed a lower level of fluorescence than those where a non-inhibitor was released. These droplets could be sorted and amplification and sequencing of the encoding DNA strands in them revealed the identity of the inhibitors.

Even more relevant to this study is pioneering work done by Meldal and colleagues over 20 years ago.^31, 32^ They demonstrated that PEG-based beads carrying a fluorogenic protease substrate in which a donor and quencher were separated by a peptide linker, as well as a small molecule, could be employed to identify compounds that act as protease inhibitors. In this system, beads that carried non-binders acquired a high level of fluorescence when exposed to the protease due to linker cleavage and subsequent release of the quencher. Beads carrying a protease inhibitor remained dark. This work introduced the concept of an assay format in which an enzyme substrate and a small molecule are co-immobilized on a bead, which we also employ here.

However, both of these assay systems differ fundamentally from ours in that they are designed to identify enzyme inhibitors and thus serve as an alternative to traditional HTS-based methods. The critical difference is that this earlier work employed good substrates for the enzyme of interest, while we purposely use a “bad” substrate (indeed, here a neo-substrate) so that in trans modification of the bead-displayed substrate by the soluble enzyme is negligible without the help of the co-immobilized small molecule. This allows one to observe the critical coupling of a ligand-enzyme biding event with a functional readout, which is the unique aspect of this system.

As mentioned above, DNA-encoded library (DEL) technology has been applied to the OBOC format^28^ and binding screens have been reported in which ligand-displaying beads have been isolated using fluorescence-activated cell sorting.^33-37^ Thus, this assay should be immediately transferable to screening OBOC DELs for molecular glues and E3 ligase ligands. These studies are underway. It also seems likely that this assay format should be effective for the discovery of non-inhibitory ligands to other PTM enzymes, such as kinases, phosphatases, Deubiquitylases, etc. so long as the bead displays a high K_M_ substrate that is not modified efficiently without small molecule-enzyme association. These studies are also in progress.

## Supporting information

Supplementary Information for article

## Acknowledgements

This work was supported by a grant from the National Institutes of Health (GM R35 GM151875). We thank Joel Cresser-Brown (Biotechne) for aid in obtaining the purified Ubiquitylation enzymes at a reduced cost, which was essential for completing this work.

## Notes

### Competing Interest Statement

T.K. is a co-founder and significant shareholder in Triana Biomedicines.

## References

1. Schreiber, S.L. The Rise of Molecular Glues. Cell 184, 3–9 (2021).

2. Sakamoto, K.M. et al. Development of Protacs to target cancer-promoting proteins for ubiquitination and degradation. Mol Cell Proteomics 2, 1350–1358 (2003).

3. Lai, A.C. & Crews, C.M. Induced protein degradation: an emerging drug discovery paradigm. Nature reviews. Drug discovery 16, 101–114 (2017).

4. Lu, G. et al. The Myeloma Drug Lenalidomide Promotes the Cereblon-Dependent Destruction of Ikaros Proteins. Science 343, 305–309 (2014).

5. Dong, G., Ding, Y., He, S. & Sheng, C. Molecular Glues for Targeted Protein Degradation: From Serendipity to Rational Discovery. Journal of Medicinal Chemistry 64, 10606–10620 (2021).

6. Han, T. et al. Anticancer sulfonamides target splicing by inducing RBM39 degradation via recruitment to DCAF15. Science 356 (2017).

7. Siriwardena, S.U. et al. Phosphorylation-Inducing Chimeric Small Molecules. Journal of the American Chemical Society 142, 14052–14057 (2020).

8. Chen, P.-H. et al. Modulation of Phosphoprotein Activity by Phosphorylation Targeting Chimeras (PhosTACs). ACS Chem. Biol. 16, 2808–2815 (2021).

9. Liu, X. & Ciulli, A. Proximity-Based Modalities for Biology and Medicine. ACS Central Science 9, 1269–1284 (2023).

10. Henning, N.J. et al. Deubiquitinase-targeting chimeras for targeted protein stabilization. Nat Chem Biol 18, 412–421 (2022).

11. Lam, K.S. et al. A new type of synthetic peptide library for identifying ligand-binding activity. Nature 354, 82–84 (1991).

12. Kodadek, T. Protein-detecting microarrays. Trends Biochem. Sci. 27, 295–300 (2002).

13. Bradner, J.E. et al. A robust small-molecule microarray platform for screening cell lysates. Chem. & Biol. 13, 493–504 (2006).

14. Neri, D. & Lerner, R.A. DNA-Encoded Chemical Libraries: A Selection System Based on Endowing Organic Compounds with Amplifiable Information. Annual review of biochemistry 87, 479–502 (2018).

15. Clark, M.A. et al. Design, synthesis and selection of DNA-encoded small-molecule libraries. Nature Chem Biol 5, 647–654 (2009).

16. Liu, S. et al. Rational Screening for Cooperativity in Small-Molecule Inducers of Protein–Protein Associations. Journal of the American Chemical Society (2023).

17. Kamura, T. et al. Activation of HIF1alpha ubiquitination by a reconstituted von Hippel-Lindau (VHL) tumor suppressor complex. Proc Natl Acad Sci U S A 97, 10430–10435 (2000).

18. Gadd, M.S. et al. Structural basis of PROTAC cooperative recognition for selective protein degradation. Nature Chemical Biology 13, 514–521 (2017).

19. Soares, P. et al. Group-Based Optimization of Potent and Cell-Active Inhibitors of the von Hippel-Lindau (VHL) E3 Ubiquitin Ligase: Structure-Activity Relationships Leading to the Chemical Probe (2S,4R)-1-((S)-2-(1-Cyanocyclopropanecarboxamido)-3,3-dimethylbutanoyl)-4-hydroxy-N-(4-(4-methylthiazol-5-yl)benzyl)pyrrolidine-2-carboxamide (VH298). J Med Chem 61, 599–618 (2018).

20. Filippakopoulos, P. et al. Selective inhibition of BET bromodomains. Nature 468, 1067–1073 (2010).

21. Winter, G.E. et al. Phthalimide conjugation as a strategy for in vivo target protein degradation. Science 348, 1376–1381 (2015).

22. Los, G.V. et al. HaloTag: a novel protein labeling technology for cell imaging and protein analysis. ACS chemical biology 3, 373–382 (2008).

23. Diehl, C.J. & Ciulli, A. Discovery of small molecule ligands for the von Hippel-Lindau (VHL) E3 ligase and their use as inhibitors and PROTAC degraders. Chem Soc Rev 51, 8216–8257 (2022).

24. Zengerle, M., Chan, K.H. & Ciulli, A. Selective Small Molecule Induced Degradation of the BET Bromodomain Protein BRD4. ACS chemical biology 10, 1770–1777 (2015).

25. Douglass, E.F., Miller, C.J., Sparer, G., Shapiro, H. & Spiegel, D.A. A Comprehensive Mathematical Model for Three-Body Binding Equilibria. Journal of the American Chemical Society 135, 6092–6099 (2013).

26. Lu, J. et al. Hijacking the E3 Ubiquitin Ligase Cereblon to Ehiciently Target BRD4. Chem Biol 22, 755–763 (2015).

27. Sikder, K. & Kodadek, T. Optimized protocols for the isolation of specific protein-binding peptides or peptoids from combinatorial libraries displayed on beads. Molecular BioSystems 2, 25–35 (2006).

28. MacConnell, A.B., McEnaney, P.J., Cavett, V.J. & Paegel, B.M. DNA-Encoded Solid-Phase Synthesis: Encoding Language Design and Complex Oligomer Library Synthesis. ACS Comb Sci 17, 518–534 (2015).

29. Cochrane, W. et al. Activity-Based DNA-Encoded Library Screening. ACS Comb Sci 21, 425–435 (2019).

30. Price, A.K., MacConnell, A.B. & Paegel, B.M. hvSABR: Photochemical dose-response bead screening. Anal. Chem. 88, 2904–2911 (2016).

31. Graven, A. et al. Combinatorial library of peptide isosters based on Diels-Alder reactions: identification of novel inhibitors against a recombinant cysteine protease from Leishmania mexicana. J. Comb. Chem. 3, 441–452 (2001).

32. Meldal, M. The one-bead two-compound assay for solid phase screening of screening of combinatorial peptide libraries. Biopolymers (Peptide Sci.) 66, 93–100 (2002).

33. Mendes, K. et al. High-throughput identification of DNA-encoded IgG ligands that distinguish active and latent mycobacterium tuberculosis infections. ACS Chem. Biol. 19, 234–243 (2017).

34. Castanha, P.M.S. et al. Identification and characterization of a non-biological small molecular mimic of a Zika virus conformational neutralizing epitope. Proc. Natl. Acad. Sci. USA 121, In press (2024).

35. McEnaney, P., Balzarini, M., Park, H. & Kodadek, T. Structural characterization of a peptoid-inspired conformationally constrained oligomer (PICCO) bound to streptavidin. Chemical Communications 56, 10560–10563 (2020).

36. Koesema, E. et al. Synthesis and Screening of a DNA-Encoded Library of Non-Peptidic Macrocycles. Angew Chem Int Ed Engl, e202116999 (2022).

37. Benhamou, R.I. et al. DNA-encoded library versus RNA-encoded library selection enables design of an oncogenic noncoding RNA inhibitor. Proc Natl Acad Sci U S A 119 (2022).

