## Supplementary Information for article for "A Functional Assay For Mining Non-Inhibitory Enzyme Ligands From One Bead One Compound Libraries: Application to E3 Ubiquitin Ligases"

**General information.**

Chemical reagents were purchased from Fisher Scientific, Millipore Sigma, ChemScene and Ambeed. $^{1}H$ and $^{13}C$ NMR spectra were obtained with Bruker AV400 NMR Spectrometer or Bruker AV600 NMR Spectrometer with a CryoProbe. Chemical shifts are reported in δ (ppm) units and $^{13}C$ and $^{1}H$ signals from deuterated solvents were used as references. 60 F254 (Merck) silica plate was used in thin layer chromatography (TLC) analysis.

Agilent 1100 series HPLC equipped with Agilent ZORBAX SB-C18 column (Part number: 861953-902, 4.6*100 mm. 3.5 µm) and an electrospray ionization (ESI) source (Agilent Technologies 6120 Quadrupole) was used to obtain mass spectra for small molecules. The following methods were used to analyze the purity of the samples in the LC-MS instrument. Solvent A contains 100% water (0.1% formic acid) and solvent B contains 100% acetonitrile (0.1% formic acid).

| Time (min) | A [%] | B [%] | Flow [mL/Min] |
|---|---|---|---|
| 0.0 | 100.0 | 0.0 | 1.0 |
| 2.0 | 100.0 | 0.0 | 1.0 |
| 12.0 | 0.0 | 100.0 | 1.0 |
| 15.0 | 0.0 | 100.0 | 1.0 |

Small molecule column chromatography was conducted on silica gel (230-400 mesh) or a SunFire Prep C18 column (100Å, 5 µm, 19 mm X 250 mm, Waters) was used at flow rate of 9 mL/min. The UV absorbance at 214, 254, 298 nm were used to monitor the purification process. Water (with 0.1% Trifluoroacetic acid) and acetonitrile (with 0.1% Trifluoroacetic acid) were used as solvent A and B respectively. The following methods were used to purify the samples.

| Time (min) | A [%] | B [%] | Flow [mL/Min] |
|---|---|---|---|
| 0.0 | 95.0 | 5.0 | 9.0 |
| 5.0 | 95.0 | 5.0 | 9.0 |
| 40.0 | 5.0 | 95.0 | 9.0 |
| 50.0 | 95.0 | 5.0 | 9.0 |
| 51.0 | 95.0 | 5.0 | 9.0 |
| 60.0 | 95.0 | 5.0 | 9.0 |

**General solid phase synthesis protocol:**

Compounds were synthesized on TentaGel resin using general SPS protocol described below. The compounds displayed on the beads were synthesized on 10 µm TentaGel M NH2 resin (Rapp-Polymere ,0.30 mmol/g) mixed with 10% of 300 µm TentaGel MB RAM resin (Rapp-Polymere, 0.19 mmol/g) for mass analysis. The beads were transferred to a fritted spin-column (Mobicol Classic, large filter, 10 µm pore size) and swelled in DMF (1 h, RT). FMOC was removed with 20% piperidine in DMF (3x resin volume after swelling, 500 µL) at RT for 10 minutes, twice. It was then washed thoroughly with DMF, DCM and DMF, and successively coupled with either amino acids or peptoid units. In general, acids (5 times of resin capacity, 1.0 mmol) were pre-activated at RT for 5 minutes with DIC/Oxyma/Collidine (1.4/1.0/1.0 mmol) in DMF, added to the resin, incubated at 37°C for one hour, and washed with DMF, DCM (5 times each, 5x resin volume). FMOC was removed where applicable.

**Supplemental Methods:**

**On-beads ubiquitination staining and flow cytometry analysis:** 10 µm TentaGel beads were stored in DMF after equipping with the chemicals indicated. $10^5$ beads were aliquot out and dissolved in 1 mL PBST buffer (0.2 g/L KCl, 0.24 g/L KH2PO4, 8 g/L NaCl, 1.44 g/L Na2HPO4 anhydrous, 0.05% Tween-20, pH 7.4). Spin down and aliquot out the supernatant. Multiscreen® 96 well Plate equipped with hydrophilic PVDF membrane (MSBN12 from Millipore Sigma) was activated by EtOH and washed by PBST buffer. Then the beads were transferred into the plate and washed three times with PBST, incubated with 150 µL PBST at 4 °C for three hours, washed three times with PBST, and the beads were blocked by 150 µL StartingBlock (PBS) blocking buffer (ThermoFisher, Cat: 37538) overnight. Then, the beads were washed three times with blocking buffer and three times with ubiquitination buffer (20 mM HEPES, 150 mM NaCl, 10 mM MgCl2, pH 7.5). Then the beads were suspended in 100 µL ubiquitination buffer, indicated concentration of E1, E2, E3, ubiquitin and 1 mM ATP were added to the beads and incubated at 37 °C for 18 hours.

Then the beads were washed with PBST 6 times, and incubated with 1 µL Alexa Fluor® 647 anti-Ubiquitin Antibody (Biolegend, Cat: 838710, 0.5 mg/mL) in PBST buffer for 2 hours at 4 °C. Then the beads were washed with PBST 6 times and analyzed by flow cytometry (BD LSR II). The Alexa Fluor 647 channel was selected for the analysis. The signal were plotted using FlowJo_V10.10.0

E1, Recombinant Human Ubiquitin Activating Enzyme (UBE1) (Cat: E-305); E2, Recombinant Human UbcH5a/UBE2D1 Protein (Cat: E2-616); Recombinant Human Elongin B/Elongin C/VHL Complex (Cat: E3-600); Recombinant Human CUL2/RBX1 Neddylated Complex Protein (Cat: E3-421); Recombinant Human CUL4A/RBX1 Neddylated Complex Protein (Cat: E3-441), Recombinant Human DDB1/CRBN Complex Protein (Cat: E3-500) were purchased from biotechne.

**Halo-BRD4bd2-His expression and purification:**

Halo-BRD4(BD2)-His gene sequence was purchased for IDT Gene Block, the gene was inserted to pET-3a vector using infusion cloning. Then the plasmid was transformed into BL21 cells and cultured at 37 °C until $OD_{600}$ reached 0.3-0.4. The incubation temperature decreased to 25 °C until $OD_{600}$ reached 0.6. Then, protein expression was induced with 1 mM IPTG. Cells were cultured for an additional 4 h at 25 °C after the induction and harvested by centrifugation at 6,000 rpm for 10 min at 4 °C. Cells were lysis and purified by Ni-NTA resin and desalted.

**Halo-BRD4bd2-His** amino acid sequence: the Halo tag protein sequence was highlighted as green and BRD4bd2 was heighted as red.

MSEIGTGFPFDPHYVEVLGERMHYVDVGPRDGTPVLFLHGNPTSSYLWRNIIPHVAPSHRCIAPDLIGMGKSDKPDLDYF
FDDHVRYLDAFIEALGLEEVVLVIHDWGSALGFHWAKRNPERVKGIACMEFIRPIPTWDEWPEFARETFQAFRTADVGR
ELIIDQNAFIEGALPKCVVRPLTEVEMDHYREPFLKPVDREPLWRFPNELPIAGEPANIVALVEAYMNWLHQSPVPKLLF
WGTPGVLIPPAEAARLAESLPNCKTVDIGPGLHYLQEDNPDLIGSEIARWLPALGGGGSGGGGSGGGSSGVDLGTENLY

FQSMKDVPDSQQHPAPEKSSKVSEQLKCCSGILKEMFAKKHAAYAWPFYKPVDVEALGLHDYCDIIKHPMDMSTIKSK LEAREYRDAQEFGADVRLMFSNCYKYNPPDHEVVAMARKLQDVFEMRFAKMPDEGSGSGSHHHHHH

**On-beads loading of Halo-BRD4bd2-His:** 10 µm TentaGel beads were washed and blocked as described above, then 10 µM Halo-BRD4bd2-His was incubated with the beads in 150 µL PBS blocking buffer for 12 hours at 4 °C. Then, the beads were washed with PBS blocking buffer for 3 times.

**Mammalian Cell Culture**. HeLa cell line (ATCC CCL-2) were cultured at 37 °C with 5% $CO_2$ in DMEM (10566016, Gibco) supplemented with 10% FBS (16140-071, ThermoFisher) in a humidified incubator. Adherent cells were passaged using TrypLE (12605-010, ThermoFisher) every 3 days.

**Immunoblotting Analysis of VHL and BRD4 protein level**. $3x10^5$ HeLa cells were plated in Corning Costar Flat Bottom Cell Culture 6-well plates (Mediatech, Inc., Manassas, VA) and allowed to adhere for 12 h in 2 mL DMEM medium supplemented with 10% FBS at 37 °C and 5% $CO_2$, then aliquot out the media and add 2 mL fresh DMEM medium supplemented with 10% FBS. Cells were treated with compounds for indicated time at 37°C at 5% $CO_2$. Post compound treatment, cells were washed 2x with DPBS (14190250, Thermo Fisher Scientific), and harvested by TrypLE™ Express Enzyme (1X), phenol red (12605010, Thermo Fisher Scientific). Harvested cells were used immediately for the following experiments or stored at -80°C. Cells were lysed with 20 µL Mammalian Permeabilization Reagent (MPER) (78501, Thermo Fisher Scientific) and incubated at RT for 10 minutes. Soluble protein supernatant was clarified by centrifugation at 15,000g for 15 minutes at 4 °C in a benchtop microcentrifuge (Eppendorf). Protein concentration was determined by Pierce™ Coomassie (Bradford) Protein Assay Kit (23200, Thermo Fisher Scientific). 40 µg cell lysate was loaded onto 4-15% or 4-20% gradient gels (Cat#4561083, BioRad). SDS-Page gels were run at 180V for 40 minutes, and transferring to nitrocellulose membrane was executed according to manufacturer's instructions (1704270, Bio-Rad). Membranes were blocked in 5% Blot-Quickblocker modified milk protein (786-011, G-Biosciences) in PBST buffer. For VHL western blotting, primary anti-VHL Rabbit (#68547, cell signaling, 1:1,000 dilution), anti-GAPDH antibody mouse monoclonal (sc-32233, Santa Cruz, 1:1,000 dilution) was incubated with membrane for 16 hours at 4 °C. For Brd4 western blotting, primary anti-BRD4 (E2A7X) Rabbit mAb (#13440, cell signaling, 1:1,000 dilution), anti-Vinculin antibody mouse monoclonal (SAB4200080, Millipore Sigma, 1:10,000) was incubated with membrane for 16 hours at 4 °C. Proteins were visualized using Licor Odyssey instrument after 1 hour RT incubation with IR dye conjugated antibodies: Donkey anti-mouse680RD (LicorCat#925-32212; RRID: AB_27116622) and Goat anti-rabbit800CW (Licor Cat#925-32211; RRID: AB_2651127) diluted 1:10,000 in Odyssey blocking buffer (Licor).

Western blots were quantified using ImageJ (RRID: SCR_003070) after imaging with LI-COR Odd-ysey system and LI-COR image studio software (RRID: SCR_015795). The quantification of BRD4 level is normalized to DMSO control as previous publication suggested.[1]

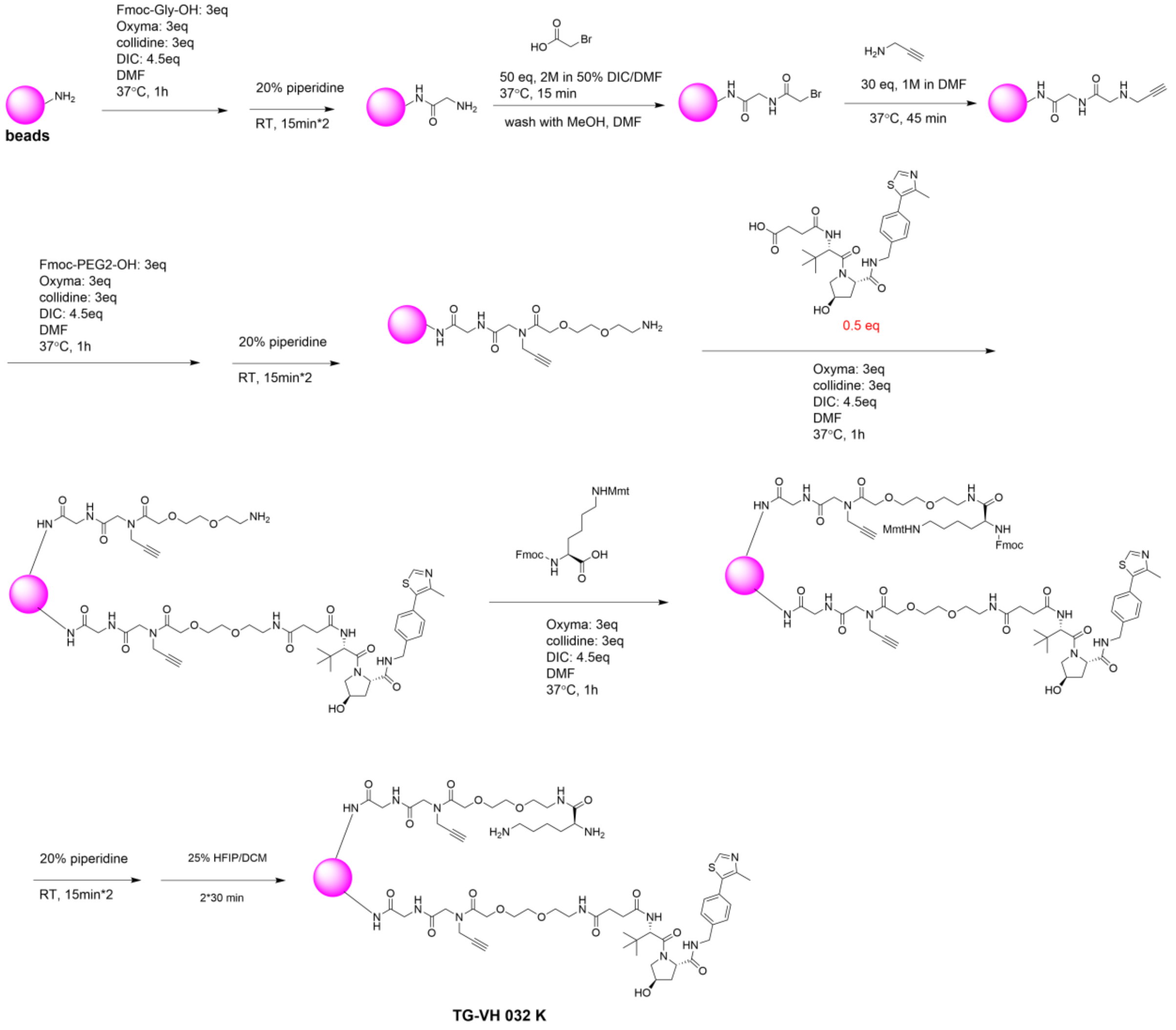


Figure S1: The scheme to synthesize TG-VH 032 K.

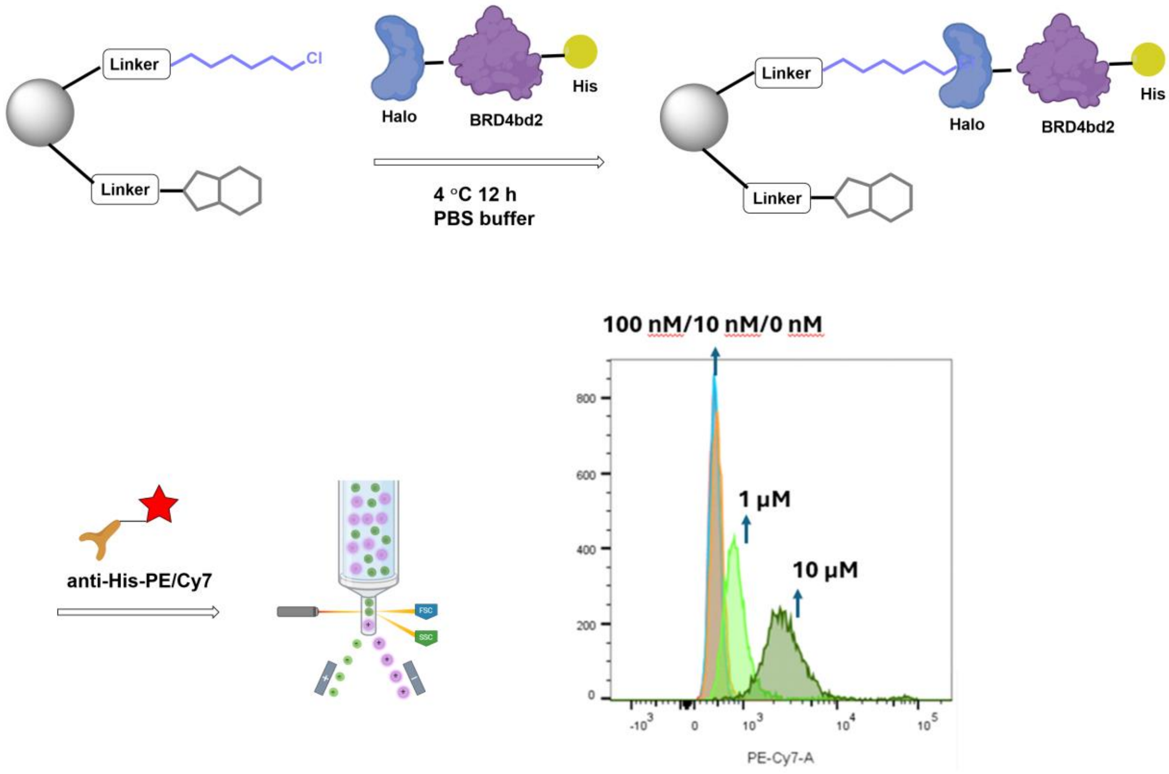


Figure S2: The procedure to load the substrate protein to the beads.

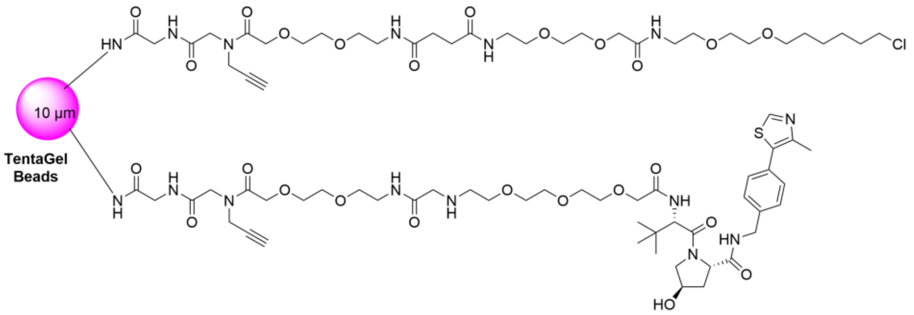


**TG-VH 032**

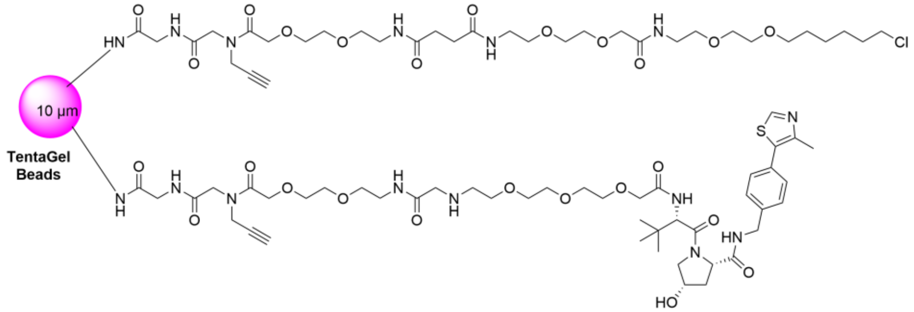


**TG-VH 032 (-)**

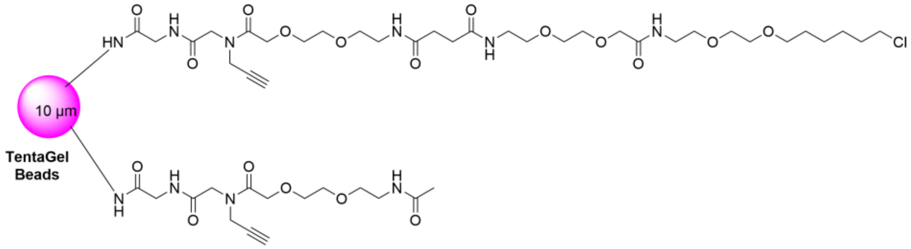


**TG-Neg.**

Figure S3: Chemical structure of TG-VH 032, TG- VH 032(1), TG-Neg.

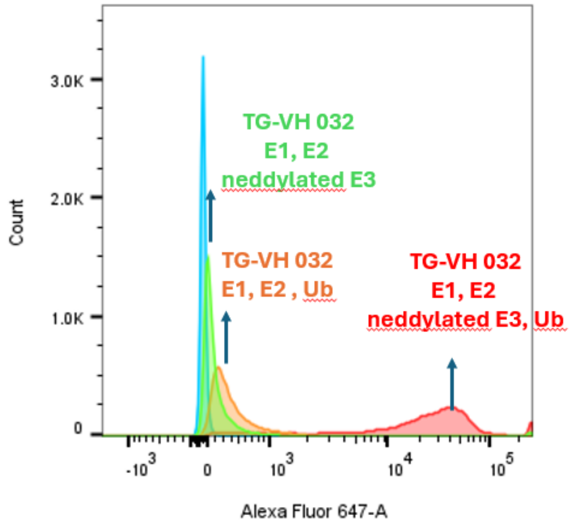


Figure S4: Flow cytometry analysis of Alexa Fluor 647 signal of TG-VH 032 after load the beads with 10 μM Halo-BRD4bd2-His, ubiquitinated under the conditions indicated, stained with Alexa Fluor 647 labeled anti-ubiquitin. The enzyme concentration use is 0.05 μM Ube1 (E1), 0.5 μM UbcH5a (E2), 0.02 μM neddylated VHL E3 ligases complex (Neddylated CUL2/RBX, Elongin B/Elongin C/VHL Complex) and 20 μM ubiquitin and 1 mM ATP in HEPES buffer, pH 7.2 at 37 ℃ for 18 h.

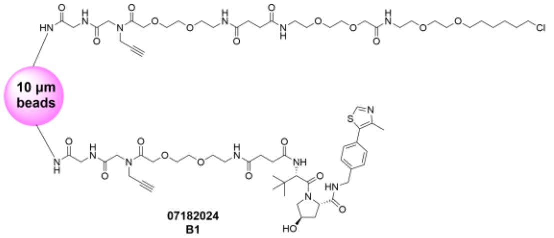


07182024
**B1**

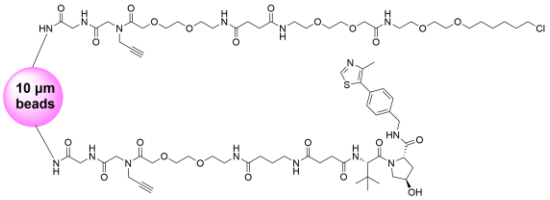


07182024
**B3**

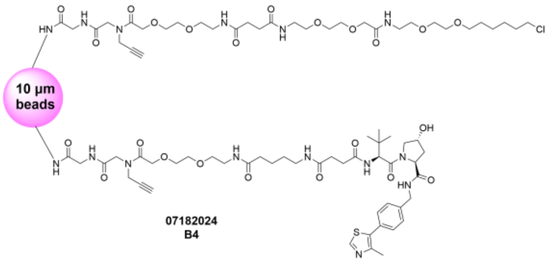


07182024
**B4**

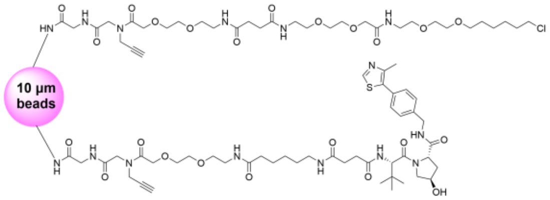


07182024
**B5**

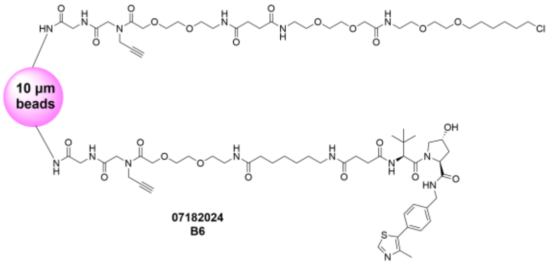


07182024
**B6**

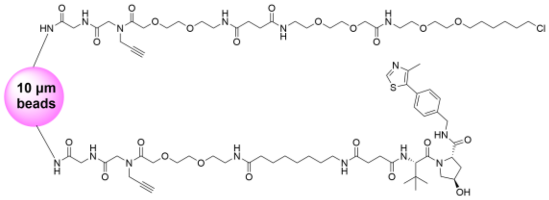


07182024
**B7**

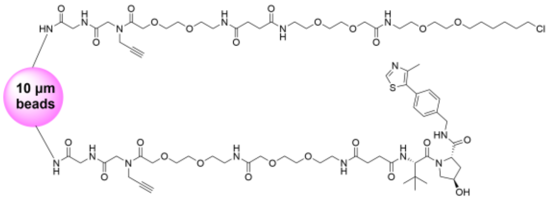


07182024
**B8**

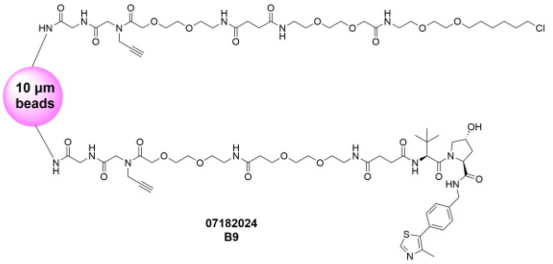


07182024
**B9**

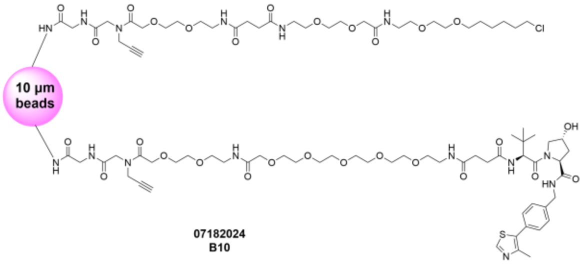


07182024
**B10**

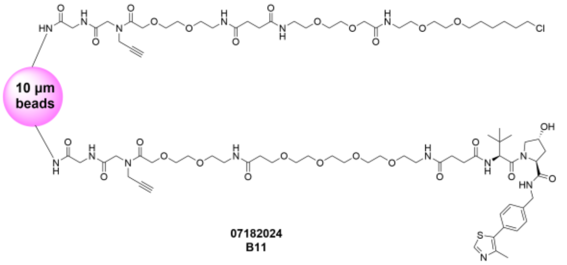


07182024
**B11**

Figure S5: Structures of TG-VH 032 with different linkers.

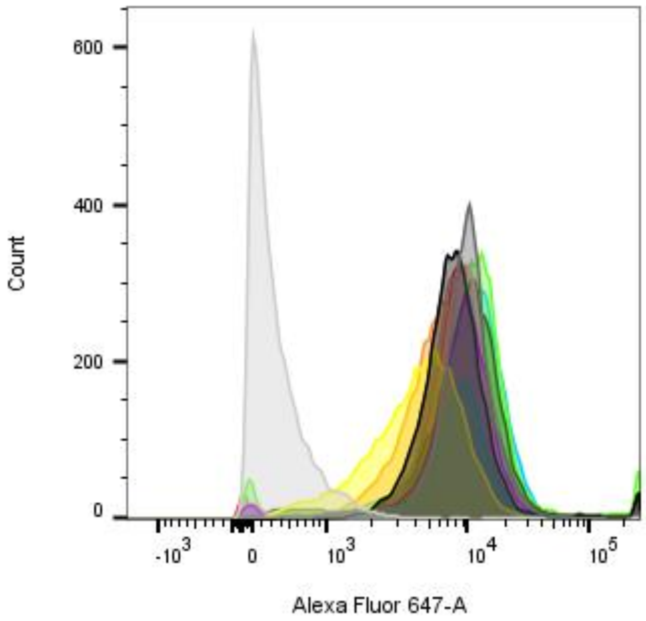


Figure S6: Flow cytometry analysis of Alexa 647 signal of TG-VH 032 with different linkers after loaded with 10 μM Halo-BRD4bd2-His, ubiquitinated with 0.05 μM Ube1 (E1), 0.5 μM UbcH5a (E2), 0.02 μM neddylated VHL E3 ligases complex (Neddylated CUL2/RBX, Elongin B/Elongin C/VHL Complex) and 200 μM ubiquitin and 1 mM ATP in HEPES buffer, pH 7.2 at 37 ℃ for 18 h, stained with Alexa Fluor 647 labeled anti-ubiquitin antibody.

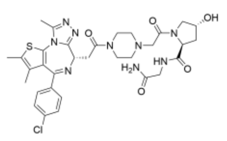

**WJ3006**

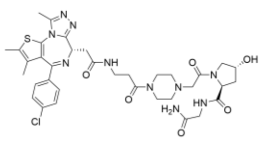

**WJ3007**

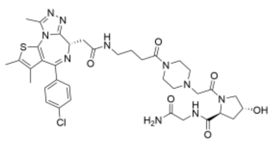

**WJ3008**

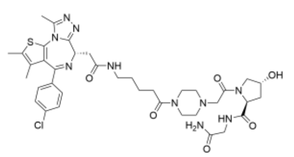

**WJ3009**

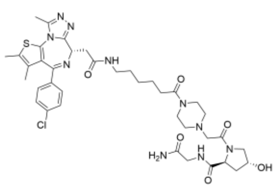

**WJ3010**

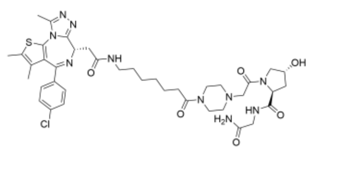

**WJ3011**

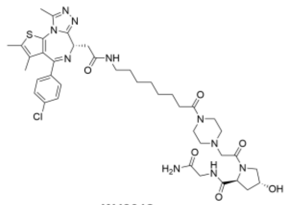

**WJ3012**

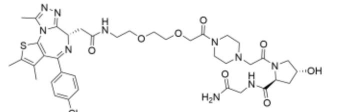

**WJ3013**

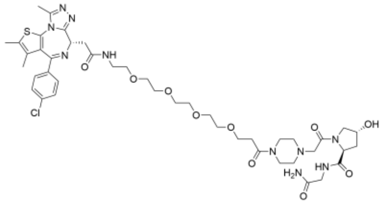

**WJ3014**

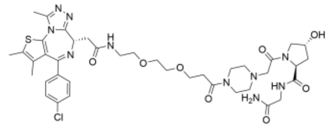

**WJ3015**

Figure S7: Chemical structure of WJ3006-WJ3015.

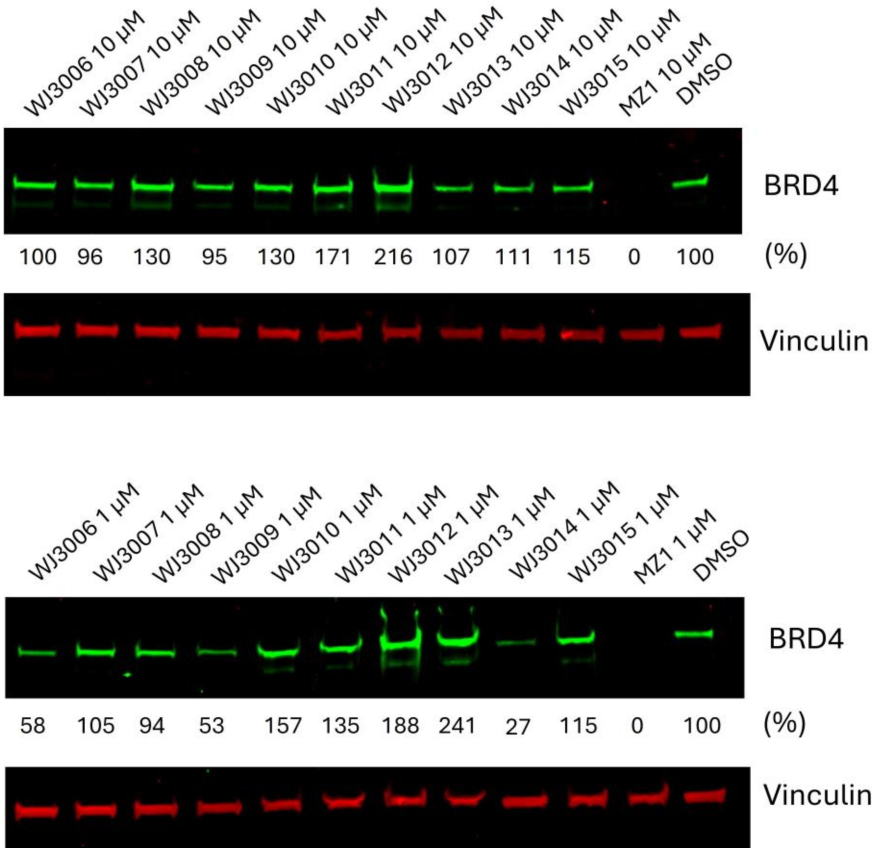


Figure S8: Western blot analysis of the level of BRD4 by adding the indicated concentration of compounds to HeLa cells and incubated for 12 h. DMSO was used for vehicle control. MZ1 was used for positive control.

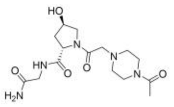


**WJ3004**

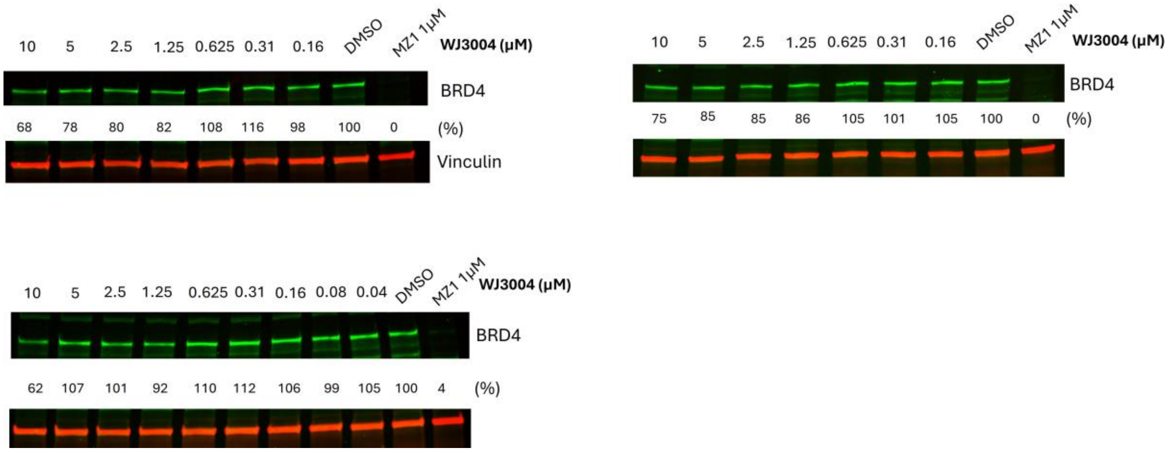


Figure S9: Western blot analysis of the level of BRD4 by adding the indicated concentration of WJ3014 to HeLa cells and incubated for 12 h. DMSO was used for vehicle control. MZ1 was used for positive control.

**LC/MS: TG-VH 032**

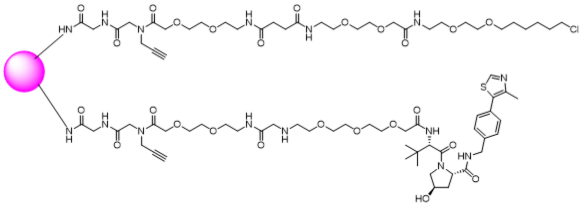

**TG-VH 032**

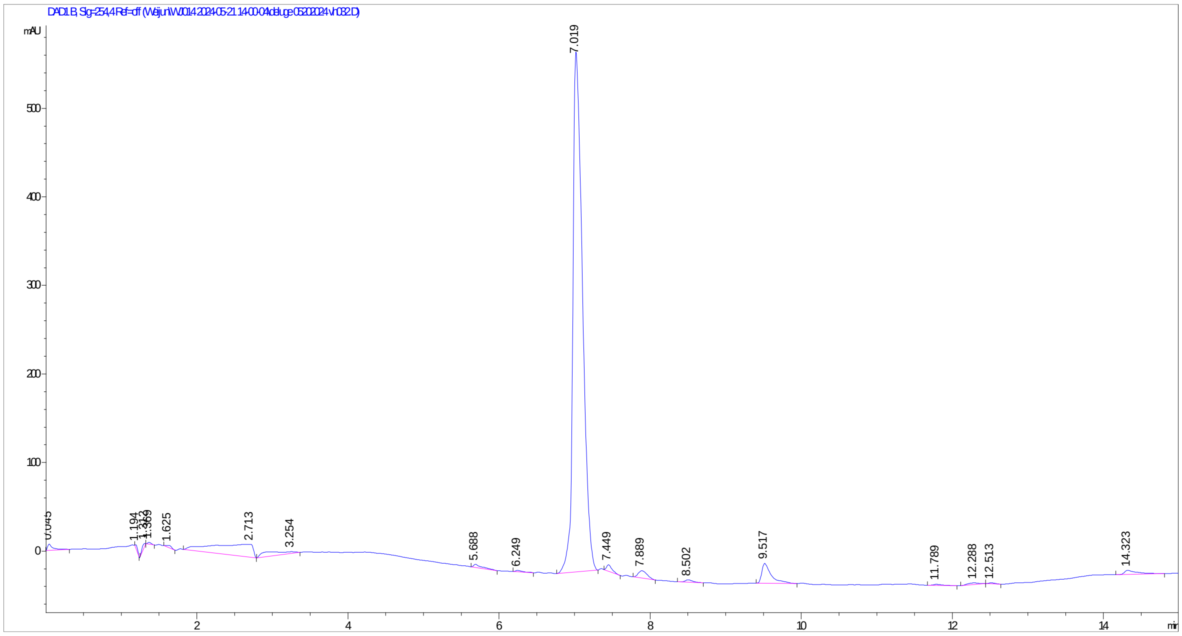

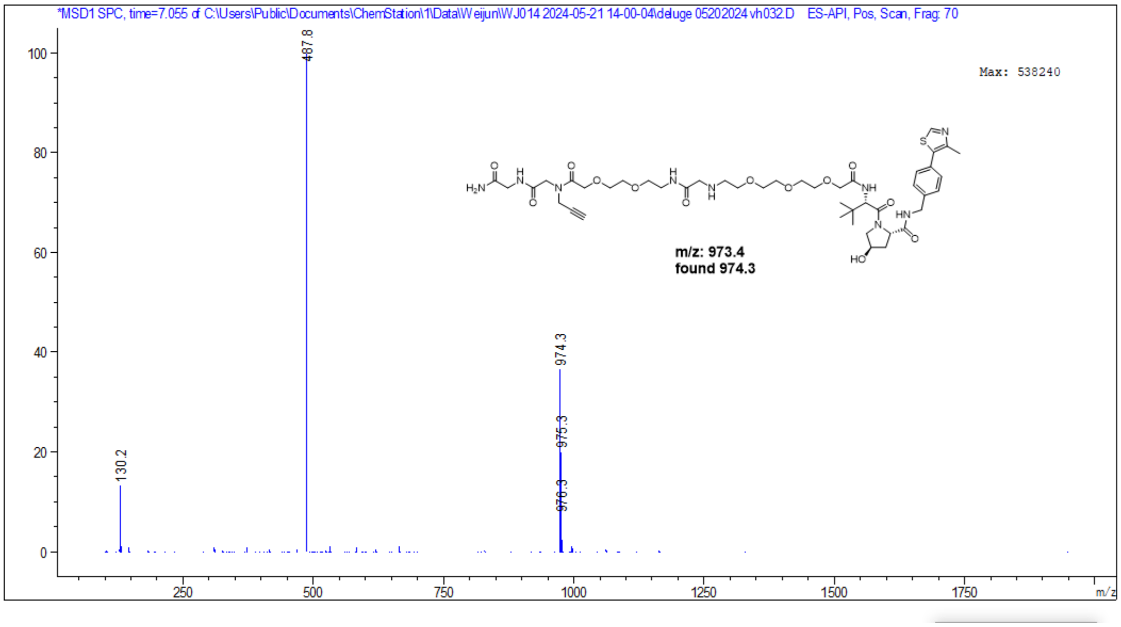
*MSD1 SPC, time=7.055 of C:\Users\Public\Documents\ChemStation\1\Data\Weijun\WJ014 2024-05-21 14-00-04\deluge 05202024 vh032.D ES-API, Pos, Scan, Frag: 70
Max: 538240
487.8
974.3
975.3
976.3
130.2
m/z: 973.4
found 974.3
m/z

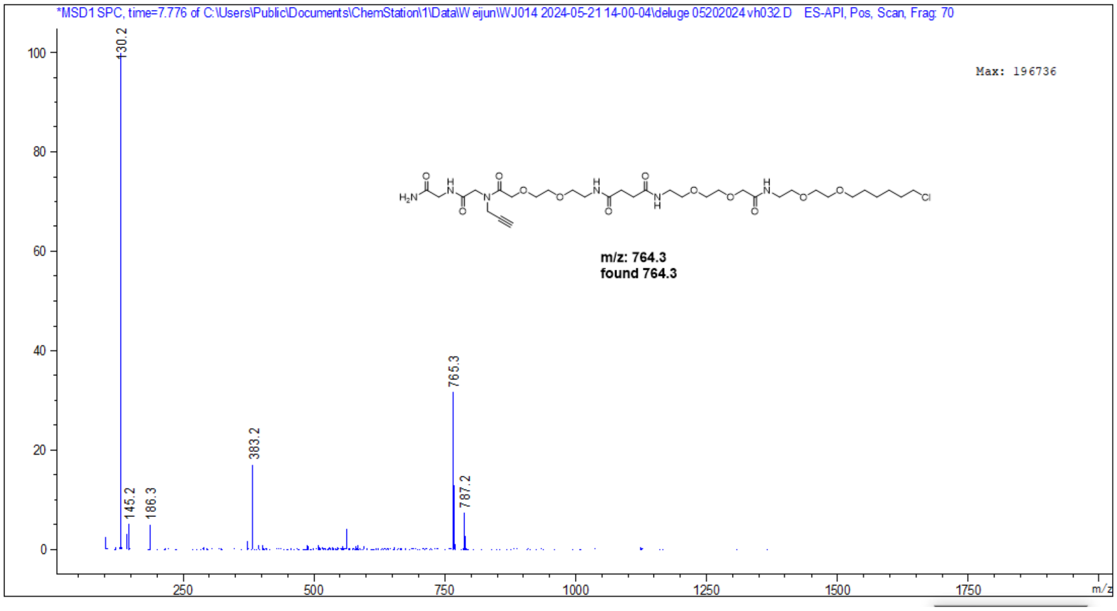
*MSD1 SPC, time=7.776 of C:\Users\Public\Documents\ChemStation\1\Data\Weijun\WJ014 2024-05-21 14-00-04\deluge 05202024 vh032.D ES-API, Pos, Scan, Frag: 70
Max: 196736
130.2
145.2
186.3
383.2
765.3
787.2
m/z: 764.3
found 764.3
m/z

**LC/MS: TG-MZ1**

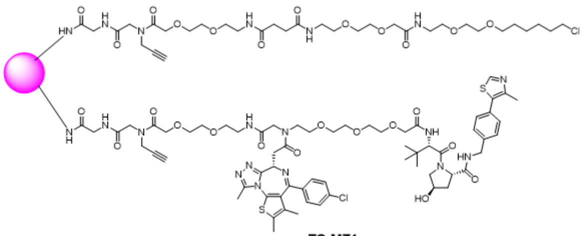


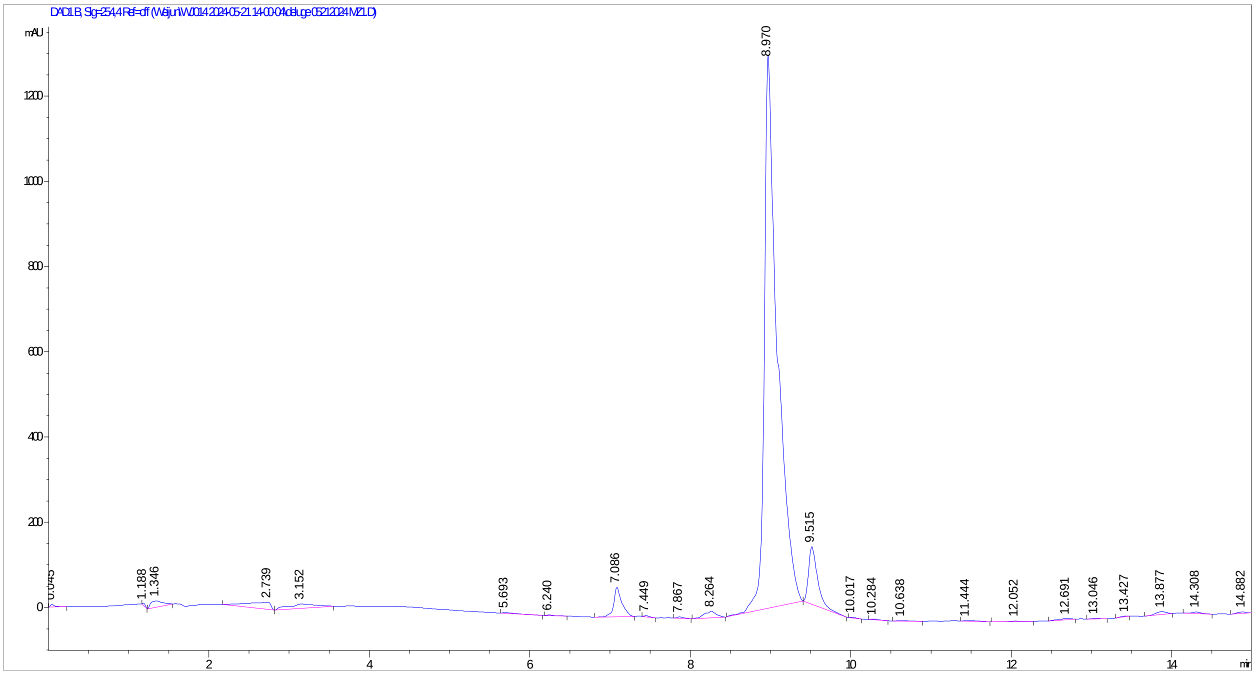

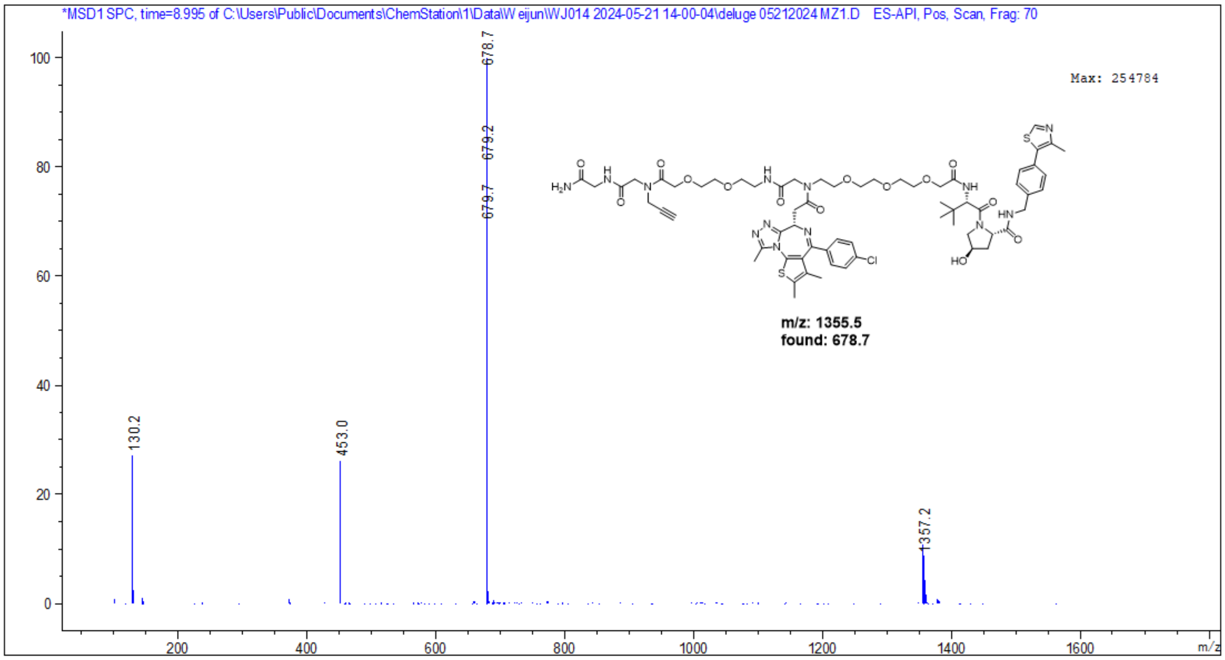
*MSD1 SPC, time=8.995 of C:\Users\Public\Documents\ChemStation\1\Data\Weijun\WJ014 2024-05-21 14-00-04\deluge 05212024 MZ1.D ES-API, Pos, Scan, Frag: 70
Max: 254784
678.7
679.2
679.7
130.2
453.0
1357.2
m/z: 1355.5
found: 678.7
m/z

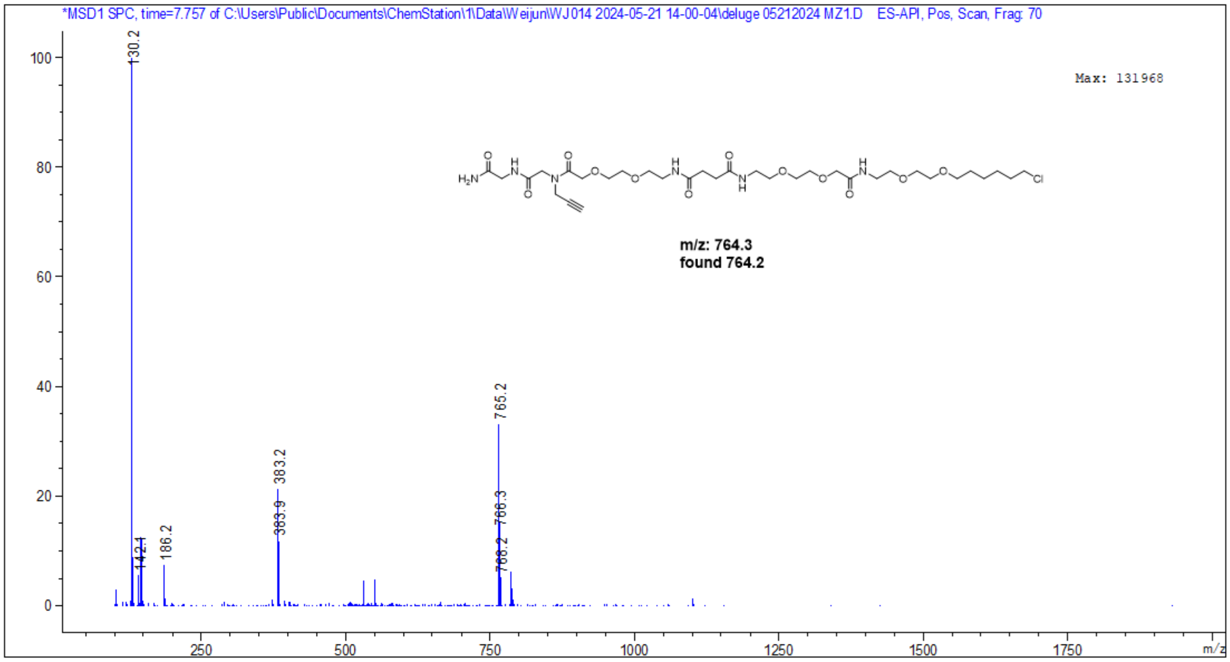
*MSD1 SPC, time=7.757 of C:\Users\Public\Documents\ChemStation\1\Data\Weijun\WJ014 2024-05-21 14-00-04\deluge 05212024 MZ1.D ES-API, Pos, Scan, Frag: 70
Max: 131968
130.2
142.1
186.2
383.2
383.9
765.2
766.3
768.2
m/z: 764.3
found 764.2
m/z

**LC/MS: TG-Pom**

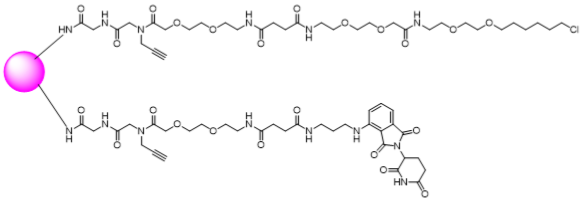


TG-Pom.

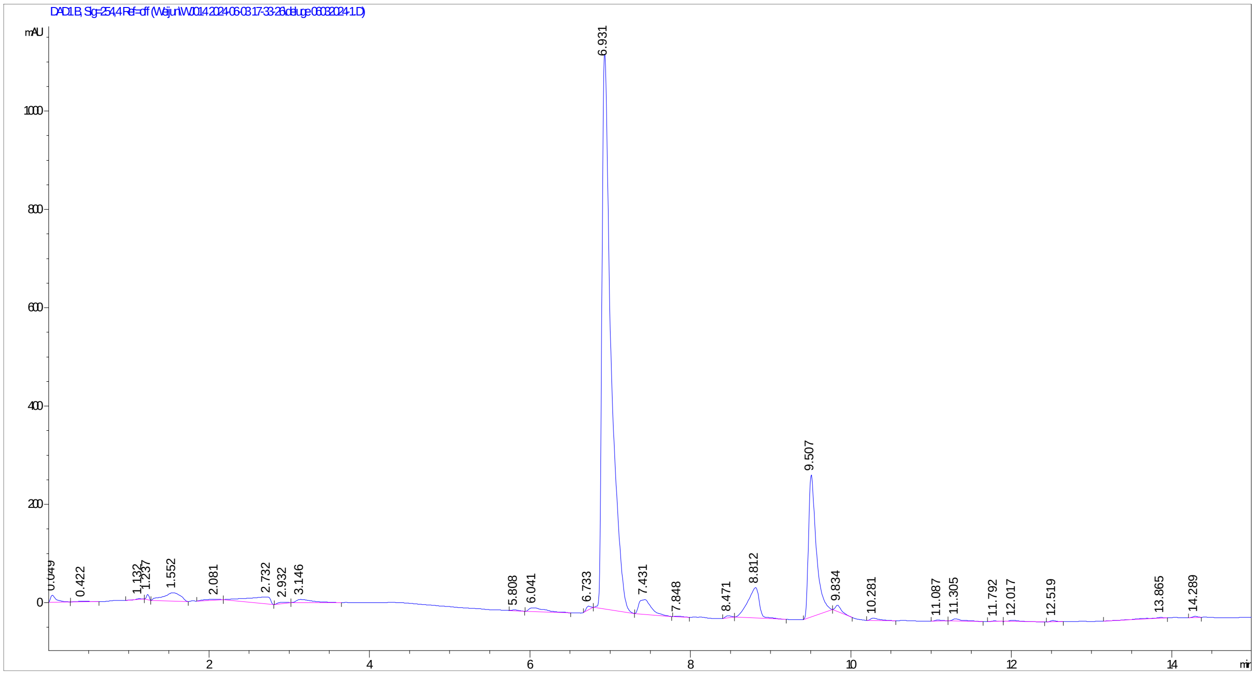

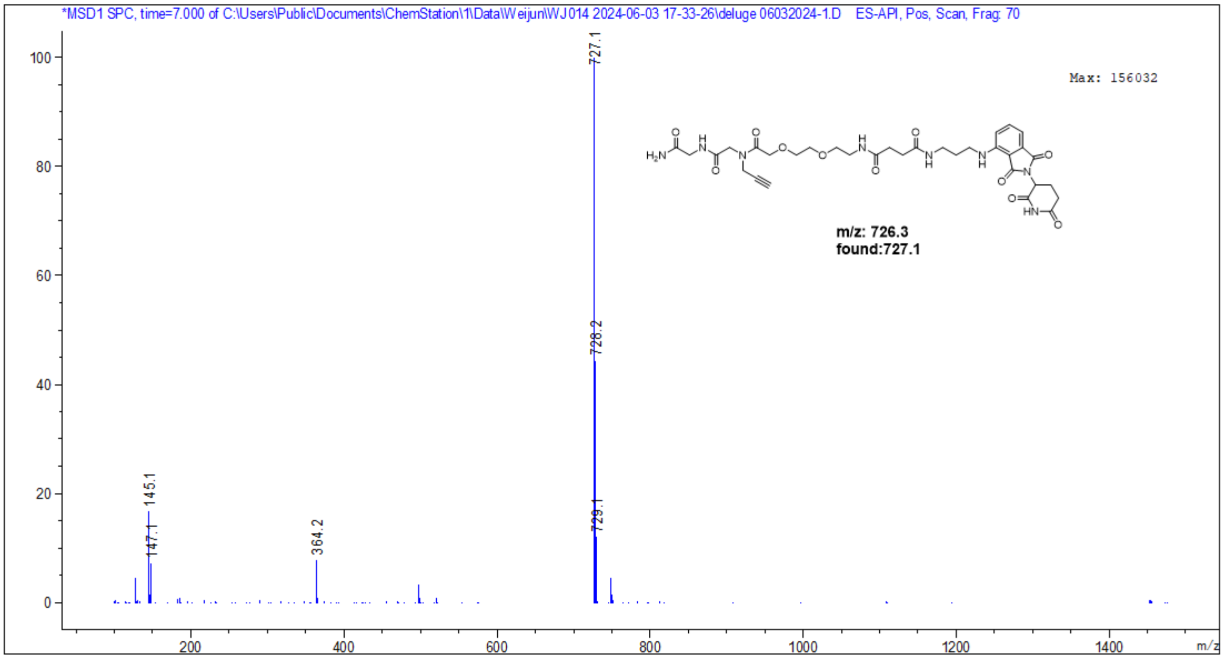
*MSD1 SPC, time=7.000 of C:\Users\Public\Documents\ChemStation\1\Data\Weijun\WJ014 2024-06-03 17-33-26\deluge 06032024-1.D ES-API, Pos, Scan, Frag: 70
Max: 156032
727.1
728.2
729.1
145.1
147.1
364.2
m/z: 726.3
found:727.1
m/z

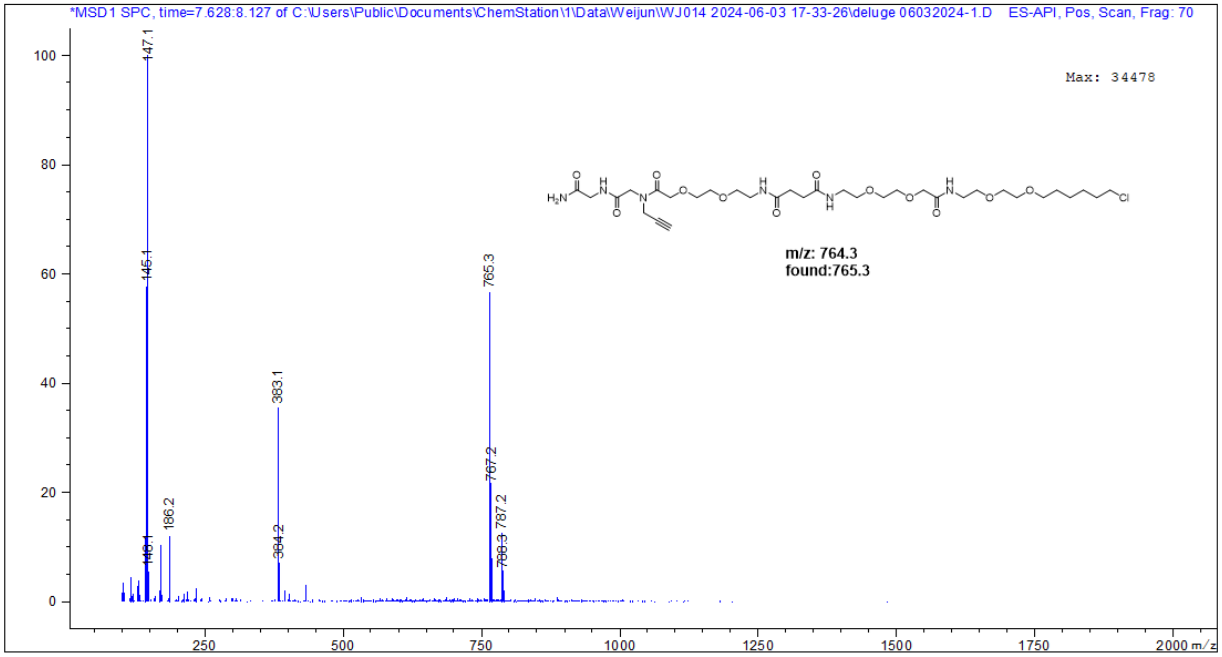
*MSD1 SPC, time=7.628:8.127 of C:\Users\Public\Documents\ChemStation\1\Data\Weijun\WJ014 2024-06-03 17-33-26\deluge 06032024-1.D ES-API, Pos, Scan, Frag: 70
Max: 34478
147.1
145.1
186.2
383.1
384.2
765.3
767.2
787.2
m/z: 764.3
found:765.3
m/z

**LC/MS: TG-mePom**

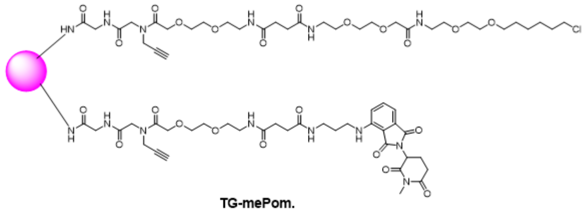

TG-mePom.

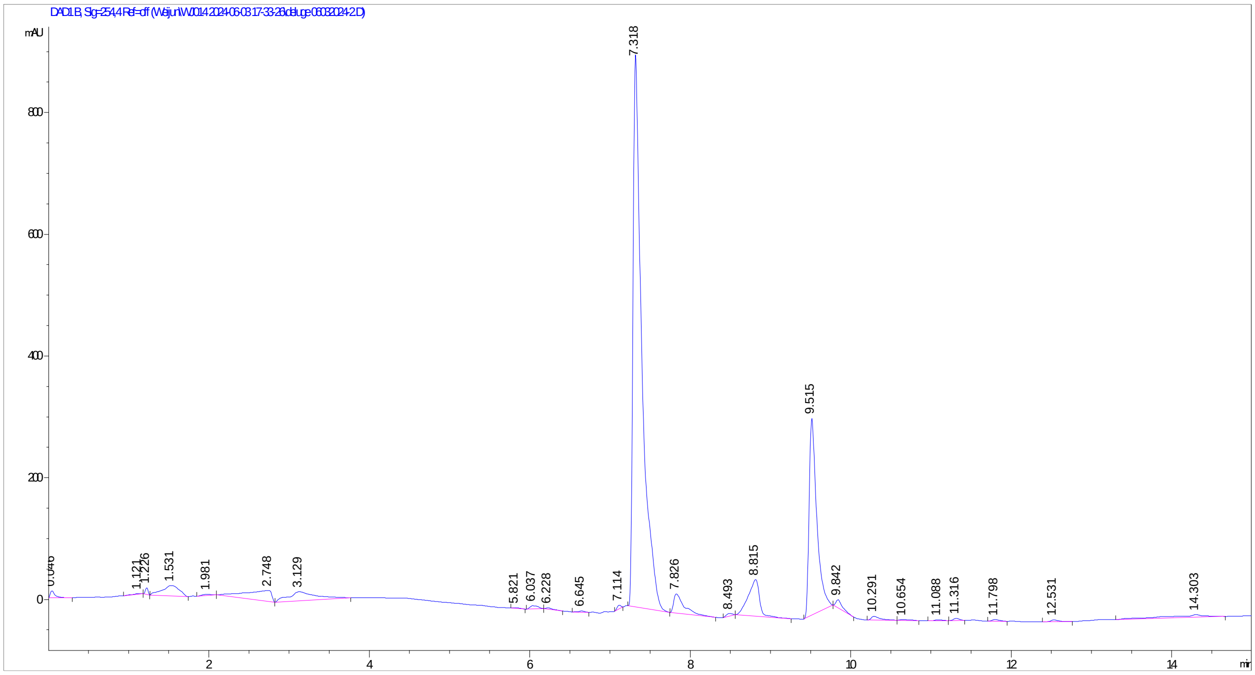

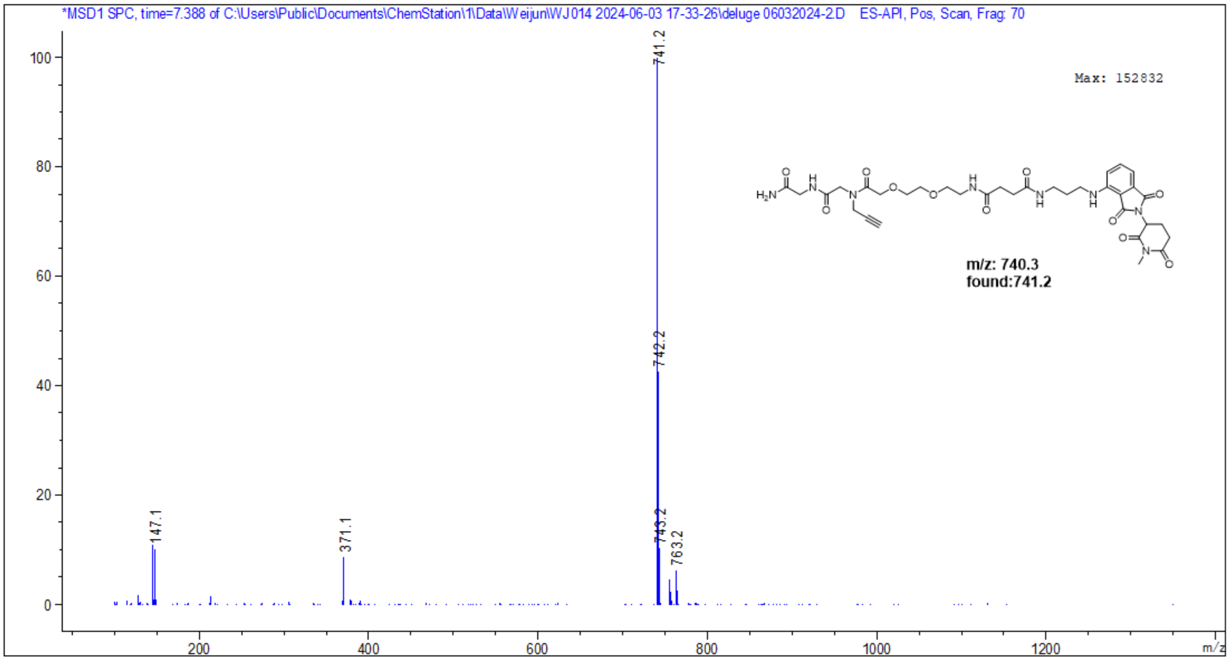
*MSD1 SPC, time=7.388 of C:\Users\Public\Documents\ChemStation\1\Data\Weijun\WJ014 2024-06-03 17-33-26\deluge 06032024-2.D ES-API, Pos, Scan, Frag: 70
Max: 152832
741.2
742.2
743.2
763.2
147.1
371.1
m/z: 740.3
found:741.2
m/z

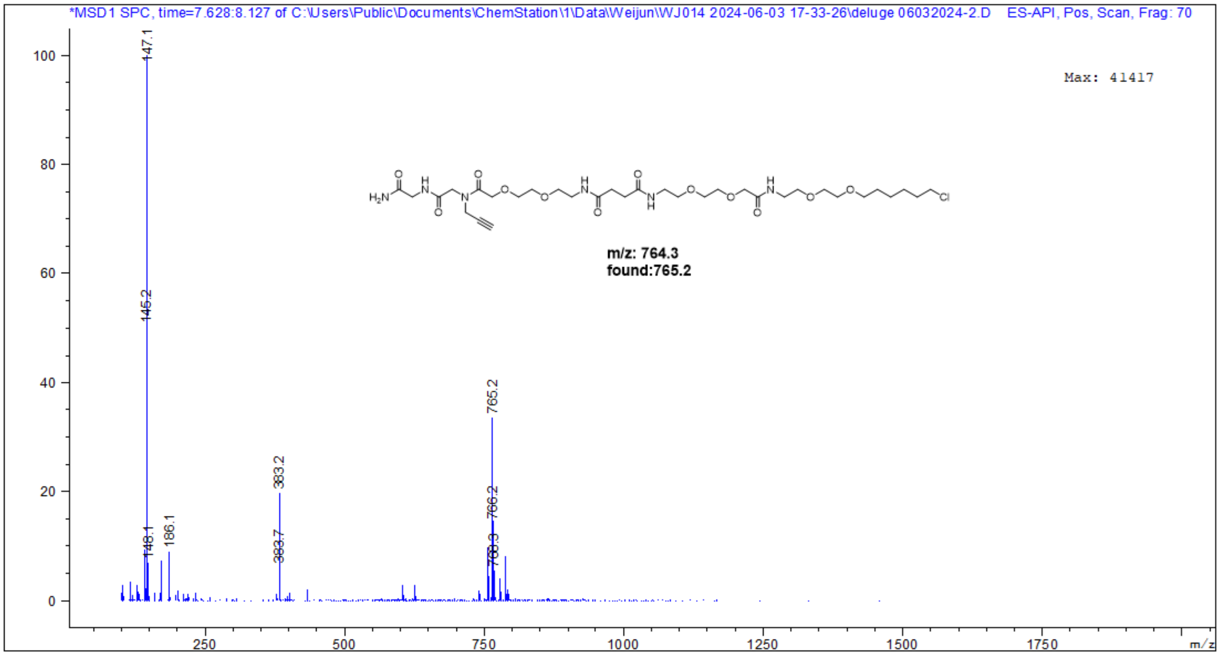
*MSD1 SPC, time=7.628:8.127 of C:\Users\Public\Documents\ChemStation\1\Data\Weijun\WJ014 2024-06-03 17-33-26\deluge 06032024-2.D ES-API, Pos, Scan, Frag: 70
Max: 41417
147.1
145.2
148.1
186.1
383.2
383.7
765.2
766.2
m/z: 764.3
found:765.2
m/z

### LC/MS: TG-Tom

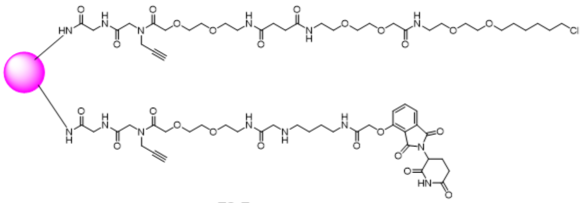


TG-Tom.

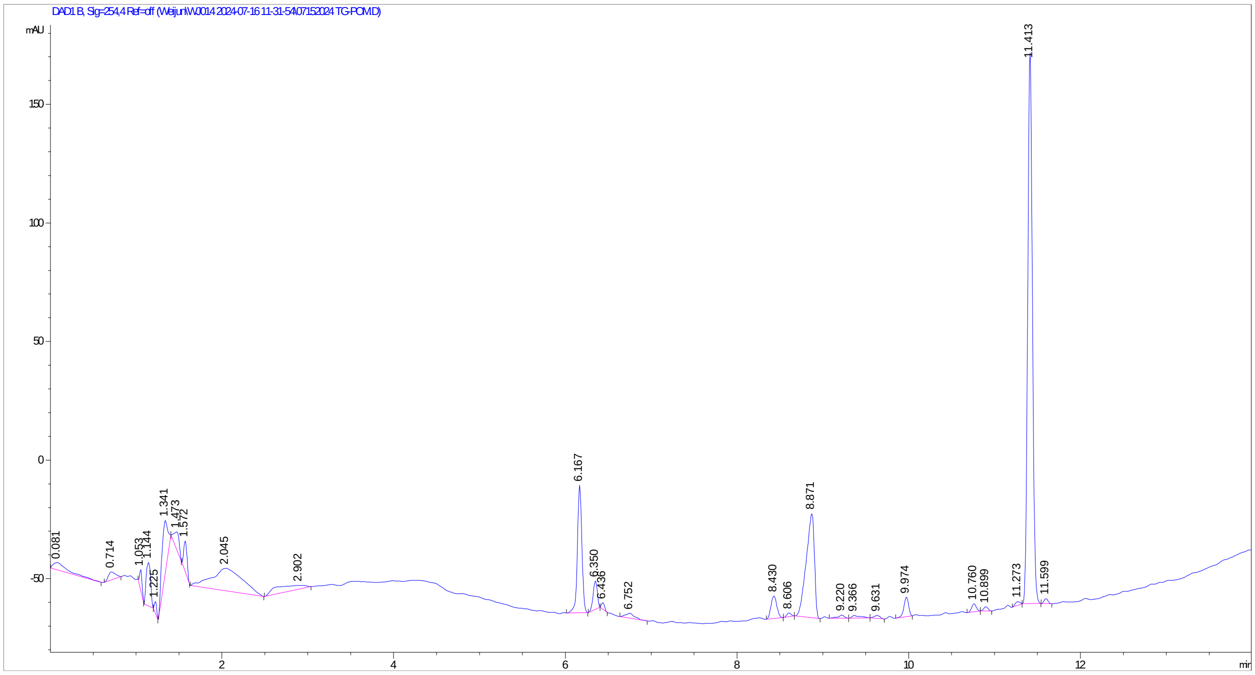

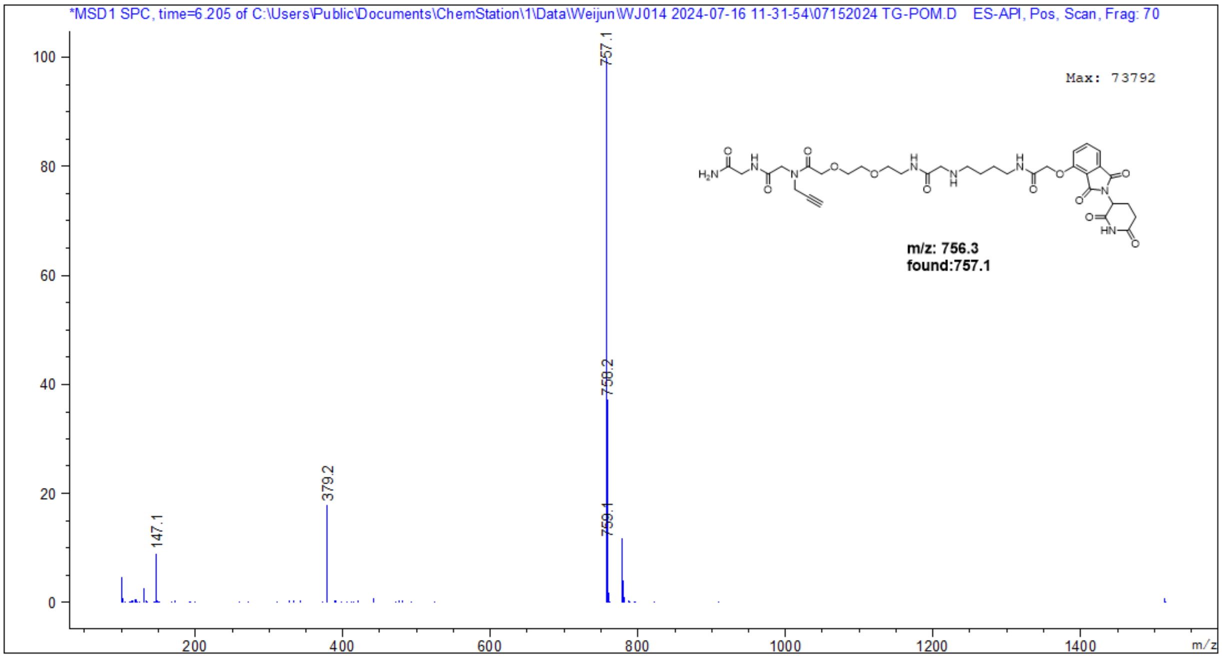
*MSD1 SPC, time=6.205 of C:\Users\Public\Documents\ChemStation\1\Data\Weijun\WJ014 2024-07-16 11-31-54\07152024 TG-POM.D ES-API, Pos, Scan, Frag: 70
Max: 73792
m/z: 756.3
found:757.1
757.1
758.2
759.1
379.2
147.1
m/z

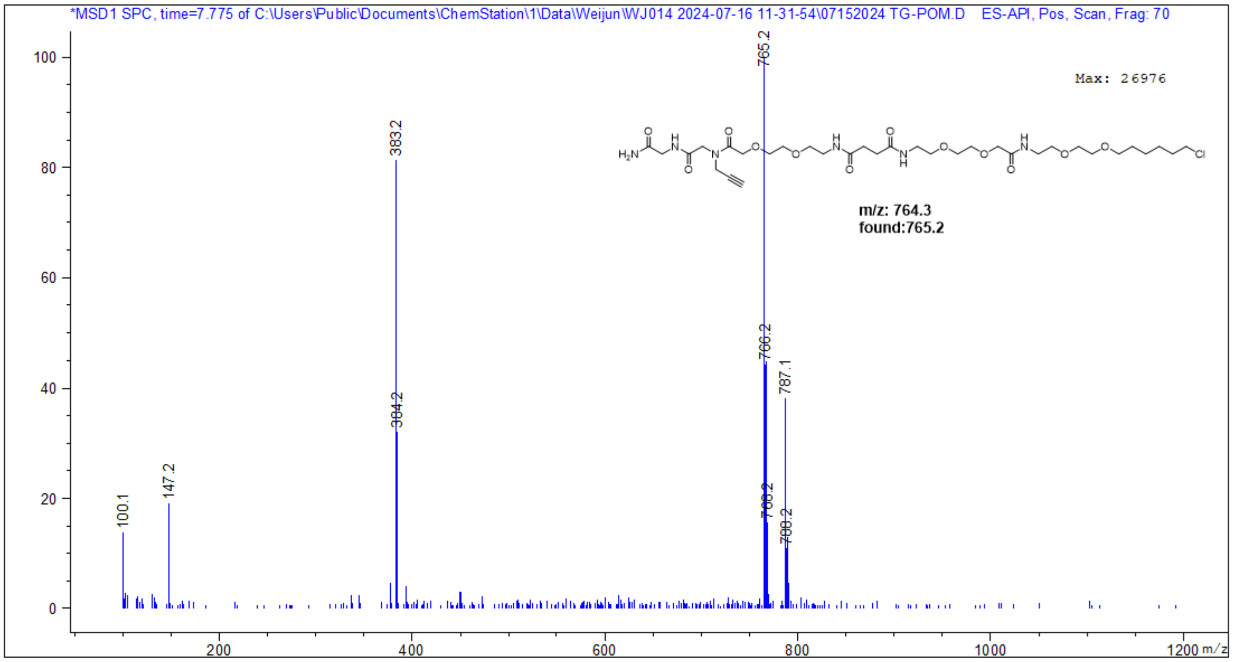
*MSD1 SPC, time=7.775 of C:\Users\Public\Documents\ChemStation\1\Data\Weijun\WJ014 2024-07-16 11-31-54\07152024 TG-POM.D ES-API, Pos, Scan, Frag: 70
Max: 26976
m/z: 764.3
found:765.2
765.2
383.2
766.2
787.1
384.2
147.2
100.1
m/z

### LC/MS: TG-dBET1

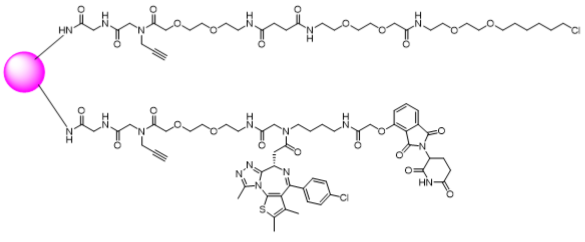


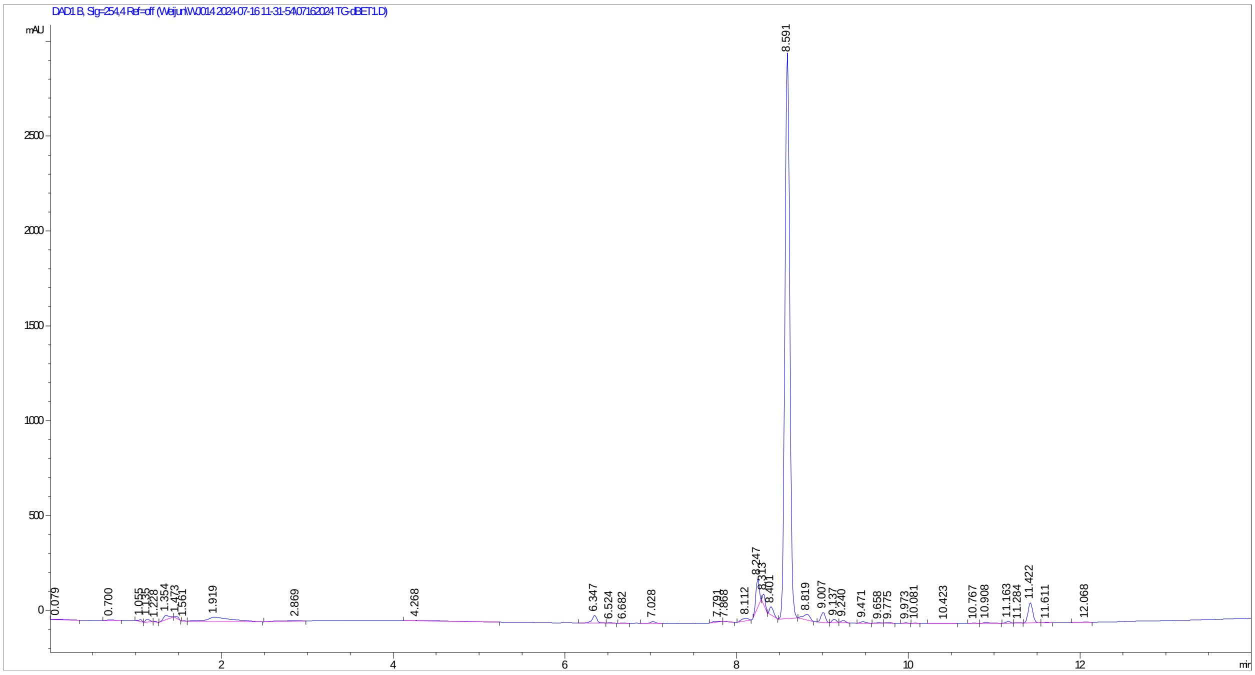

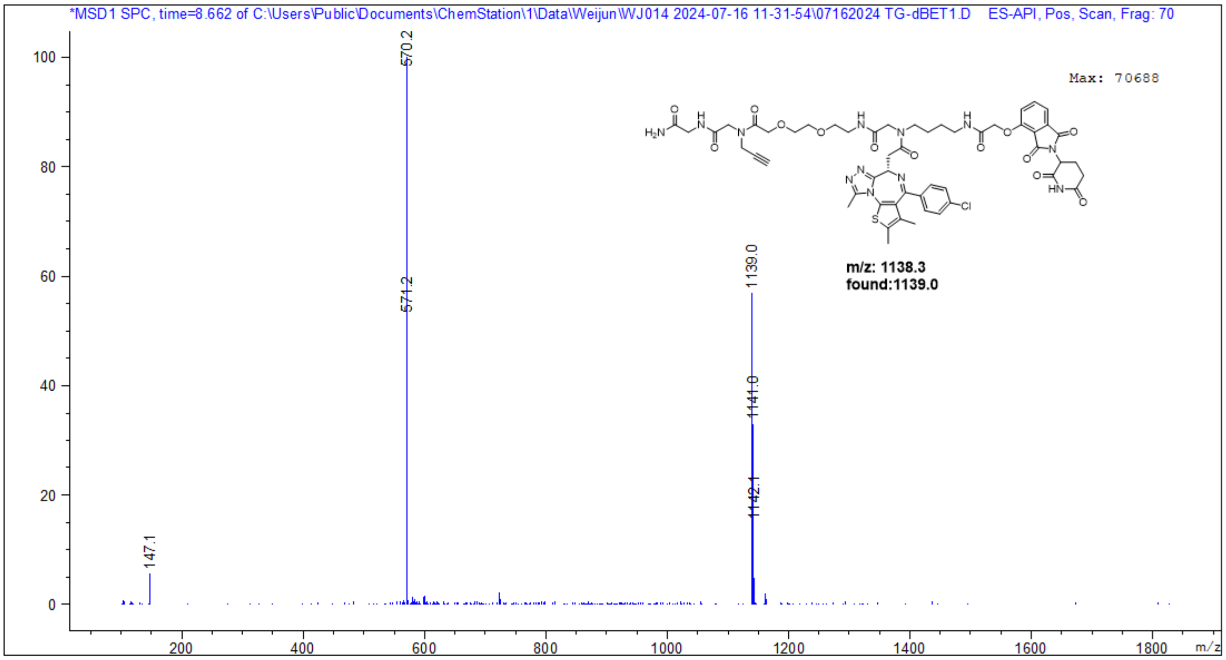
*MSD1 SPC, time=8.662 of C:\Users\Public\Documents\ChemStation\1\Data\Weijun\WJ014 2024-07-16 11-31-54\07162024 TG-dBET1.D ES-API, Pos, Scan, Frag: 70
Max: 70688
570.2
571.2
1139.0
1141.0
1142.1
147.1
m/z: 1138.3
found:1139.0
m/z

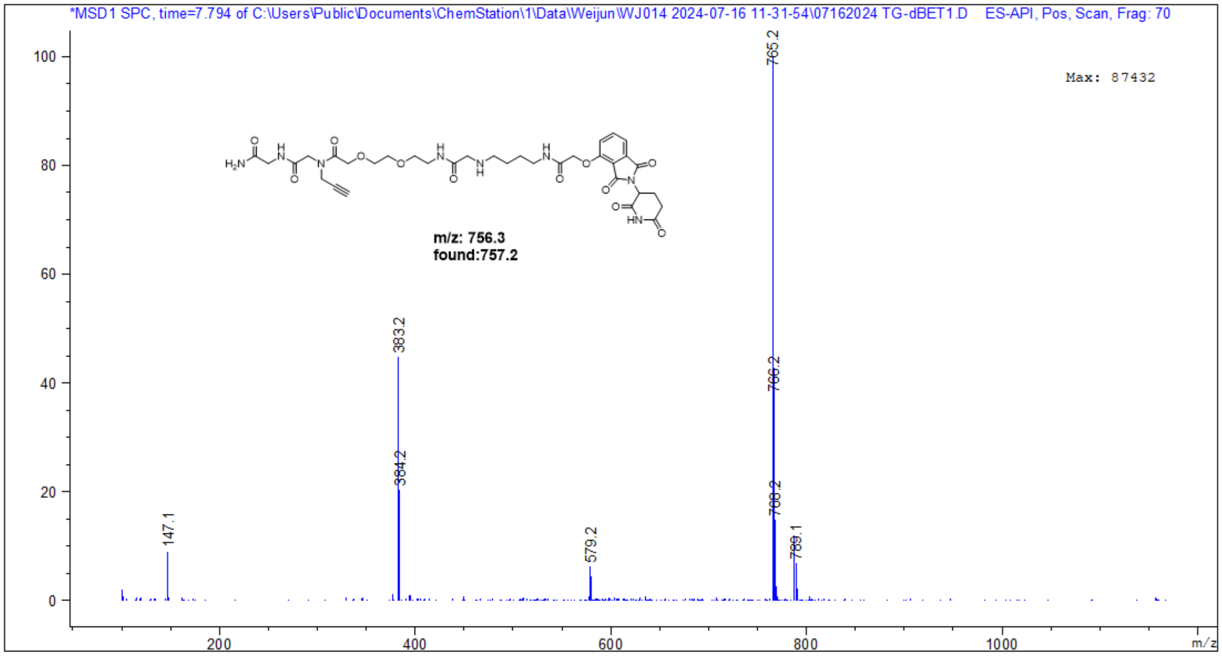
*MSD1 SPC, time=7.794 of C:\Users\Public\Documents\ChemStation\1\Data\Weijun\WJ014 2024-07-16 11-31-54\07162024 TG-dBET1.D ES-API, Pos, Scan, Frag: 70
Max: 87432
765.2
766.2
768.2
383.2
384.2
579.2
147.1
m/z: 756.3
found:757.2
m/z

**WJ3004: (2S,4R)-1-(2-(4-acetylpiperazin-1-yl)acetyl)-N-(2-amino-2-oxoethyl)-4-hydroxypyrrolidine-2-carboxamide.** $^{1}$H NMR (600 MHz, DMSO-$d_6$) δ 10.23 (s, 1H), 8.44 (s, 1H), 7.13 (d, *J* = 2.7 Hz, 2H), 4.50 – 4.36 (m, 3H), 4.33 – 4.15 (m, 4H), 3.73 (dd, *J* = 16.6, 6.1 Hz, 1H), 3.61 (tp, *J* = 16.8, 5.7 Hz, 4H), 3.45 (dd, *J* = 12.0, 4.4 Hz, 3H), 3.33 (dt, *J* = 10.0, 1.8 Hz, 1H), 2.12 – 2.06 (m, 1H), 2.04 (s, 3H), 1.94 (td, *J* = 7.8, 3.9 Hz, 1H). $^{13}$C NMR (151 MHz, DMSO) δ 171.50, 171.41, 171.33, 169.12, 69.04, 67.50, 59.51, 58.65, 54.77, 52.47, 42.36, 38.25, 21.41. MS (ESI, positive) m/z calculated for $C_{15}H_{26}N_5O_5$ [M+H]$^+$ : 356.2, found: 356.2.

**WJ3006: (2S,4R)-N-(2-amino-2-oxoethyl)-1-(2-(4-(2-((S)-4-(4-chlorophenyl)-2,3,9-trimethyl-6H-thieno[3,2-f][1,2,4]triazolo[4,3-a][1,4]diazepin-6-yl)acetyl)piperazin-1-yl)acetyl)-4-hydroxypyrrolidine-2-carboxamide.** $^{1}$H NMR (600 MHz, DMSO-d6) δ 10.31 (s, 1H), 8.45 (t, J = 6.0 Hz, 1H), 7.52 – 7.47 (m, 2H), 7.44 (dd, J = 8.4, 1.5 Hz, 2H), 7.15 (d, J = 3.1 Hz, 2H), 4.62 – 4.55 (m, 1H), 4.53 – 4.22 (m, 6H), 3.79 – 3.55 (m, 6H), 3.48 (dd, J = 12.0, 4.4 Hz, 2H), 3.38 – 3.32 (m, 1H), 2.61 (d, J = 1.6 Hz, 3H), 2.42 (s, 3H), 2.08 (tdd, J = 19.1, 9.2, 4.4 Hz, 1H), 1.95 (ddd, J = 12.6, 7.7, 4.8 Hz, 1H), 1.63 (s, 3H). $^{13}$C NMR (151 MHz, DMSO) δ 171.50, 171.33, 169.12, 164.13, 163.66, 163.56, 158.90, 158.66, 150.44, 137.16, 135.77, 132.62, 131.33, 130.65, 130.41, 130.13, 128.98, 119.05, 117.11, 115.18, 113.25, 69.06, 67.54, 59.53, 56.22, 54.76, 54.47, 52.59, 42.37, 42.01, 38.28, 35.09, 14.48, 13.15, 11.73. MS (ESI, positive) m/z calculated for $C_{32}H_{39}ClN_9O_5S$ [M+H]$^+$ : 696.2, found: 696.3.

**WJ3007: (2S,4R)-N-(2-amino-2-oxoethyl)-1-(2-(4-(3-(2-((S)-4-(4-chlorophenyl)-2,3,9-trimethyl-6H-thieno[3,2-f][1,2,4]triazolo[4,3-a][1,4]diazepin-6-yl)acetamido)propanoyl)piperazin-1-yl)acetyl)-4-hydroxypyrrolidine-2-carboxamide.** $^{1}$H NMR (600 MHz, DMSO-d6) δ 10.27 (s, 1H), 8.45 (t, J = 5.9 Hz, 1H), 8.27 (t, J = 5.7 Hz, 1H), 7.48 (d, J = 8.8 Hz, 2H), 7.43 (d, J = 8.3 Hz, 2H), 7.14 (d, J = 8.5 Hz, 2H), 4.52 (td, J = 7.1, 1.4 Hz, 1H), 4.50 – 4.38 (m, 2H), 4.35 – 4.19 (m, 3H), 4.03 (s, 1H), 3.77 – 3.57 (m, 4H), 3.46 (dd, J = 12.0, 4.4 Hz, 2H), 3.38 – 3.30 (m, 3H), 3.24 (d, J = 7.1 Hz, 2H), 3.07 (d, J = 54.5 Hz, 3H), 2.61 (s, 3H), 2.56 (d, J = 7.0 Hz, 2H), 2.41 (s, 3H), 2.12 – 2.00 (m, 1H), 1.94 (ddd, J = 12.7, 7.7, 4.8 Hz, 1H), 1.62 (s, 3H). $^{13}$C NMR (151 MHz, DMSO) δ 171.90, 171.50, 171.34, 170.08, 169.95, 164.07, 163.66, 163.62, 158.95, 158.71, 155.55, 150.45, 137.15, 135.77, 132.65, 131.32, 130.68, 130.38, 130.11, 128.96, 119.10, 117.16, 115.23, 113.30, 69.04, 67.51, 59.52, 58.65, 56.25, 54.74, 54.20, 52.46, 42.37, 38.26, 37.94, 35.38, 32.80, 14.50, 13.13, 11.73. MS (ESI, positive) m/z calculated for $C_{35}H_{44}ClN_{10}O_6S$ [M+H]$^+$ : 767.3, found: 767.3.

**WJ3008: (2S,4R)-N-(2-amino-2-oxoethyl)-1-(2-(4-(4-(2-((S)-4-(4-chlorophenyl)-3,9-dimethyl-6H-thieno[3,2-f][1,2,4]triazolo[4,3-a][1,4]diazepin-6-yl)acetamido)butanoyl)piperazin-1-yl)acetyl)-4-hydroxypyrrolidine-2-carboxamide.** $^{1}$H NMR (600 MHz, DMSO-d6) δ 10.23 (s, 1H), 8.44 (t, J = 5.9 Hz, 1H), 8.22 (t, J = 5.7 Hz, 1H), 7.49 (d, J = 8.6 Hz, 2H), 7.43 (d, J = 8.2 Hz, 2H), 7.14 (d, J = 4.3 Hz, 2H), 4.51 (t, J = 7.3 Hz, 1H), 4.46 – 4.39 (m, 2H), 4.35 – 4.18 (m, 3H), 4.01 (s, 1H), 3.70 – 3.57 (m, 3H), 3.47 (dt, J = 12.0, 6.1 Hz, 3H), 3.36 – 3.30 (m, 1H), 3.29 – 2.98 (m, 7H), 2.60 (s, 3H), 2.41 (s, 5H), 2.12 – 2.01 (m, 1H), 1.94 (ddd, J = 12.7, 7.7, 4.8 Hz, 1H), 1.69 (d, J = 8.4 Hz, 2H), 1.62 (s, 3H). $^{13}$C NMR (151 MHz, DMSO) δ 171.49, 171.32, 171.21, 169.92, 164.10, 163.68, 163.61, 159.13, 158.89, 158.65, 158.41, 155.58, 150.44, 137.19, 135.76, 132.70, 131.30, 130.66, 130.37, 130.06, 128.98, 119.04, 117.11, 115.17, 113.24, 69.05, 67.51, 59.51, 58.66, 56.24, 55.45, 54.76, 54.32, 52.50, 42.36, 38.46, 38.27, 38.12, 29.73, 25.26, 25.21, 14.52, 13.15, 11.76. MS (ESI, positive) m/z calculated for $C_{36}H_{46}ClN_{10}O_6S$ [M+H]$^+$ : 781.3, found: 781.3.

**WJ3009: (2S,4R)-N-(2-amino-2-oxoethyl)-1-(2-(4-(5-(2-((S)-4-(4-chlorophenyl)-2,3,9-trimethyl-6H-thieno[3,2-f][1,2,4]triazolo[4,3-a][1,4]diazepin-6-yl)acetamido)pentanoyl)piperazin-1-yl)acetyl)-4-hydroxypyrrolidine-2-carboxamide.** $^{1}$H NMR (600 MHz, DMSO-d6) δ 10.25 (s, 1H), 8.44 (s, 1H), 8.21 (s, 1H), 7.49 (d, J = 8.8 Hz, 2H), 7.42 (d, J = 8.5 Hz, 2H), 7.14 (d, J = 3.1 Hz, 2H), 4.54 – 4.47 (m, 1H), 4.47 – 4.38 (m, 2H), 4.36 – 4.17 (m, 3H), 4.06 (s, 1H), 3.75 – 3.56 (m, 4H), 3.47 (td, J = 11.9, 10.9, 4.3 Hz, 3H), 3.36 – 3.30 (m, 1H), 3.29 – 3.15 (m, 3H), 3.11 (tt, J = 13.4, 6.9 Hz, 3H), 2.60 (s, 3H), 2.41 (s, 3H), 2.38 (s, 2H), 2.12 – 2.01 (m, 1H), 1.94 (ddd, J = 12.7, 7.8, 4.7 Hz, 1H), 1.62 (s, 3H), 1.53 (d, J = 6.8 Hz, 2H), 1.50 – 1.42 (m, 2H). $^{13}$C NMR (151 MHz, DMSO) δ 171.49, 171.38, 171.33, 169.82, 163.58, 159.10, 158.86, 158.63, 158.39, 155.58, 150.39, 137.18, 135.73, 132.70, 131.27, 130.64, 130.34, 130.08, 128.96, 119.26, 117.32, 115.38, 113.44, 69.04, 67.50, 59.51, 56.23, 54.76, 54.31, 42.37, 42.01, 38.78, 38.26, 38.07, 32.10, 29.31, 22.49, 14.52, 13.14, 11.75. MS (ESI, positive) m/z calculated for $C_{37}H_{48}ClN_{10}O_6S$ [M+H]$^+$ : 795.3, found: 795.3.

**WJ3010: (2S,4R)-N-(2-amino-2-oxoethyl)-1-(2-(4-(6-(2-((S)-4-(4-chlorophenyl)-2,3,9-trimethyl-6H-thieno[3,2-f][1,2,4]triazolo[4,3-a][1,4]diazepin-6-yl)acetamido)hexanoyl)piperazin-1-yl)acetyl)-4-hydroxypyrrolidine-2-carboxamide.** $^{1}$H NMR (600 MHz, DMSO-d6) δ 10.25 (s, 1H), 8.45 (s, 1H), 8.18 (d, J = 5.8 Hz, 1H), 7.50 – 7.46 (m, 2H), 7.42 (d, J = 8.4 Hz, 2H), 7.14 (d, J = 3.5 Hz, 2H), 4.51 (dd, J = 7.9, 6.3 Hz, 1H), 4.50 – 4.38 (m, 2H), 4.34 – 4.15 (m, 3H), 4.03 (s, 1H), 3.73 – 3.56 (m, 3H), 3.55 – 3.37 (m, 3H), 3.33 (dd, J = 10.6, 2.1 Hz, 1H), 3.22 (qd, J = 15.0, 7.1 Hz, 2H), 3.10 (q, J = 6.7 Hz, 5H), 2.60 (s, 3H), 2.41 (s, 3H), 2.34 (d, J = 7.7 Hz, 2H), 2.13 – 2.00 (m, 1H), 1.94 (ddd, J = 12.7, 7.8, 4.8 Hz, 1H), 1.62 (s, 3H), 1.48 (dt, J = 31.9, 7.5 Hz, 4H), 1.31 (t, J = 7.7 Hz, 2H). $^{13}$C NMR (151 MHz, DMSO) δ 171.49, 171.43, 171.33, 169.79, 163.57, 158.89, 158.65, 155.57, 150.39, 137.17, 135.77, 132.68, 131.30, 130.62, 130.33, 130.08, 128.96, 119.23, 117.30, 115.36, 113.42, 69.04, 67.50, 59.51, 58.66, 56.22, 54.76, 54.33, 42.37, 38.80, 38.26, 38.07, 32.41, 29.54, 26.55, 24.76, 24.72, 14.52, 13.14, 11.74. MS (ESI, positive) m/z calculated for $C_{38}H_{50}ClN_{10}O_6S$ [M+H]$^+$ : 809.3, found: 809.3.

**WJ3011: (2S,4R)-N-(2-amino-2-oxoethyl)-1-(2-(4-(7-(2-((S)-4-(4-chlorophenyl)-2,3,9-trimethyl-6H-thieno[3,2-f][1,2,4]triazolo[4,3-a][1,4]diazepin-6-yl)acetamido)heptanoyl)piperazin-1-yl)acetyl)-4-hydroxypyrrolidine-2-carboxamide.** $^{1}$H NMR (600 MHz, DMSO-d6) δ 10.27 (s, 1H), 8.46 (s, 1H), 8.18 (d, J = 1.6 Hz, 1H), 7.50 – 7.45 (m, 2H), 7.45 – 7.38 (m, 2H), 7.14 (d, J = 4.4 Hz, 2H), 4.52 (dd, J = 8.1, 6.1 Hz, 1H), 4.49 – 4.38 (m, 2H), 4.36 – 4.18 (m, 3H), 4.04 (s, 1H), 3.71 – 3.58 (m, 3H), 3.46 (dd, J = 12.0, 4.5 Hz, 3H), 3.36 – 3.31 (m, 1H), 3.28 – 3.16 (m, 3H), 3.10 (ddt, J = 21.2, 13.1, 6.2 Hz, 4H), 2.60 (s, 3H), 2.41 (s, 3H), 2.34 (s, 2H), 2.12 – 2.01 (m, 1H), 1.94 (ddd, J = 12.6, 7.7, 4.8 Hz, 1H), 1.62 (s, 3H), 1.46 (dt, J = 28.4, 7.3 Hz, 4H), 1.34 – 1.25 (m, 4H). $^{13}$C NMR (151 MHz, DMSO) δ 171.51, 171.46, 171.36, 169.77, 163.58, 158.95, 158.71, 155.58, 150.41, 137.15, 135.79, 132.66, 131.33, 130.60, 130.34, 130.11, 128.95, 119.24, 117.31, 115.37, 113.43, 69.05, 67.51, 59.53, 56.22, 54.76, 54.32, 42.38, 38.92, 38.24, 38.05, 32.33, 29.60, 28.94, 26.69, 24.94, 14.50, 13.12, 11.72. MS (ESI, positive) m/z calculated for $C_{39}H_{52}ClN_{10}O_6S$ $[M+H]^+$ : 823.3, found: 823.3.

**WJ3012: (2S,4R)-N-(2-amino-2-oxoethyl)-1-(2-(4-(8-(2-((S)-4-(4-chlorophenyl)-2,3,9-trimethyl-6H-thieno[3,2-f][1,2,4]triazolo[4,3-a][1,4]diazepin-6-yl)acetamido)octanoyl)piperazin-1-yl)acetyl)-4-hydroxypyrrolidine-2-carboxamide.** $^{1}$H NMR (600 MHz, DMSO-d6) δ 10.24 (s, 1H), 8.45 (s, 1H), 8.18 (t, J = 5.7 Hz, 1H), 7.50 – 7.45 (m, 2H), 7.44 – 7.36 (m, 2H), 7.14 (s, 2H), 4.51 (dd, J = 8.1, 6.1 Hz, 1H), 4.49 – 4.38 (m, 2H), 4.36 – 4.15 (m, 3H), 4.03 (s, 1H), 3.69 – 3.57 (m, 3H), 3.46 (dd, J = 12.0, 4.4 Hz, 3H), 3.35 – 3.29 (m, 1H), 3.29 – 3.14 (m, 3H), 3.09 (ddq, J = 19.7, 13.0, 6.5 Hz, 4H), 2.60 (s, 3H), 2.41 (s, 3H), 2.33 (d, J = 7.2 Hz, 2H), 2.13 – 2.00 (m, 1H), 1.94 (ddd, J = 12.7, 7.8, 4.8 Hz, 1H), 1.62 (s, 3H), 1.46 (dt, J = 29.1, 7.0 Hz, 4H), 1.28 (s, 6H). $^{13}$C NMR (151 MHz, DMSO) δ 171.49, 171.46, 171.33, 169.77, 163.63, 163.55, 158.89, 158.65, 155.59, 150.38, 137.16, 135.77, 132.68, 131.30, 130.62, 130.32, 130.09, 128.94, 119.20, 117.26, 115.32, 113.38, 69.04, 67.50, 59.51, 56.22, 54.76, 54.34, 42.37, 38.96, 38.26, 38.06, 32.35, 29.68, 29.20, 29.08, 26.82, 24.97, 24.94, 14.51, 13.14, 11.74. MS (ESI, positive) m/z calculated for $C_{40}H_{54}ClN_{10}O_6S$ $[M+H]^+$ : 837.4, found: 837.4.

**WJ3013: (2S,4R)-N-(2-amino-2-oxoethyl)-1-(2-(4-(2-(2-(2-(2-((S)-4-(4-chlorophenyl)-2,3,9-trimethyl-6H-thieno[3,2-f][1,2,4]triazolo[4,3-a][1,4]diazepin-6-yl)acetamido)ethoxy)ethoxy)acetyl)piperazin-1-yl)acetyl)-4-hydroxypyrrolidine-2-carboxamide.** $^{1}$H NMR (600 MHz, DMSO-d6) δ 10.27 (s, 1H), 8.44 (s, 1H), 8.29 (s, 1H), 7.50 – 7.44 (m, 2H), 7.44 – 7.36 (m, 2H), 7.20 – 7.00 (m, 2H), 4.55 – 4.50 (m, 1H), 4.50 – 4.37 (m, 2H), 4.36 – 4.18 (m, 5H), 4.00 (d, J = 36.4 Hz, 1H), 3.68 – 3.53 (m, 8H), 3.50 – 3.40 (m, 4H), 3.36 – 3.19 (m, 6H), 3.18 – 3.01 (m, 2H), 2.60 (s, 3H), 2.41 (s, 3H), 2.12 – 2.01 (m, 1H), 1.94 (dd, J = 8.0, 4.8 Hz, 1H), 1.62 (s, 3H). $^{13}$C NMR (151 MHz, DMSO) δ 171.48, 171.33, 170.18, 168.16, 163.62, 163.59, 159.10, 158.87, 158.63, 158.39, 155.57, 150.42, 137.18, 135.76, 132.69, 131.30, 130.64, 130.34, 130.08, 128.96, 119.24, 117.30, 115.36, 113.42, 70.36, 69.87, 69.84, 69.62, 69.55, 69.04, 67.50, 59.51, 58.66, 56.27, 54.75, 54.26, 42.36, 42.00, 39.12, 38.26, 37.93, 14.52, 13.14, 11.74. MS (ESI, positive) m/z calculated for $C_{38}H_{50}ClN_{10}O_8S$ $[M+H]^+$ : 841.3, found: 841.3.

**WJ3014: (2S,4R)-N-(2-amino-2-oxoethyl)-1-(2-(4-(1-((S)-4-(4-chlorophenyl)-2,3,9-trimethyl-6H-thieno[3,2-f][1,2,4]triazolo[4,3-a][1,4]diazepin-6-yl)-2-oxo-6,9,12,15-tetraoxa-3-azaoctadecan-18-oyl)piperazin-1-yl)acetyl)-4-hydroxypyrrolidine-2-carboxamide.** $^{1}$H NMR (600 MHz, DMSO-d6) δ 10.23 (s, 1H), 8.43 (s, 1H), 8.28 (s, 1H), 7.49 (d, J = 8.8 Hz, 2H), 7.43 (d, J = 8.5 Hz, 2H), 7.14 (d, J = 3.1 Hz, 2H), 4.52 (dd, J = 8.1, 6.0 Hz, 1H), 4.49 – 4.38 (m, 2H), 4.33 – 4.19 (m, 2H), 4.07 (s, 1H), 3.68 – 3.57 (m, 5H), 3.55 – 3.48 (m, 14H), 3.46 (t, J = 5.9 Hz, 3H), 3.33 – 3.20 (m, 5H), 3.05 (d, J = 32.7 Hz, 2H), 2.60 (s, 5H), 2.41 (s, 3H), 2.08 (d, J = 9.7 Hz, 1H), 1.94 (ddd, J = 12.7, 7.7, 4.8 Hz, 1H), 1.62 (s, 3H). $^{13}$C NMR (151 MHz, DMSO) δ 171.48, 171.31, 170.12, 169.69, 163.61, 163.59, 159.16, 158.92, 158.67, 158.43, 155.55, 150.44, 137.15, 135.77, 132.68, 131.35, 130.67, 130.10, 128.95, 118.92, 116.99, 115.06, 113.13, 70.28, 70.21, 70.18, 70.14, 70.09, 69.67, 69.04, 67.03, 59.50, 56.22, 54.74, 54.24, 42.35, 39.09, 38.26, 37.90, 33.00, 14.52, 13.14, 11.74. MS (ESI, positive) m/z calculated for $C_{43}H_{60}ClN_{10}O_{10}S$ $[M+H]^+$ : 943.4, found: 943.4.

**WJ3015: (2S,4R)-N-(2-amino-2-oxoethyl)-1-(2-(4-(3-(2-(2-(2-((S)-4-(4-chlorophenyl)-2,3,9-trimethyl-6H-thieno[3,2-f][1,2,4]triazolo[4,3-a][1,4]diazepin-6-yl)acetamido)ethoxy)ethoxy)propanoyl)piperazin-1-yl)acetyl)-4-hydroxypyrrolidine-2-carboxamide.** $^{1}$H NMR (600 MHz, DMSO-d6) δ 10.25 (s, 1H), 8.45 (s, 1H), 8.28 (s, 1H), 7.50 – 7.44 (m, 2H), 7.43 (d, J = 8.5 Hz, 2H), 7.14 (d, J = 4.3 Hz, 2H), 4.52 (dd, J = 8.0, 6.1 Hz, 1H), 4.50 – 4.38 (m, 2H), 4.35 – 4.18 (m, 3H), 4.07 (s, 1H), 3.68 – 3.56 (m, 5H), 3.53 (d, J = 3.0 Hz, 5H), 3.45 (td, J = 6.0, 3.2 Hz, 4H), 3.36 – 3.19 (m, 6H), 3.09 (s, 2H), 2.67 – 2.61 (m, 2H), 2.60 (s, 3H), 2.41 (s, 3H), 2.13 – 2.00 (m, 1H), 1.94 (ddd, J = 12.7, 7.7, 4.8 Hz, 1H), 1.62 (s, 3H). $^{13}$C NMR (151 MHz, DMSO) δ 171.49, 171.33, 170.15, 169.70, 163.62, 163.60, 159.16, 158.92, 158.68, 158.44, 155.56, 150.43, 137.16, 135.77, 132.68, 131.33, 130.66, 130.34, 130.09, 128.95, 119.13, 117.20, 115.26, 113.33, 70.09, 70.04, 70.01, 69.63, 69.04, 67.51, 66.99, 66.97, 59.51, 58.66, 56.22, 55.43, 54.75, 54.25, 42.36, 42.00, 39.09, 38.26, 37.91, 33.00, 14.51, 13.14, 11.73. MS (ESI, positive) m/z calculated for $C_{39}H_{52}ClN_{10}O_{10}S$ $[M+H]^+$ : 855.3, found: 855.3.

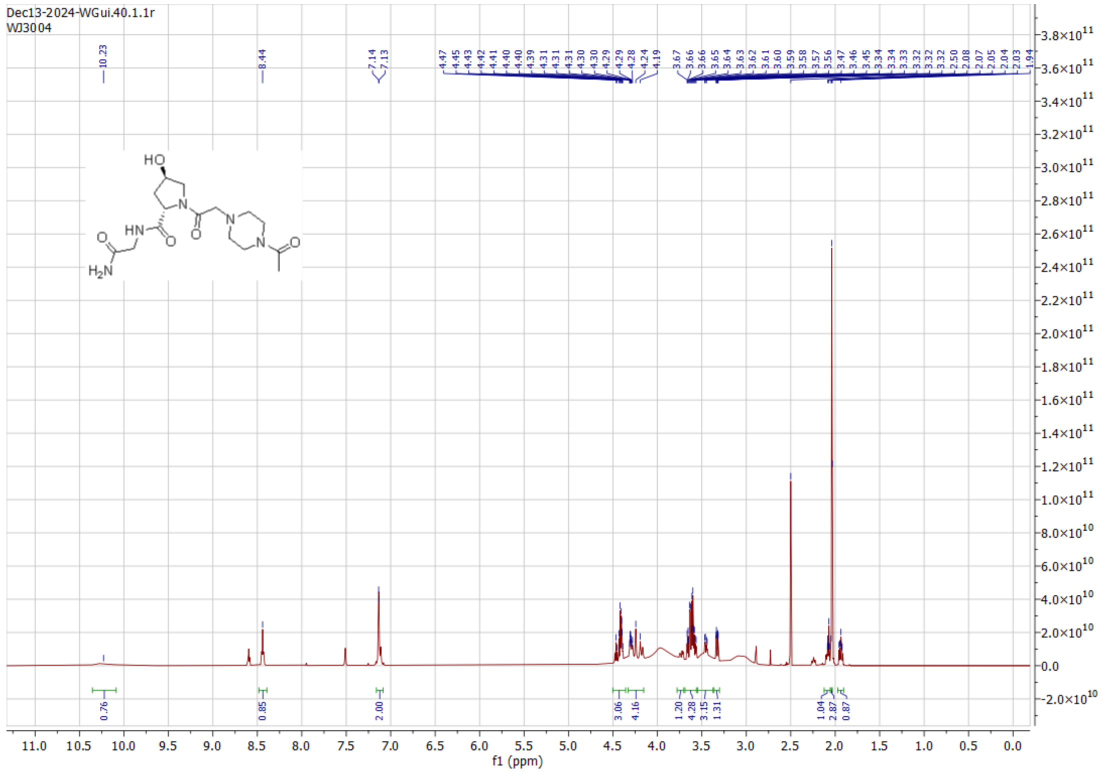
Dec13-2024-WGui.40.1.1r
WJ3004
f1 (ppm)

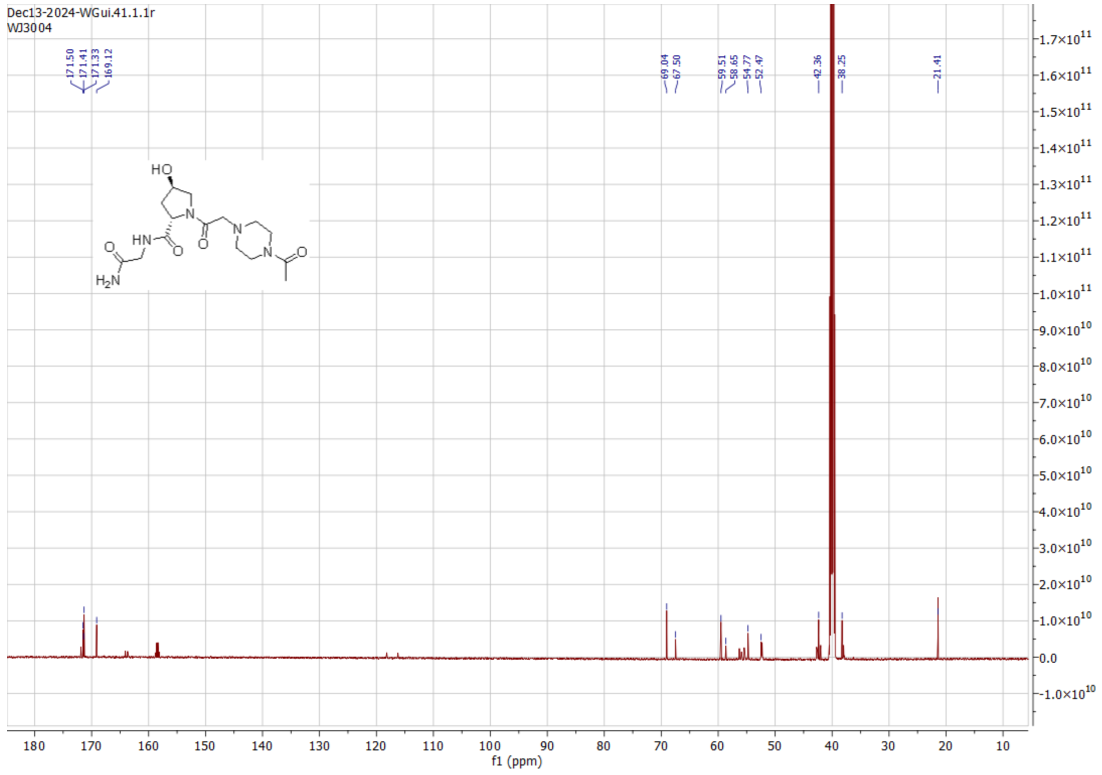
Dec13-2024-WGui.41.1.1r
WJ3004
f1 (ppm)

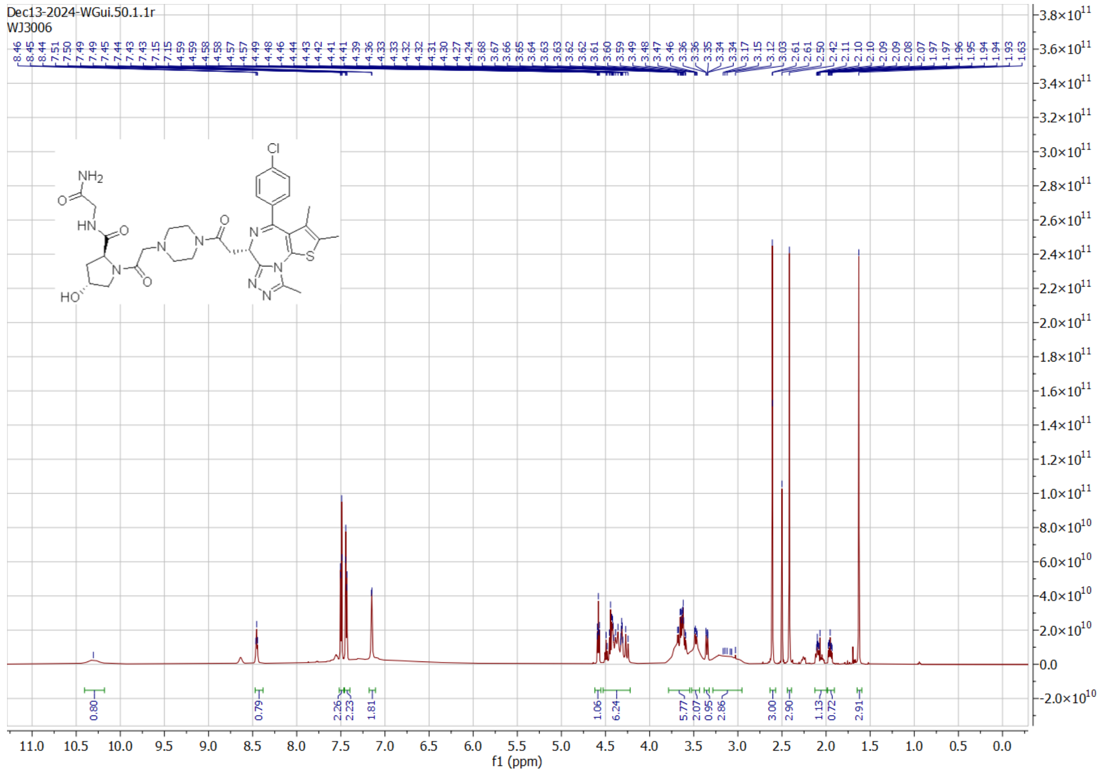
Dec13-2024-WGui.50.1.1r
WJ3006
f1 (ppm)

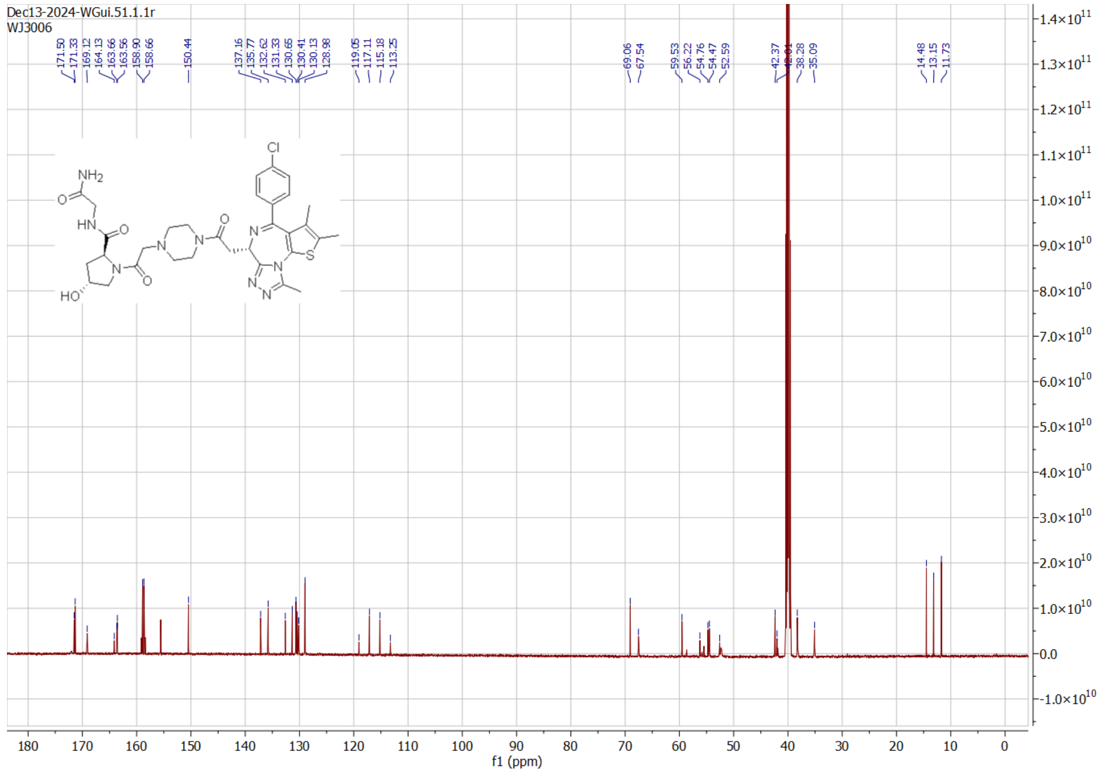
Dec13-2024-WGui.51.1.1r
WJ3006
f1 (ppm)

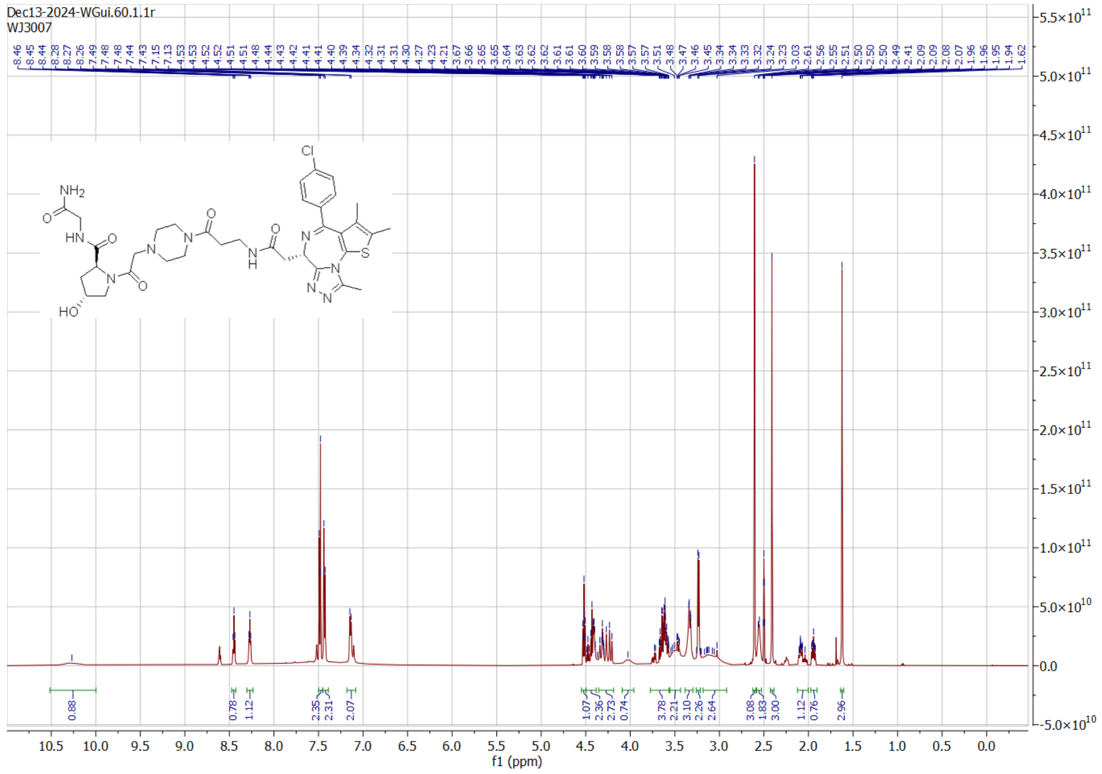
Dec13-2024-WGui.60.1.1r
WJ3007
f1 (ppm)

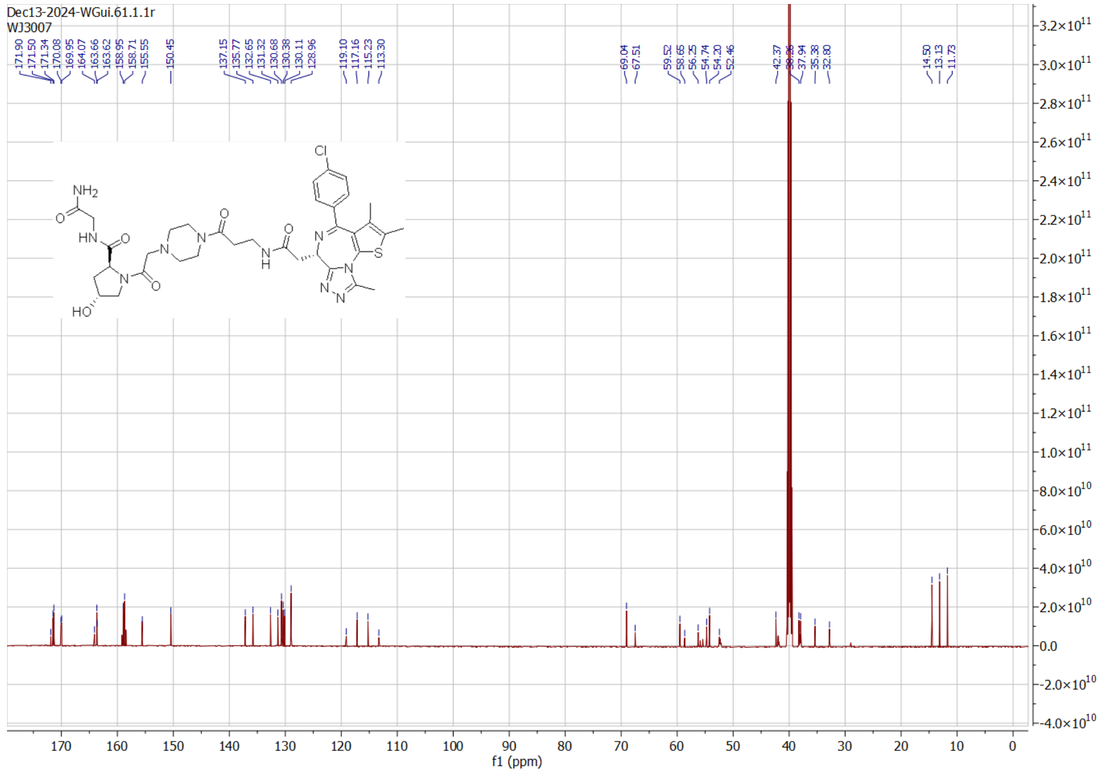
Dec13-2024-WGui.61.1.1r
WJ3007
f1 (ppm)

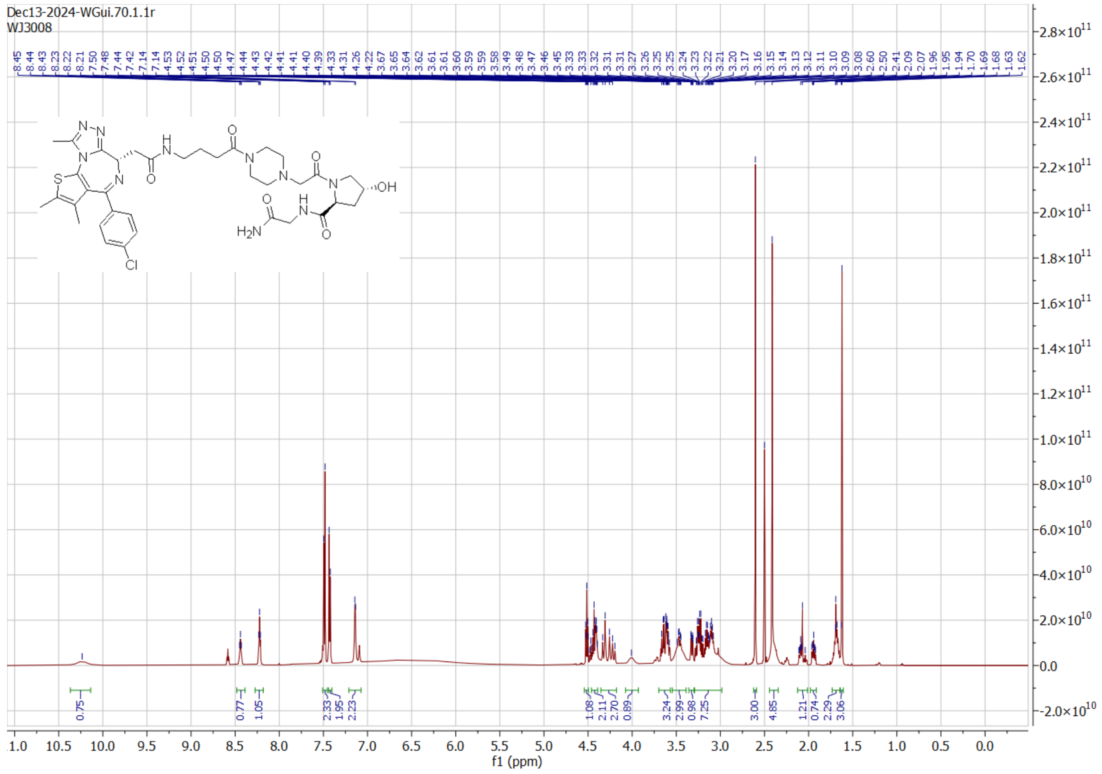

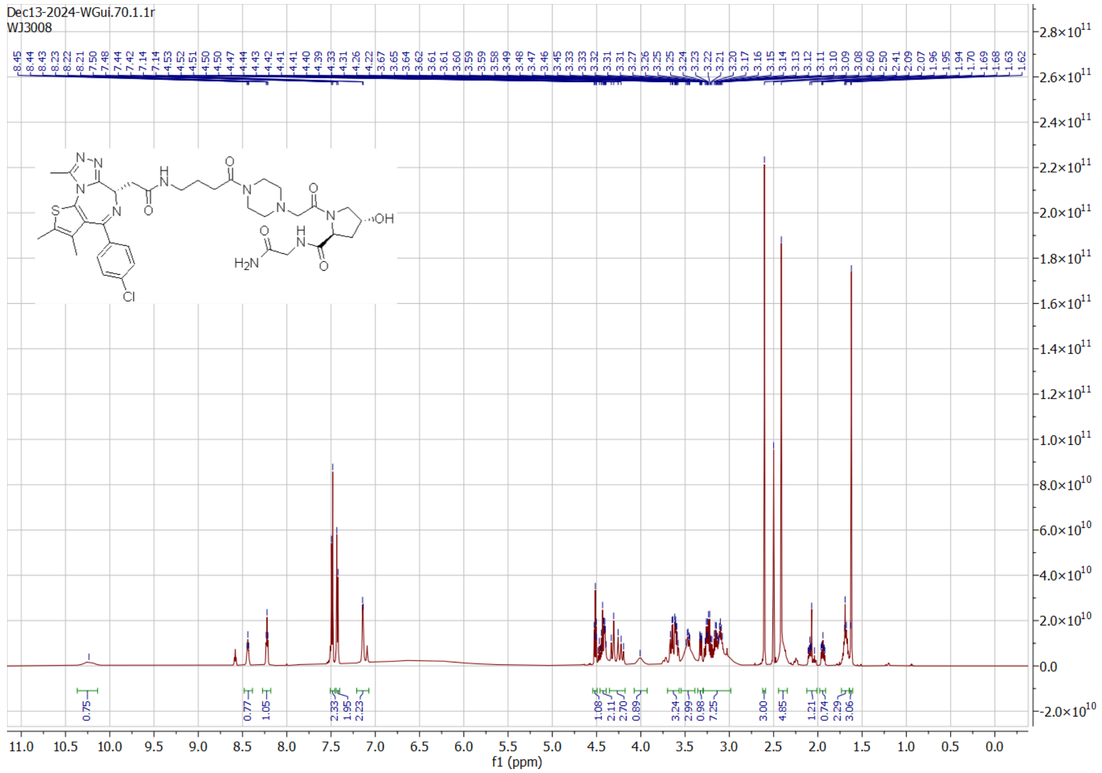

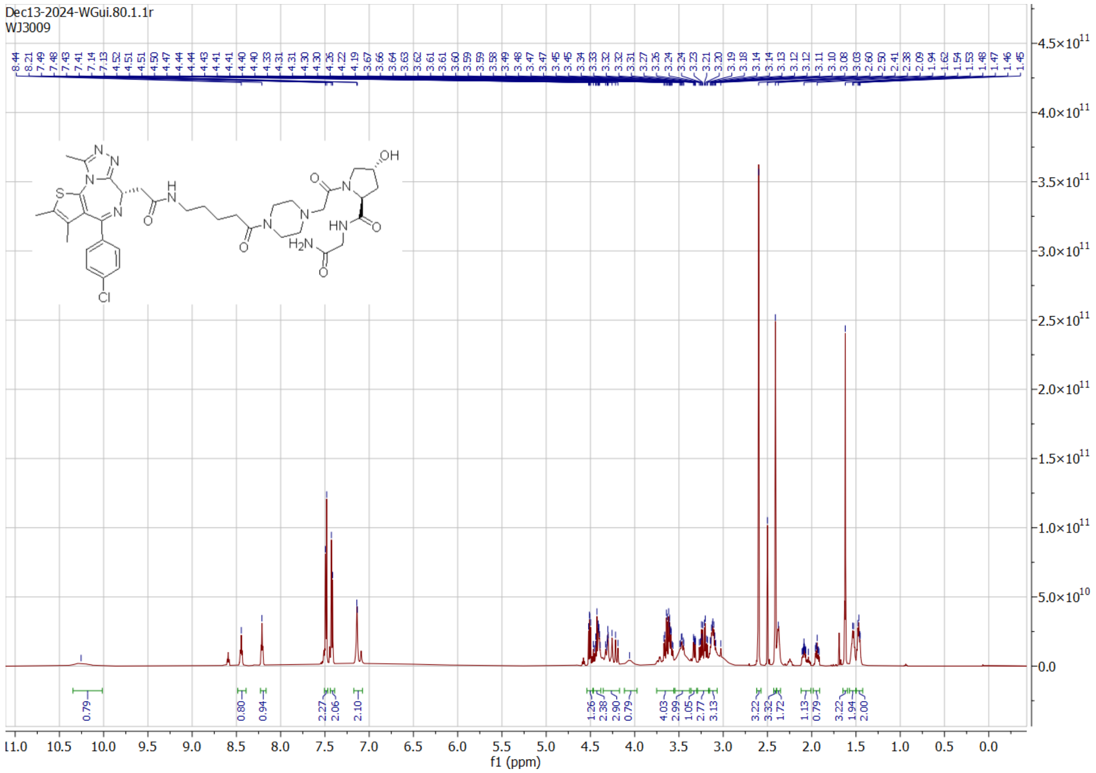

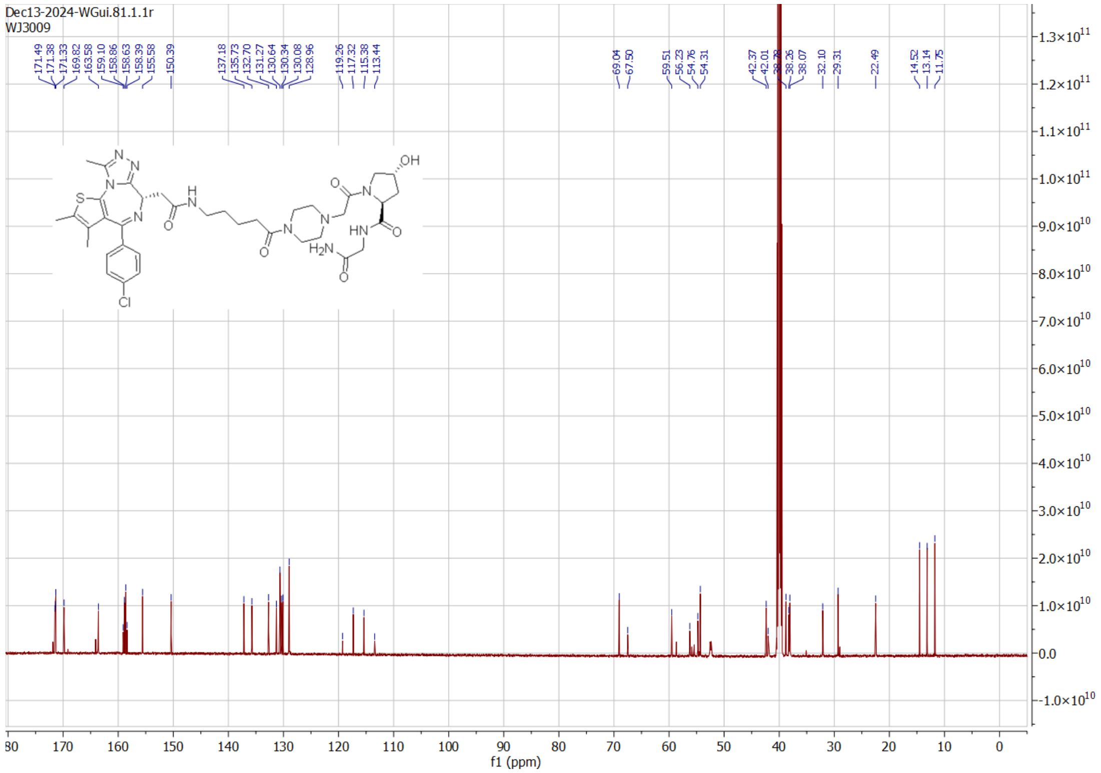

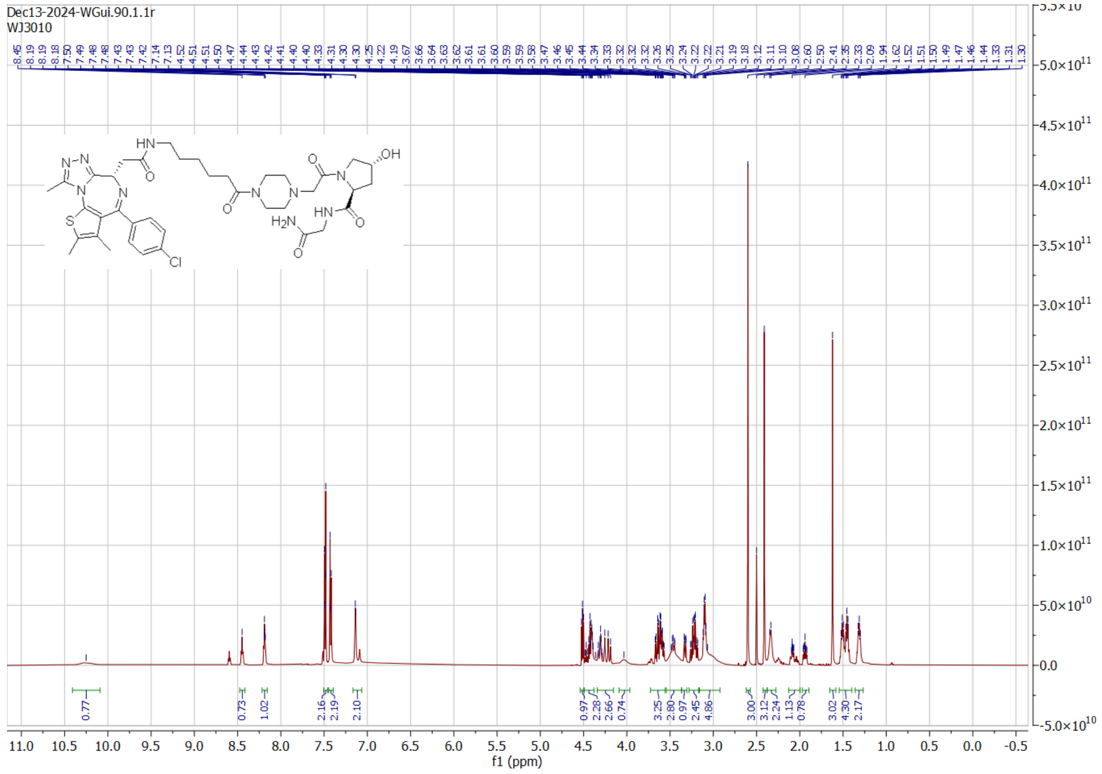

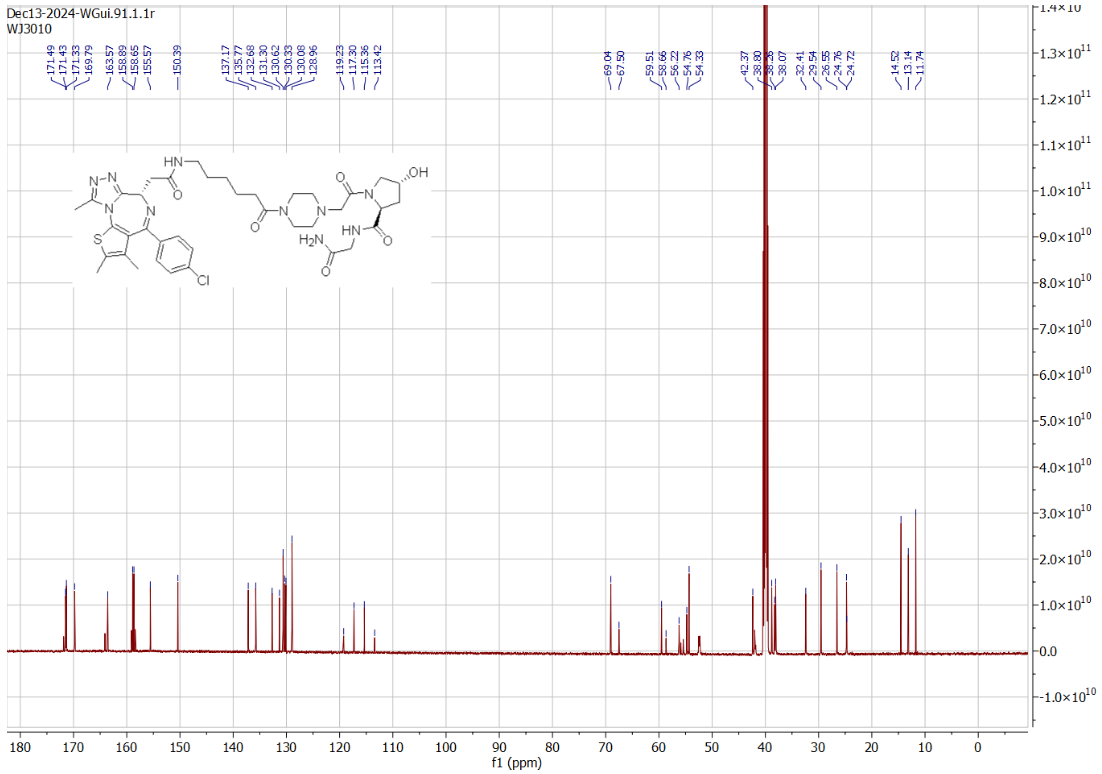

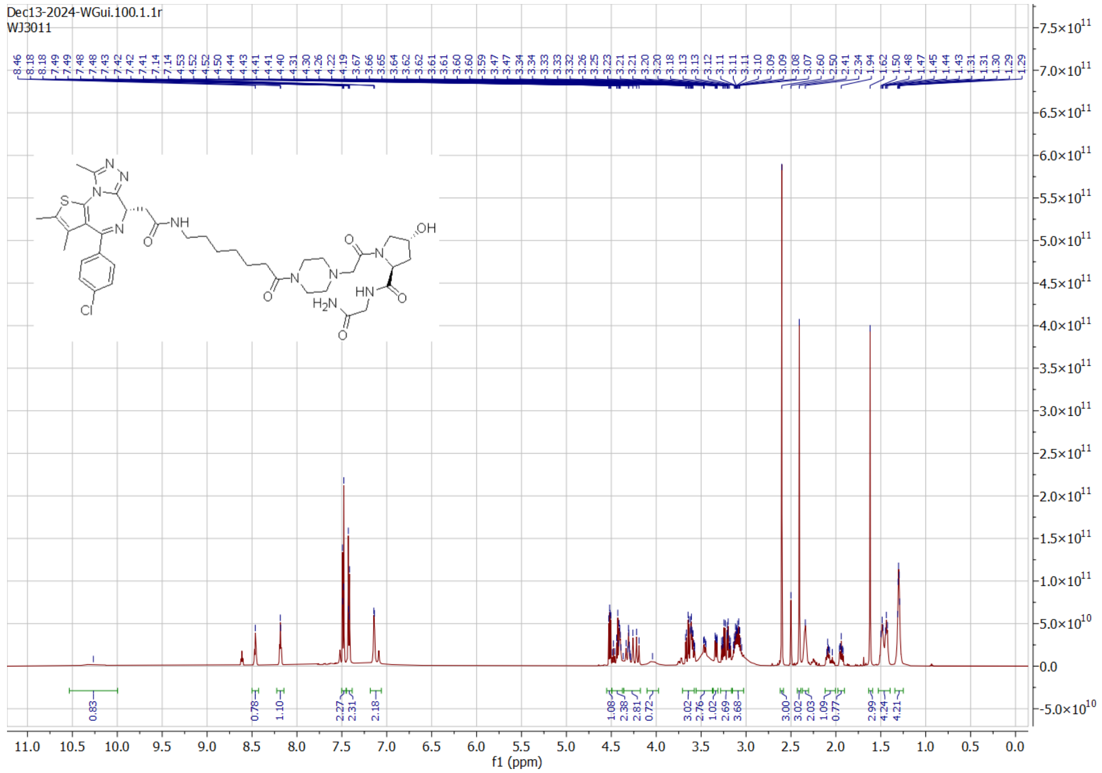

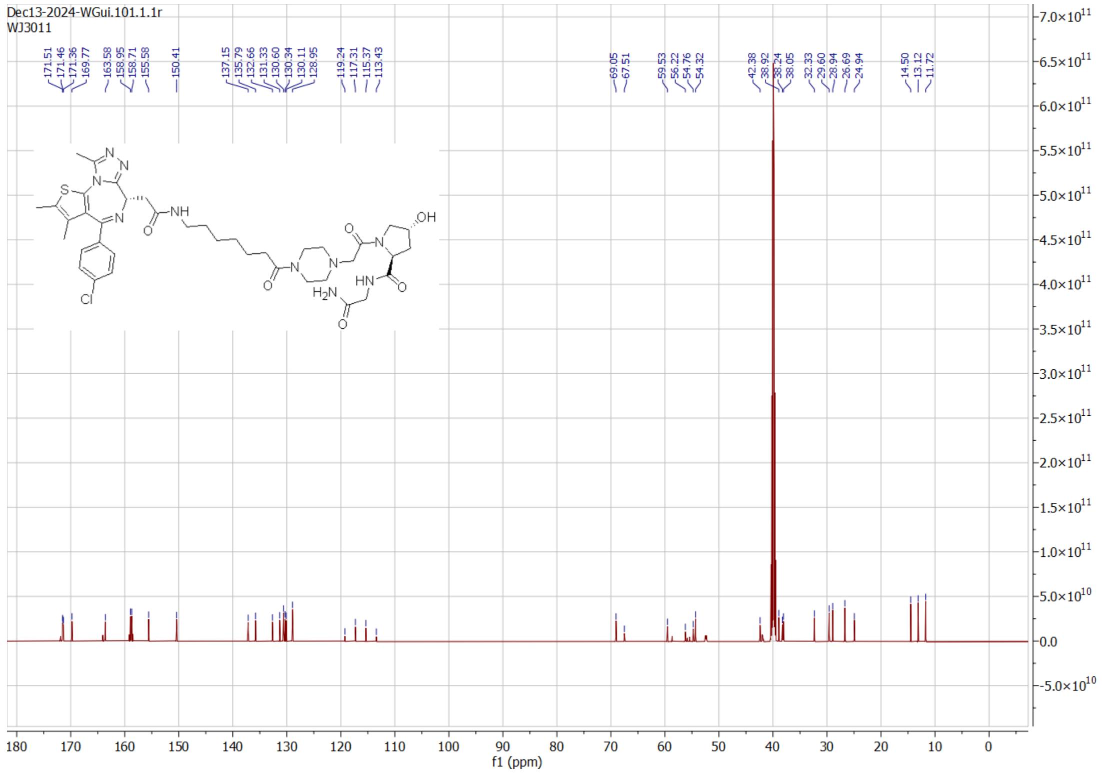

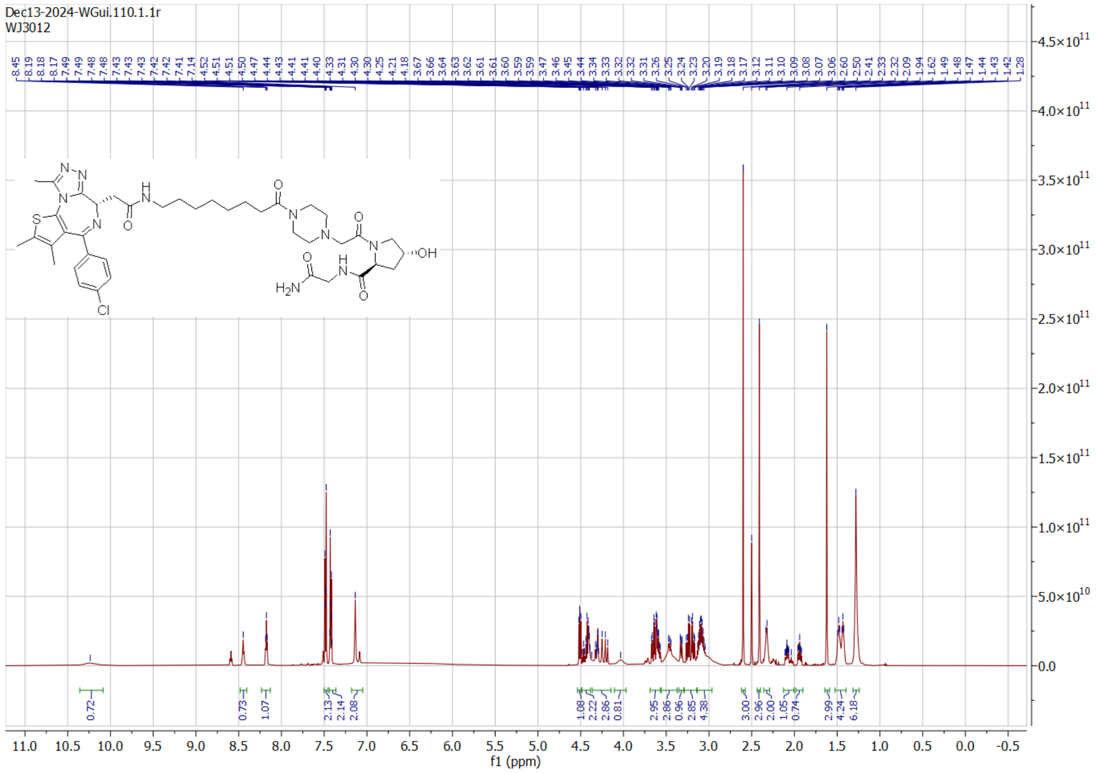
Dec13-2024-WGui.110.1.1r
WJ3012
f1 (ppm)

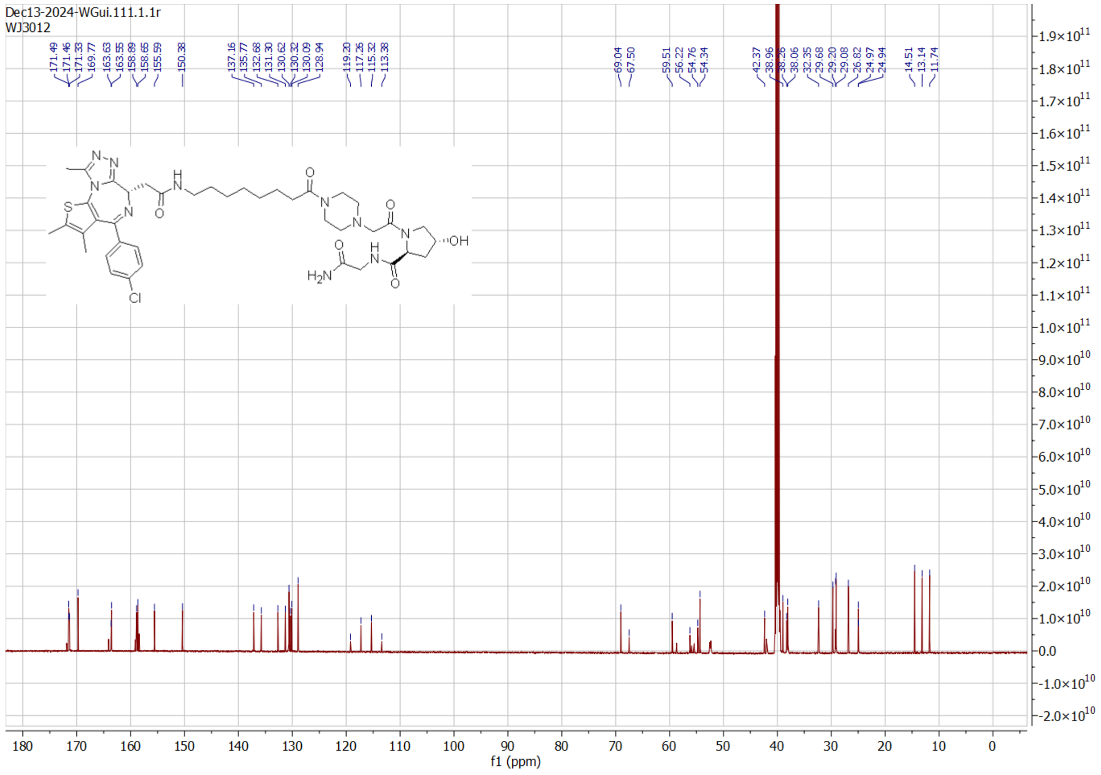
Dec13-2024-WGui.111.1.1r
WJ3012
f1 (ppm)

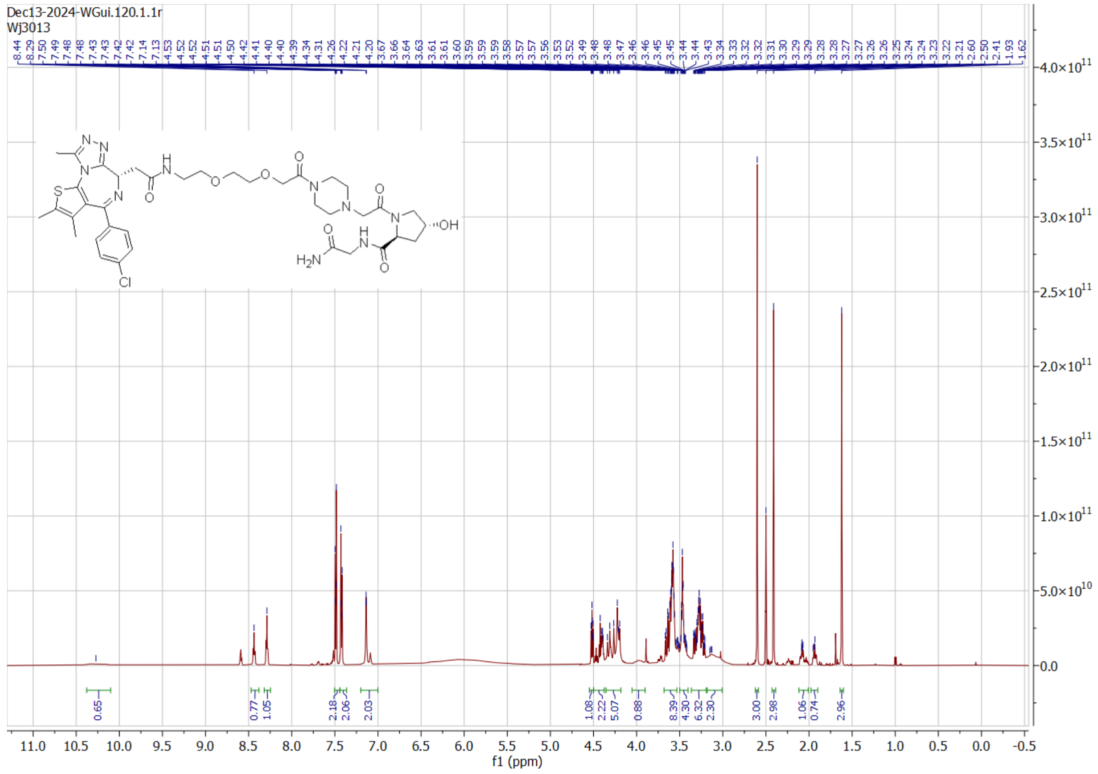
Dec13-2024-WGui.120.1.1r
Wj3013
f1 (ppm)
Cl
H2N
OH

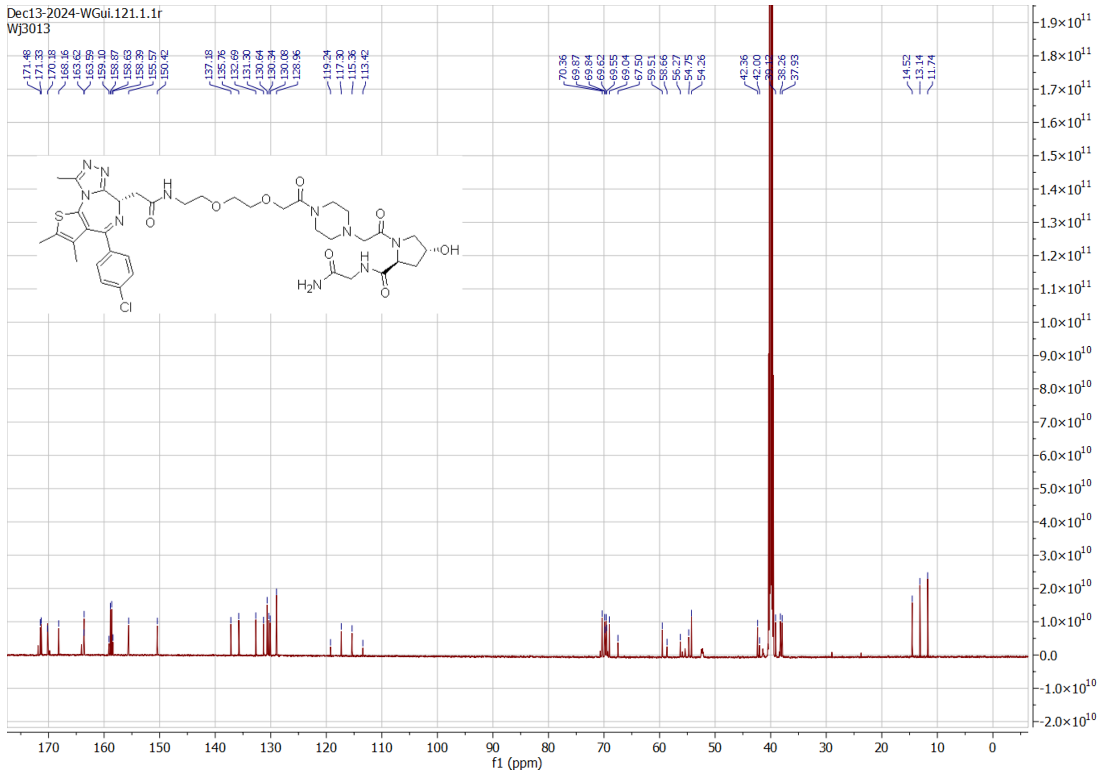
Dec13-2024-WGui.121.1.1r
Wj3013
f1 (ppm)
Cl
H2N
OH

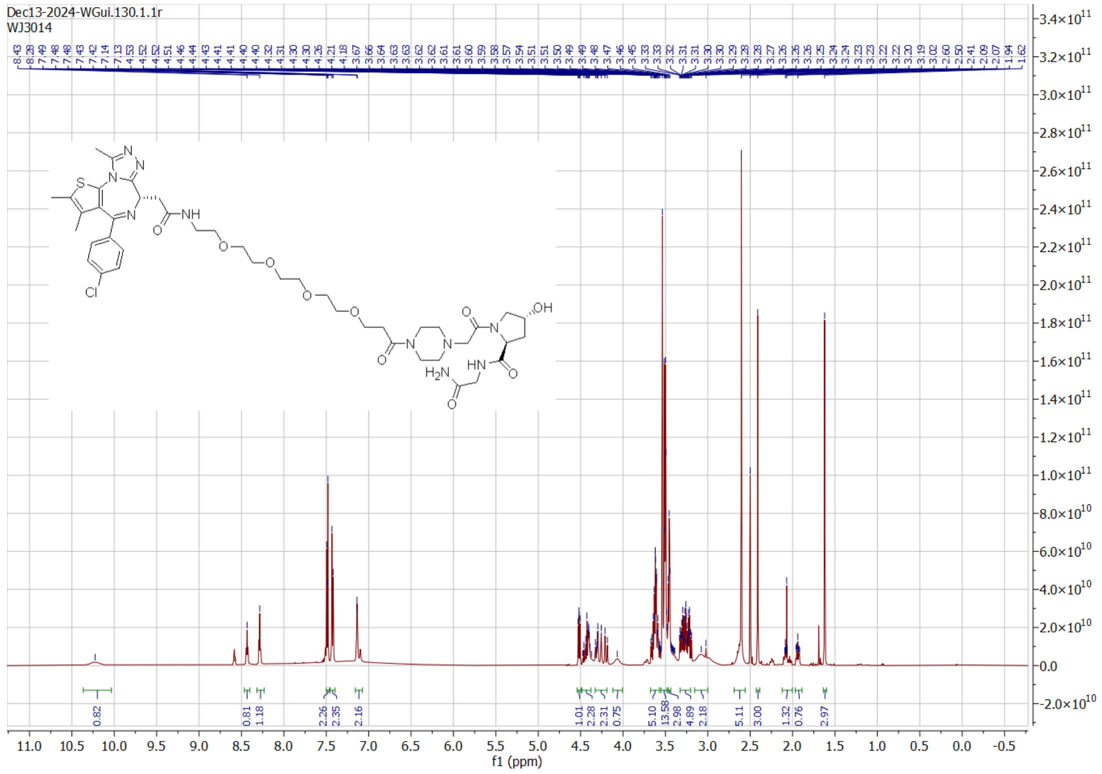
Dec13-2024-WGui.130.1.1r
WJ3014
f1 (ppm)

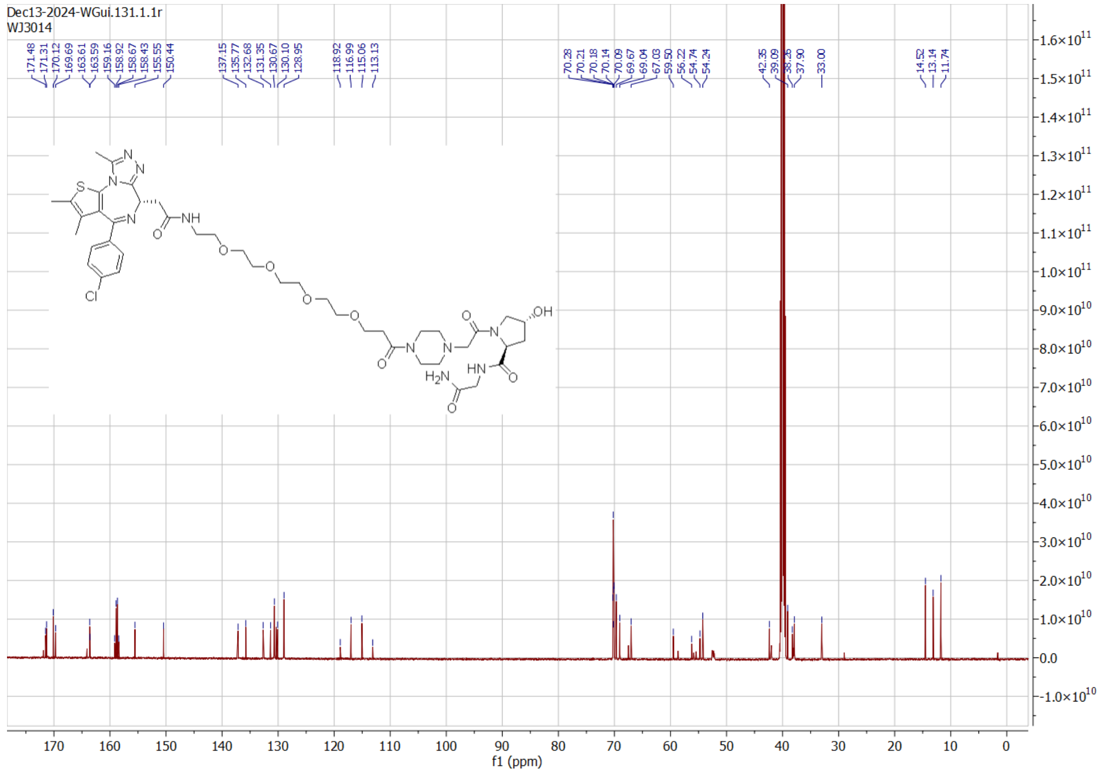
Dec13-2024-WGui.131.1.1r
WJ3014
f1 (ppm)

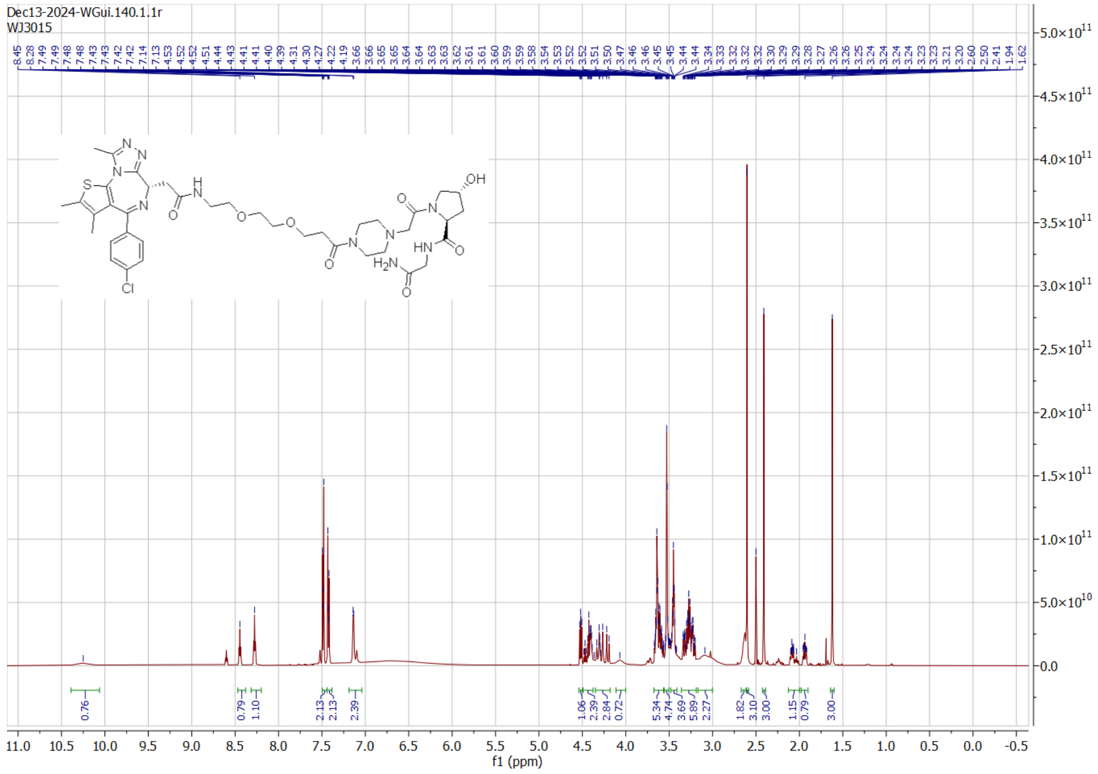
Dec13-2024-WGui.140.1.1r
WJ3015
f1 (ppm)

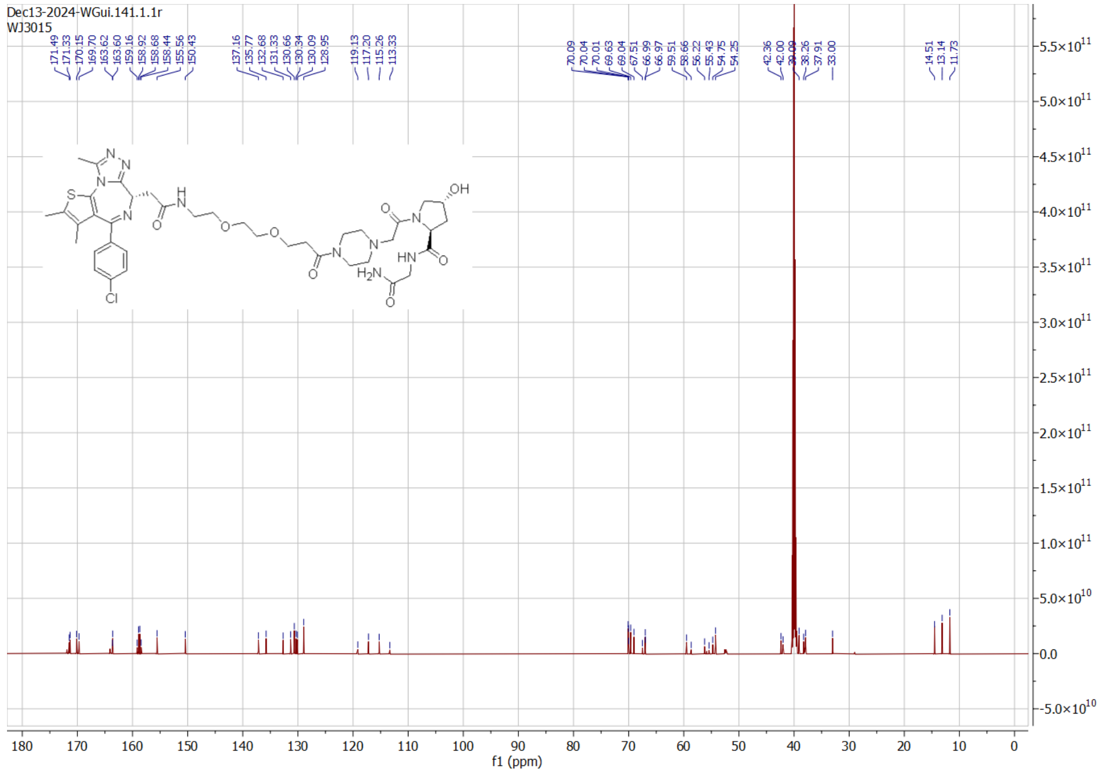
Dec13-2024-WGui.141.1.1r
WJ3015
f1 (ppm)
